# A precise atlas of the human subcortex

**DOI:** 10.64898/2026.02.13.705755

**Authors:** Helen Friedrich, Ilkem Aysu Sahin, Eduardo J. L. Alho, Nanditha Rajamani, Vanessa Milanese Holanda, Simon Oxenford, Malte Brammerloh, Maike Mustin, Patricia Zvarova, Lara Krüger, Richard Demann, Lukas L. Goede, Garance M. Meyer, Evgeniya Kirilina, Cordula Matthies, Jens Volkmann, Julianna Pijar, Shiro Horisawa, Calvin Howard, Arun Garimella, Ningfei Li, Michael D. Fox, Maximilian U. Friedrich, Anja K. E. Horn, Brian L. Edlow, Clemens Neudorfer, Andreas Horn

**Author notes:** Corresponding authors: Further information and request for resources should be directed to and will be fulfilled by the lead contacts, Helen Friedrich and Andreas Horn.

## Abstract

Clinical interventions and neuroimaging in the subcortex require anatomical definitions that exceed the resolution and anatomical detail of currently available deformable brain atlases. Here, we introduce a high-resolution human brain atlas comprising 85 bilateral and 10 midline manually segmented grey and white matter structures, corresponding to 180 anatomical segmentations in total, as well as 77 white matter tracts, compiled from a multitude of resources including ex-vivo MRI, histology, fibre dissections, and neuroanatomy textbooks. The atlas is defined at an isotropic resolution of 100 µm and can be precisely deformed to individual subject brain anatomy. By providing precise definitions of both grey and white matter structures within and around the basal ganglia, thalamus, subthalamus, midbrain and cerebellum, the atlas provides a foundational resource for stereotactic surgery and subcortical brain imaging research, as well as for development of next-generation neuromodulation strategies.

## Introduction

The subcortical anatomy of the human brain is complex and highly fragmented with numerous small structures that border and interact with one another. As outlined by ^1^, the Federative Community on Anatomical Terminology lists 455 structures in the subcortex, of which only 7% have been mapped in standard neuroimaging atlases, motivating the authors to speak of the subcortex as the ‘terra incognita’^1^ of the human brain. This is a critical gap, as subcortical structures play an increasing role in anatomically targeted medical interventions such as deep brain stimulation (DBS), gene therapies, and ablative neurosurgery, including modern incisionless interventions such as MR guided focused ultrasound or Gamma Knife surgery. At the same time, recent advances in imaging and atlas-informed DBS programming increase the reliance on standardized anatomical reference frameworks. In fact, recently, the first FDA approved software for image guided DBS programming has entered clinical practice. With over 300,000 patients successfully treated with DBS for various brain disorders such as Parkinson’s disease, essential tremor, dystonia, obsessive compulsive disorder, epilepsy, chronic pain, depression or Alzheimer’s disease^2^, the need for anatomically precise and scalable atlas resources has become increasingly pressing. Beyond clinical applications, the field of brain imaging is set to explore the subcortex^1^, with the advent of areas such as precision functional mapping^3^ or brainstem structural connectivity mapping^4^. Hence, precisely mapping the subcortex has never been more important and is now within reach.

Once subcortical regions are mapped in normative resources, the resulting brain atlases can be co-registered to individual patient anatomy to delineate structures in individuals, such as for functional brain mapping experiments or patients undergoing surgical interventions. Alternative strategies that directly aim at segmenting brain structures^5–7^ require similar normative atlas information ^5,6^. This remains true for modern artificial intelligence (AI) based approaches, in which normative information is stored in the form of neural networks^5,8^ that were trained to output brain segmentation labels. Accordingly, for any type of currently available approach that aims at segmenting subcortical regions in individual brain scans, detailed normative atlas information is required. The process of mapping these atlas delineations onto individual brains remains challenging but has become increasingly precise over recent years, regardless of whether using conventional^9,10^ or AI-based methods^11^.

A large number of brain atlases have been published and many of them have been applied in human brain mapping or functional neurosurgery^12–15^. In most of them, however, subcortical definitions are sparse and either focus on key surgical target structures themselves (but sparing neighbouring definitions) or map the subcortex in rather coarse fashion^1^. As a result, there is less potential for serendipitous insights^16^ into the effects of lesions or brain stimulation in these regions, possibly leaving clinical relevance of some regions undiscovered, since they are simply not defined in the atlases we use today^16–22^. Beyond limiting anatomical interpretation, insufficient visualization of subcortical structures may also constrain DBS targeting. Consistent with this idea, ^23^ reported that 7T-versus 3T-based direct targeting for essential tremor yielded greater tremor improvement, lower stimulation currents, and less variability in final electrode location, likely because there was more anatomical detail to be seen in the targeted areas^23^.

Current brain atlases can broadly be grouped into two categories: atlases directly derived from histology^11,14,24–27^ and atlases that were anchored on MRI scans^12,13^ – which may often include histological information that was registered to the MRI^15^. Each of the two approaches has advantages and limitations. While histology represents the gold-standard for anatomical definitions^11^, its use in 3D atlasing includes challenges when i) registering slices to tissue-blocks and when ii) making them deformable to individual patients^28^. Both processes are non-trivial and face unsolved technical challenges such as slice-wise distortions and cumulative curvature arising during 3D reconstruction of serial histological sections (the z-effect, or ‘banana-problem’)^29,30^. In addition, said processes require to either register histology to the same brain scanned using MRI^28^ or the use of ‘pseudo-MRI’ volumes generated directly from histological segmentations^15^. When instead using MRI derived anatomical definitions, registration issues are less reliant on assumptions. However, given that even dedicated MRI scans used in atlasing have limited contrast and resolutions of around 0.5 –1 mm^15^ at best, small and often poorly defined subcortical structures are easily missed on MRI.

In 2019, a team at Massachusetts General Hospital introduced an MRI template of unprecedented quality, which did not require 2D to 3D registration but still featured the anatomical detail needed to constitute the backbone for a high-resolution atlas of the subcortex^31^. This ultra-high-resolution MRI scan was acquired by scanning a postmortem specimen of a 58-year-old woman over an entire weekend, with over 100 hours of scan time and use of a custom-built coil at 7 Tesla. The result was the most detailed whole-brain MR dataset of the human brain available to date, featuring an isotropic resolution of 100 µm, i.e. ∼1,000 times higher than conventional MRIs. The template yields exceptional tissue contrast, preserving the brain’s intact three-dimensional geometry without substantial distortions and without requiring 2D slice registrations plagued by the unresolved technical challenges outlined above. A version of the 7T template registered to standard stereotactic space has since become widely adopted, serving as an anatomical reference in over 270 studies^9,17,32–37^. Moreover, histological sections as well as diffusion-weighted MRI data from the same brain are available, which make it possible to refine structural definitions further.

Here, we leveraged this unique specimen to create an ultra-high resolution brain atlas, circumventing typical registration problems of histological data while preserving the highest possible level of anatomical detail. This multimodal dataset (diffusion MRI, ex-vivo structural MRI, histological stacks of the same brain) served as a backbone to anchor our atlas. However, we further augmented its definition by the use of an extensive library of 48 additional resources that included gross-dissectional anatomical specimens as well as histological stacks and stainings from other sources, libraries of structural and diffusion MRI, and neuroanatomical textbooks. Consequently, each definition of a subcortical nucleus or white matter tract in the resulting atlas reflects a synthesis of multiple anatomical sources critically evaluated by a multidisciplinary team of neuroanatomists, neurologists, neurosurgeons, and MR physicists.

As a result, the *FOCUS* atlas is intended to be a **f**lexible **o**pen atlas for **c**linical **u**se in the **s**ubcortex. In its current first version, we focus on intervention zones of highest clinical interest by providing refined, anatomically differentiated definitions of well-established structures as well as neighbouring structures, internal subdivisions, and previously underrepresented elements within these zones. We see the atlas as an open data platform which will be iteratively developed further, creating opportunities for other laboratories to contribute refinements or additional segmentations.

## Results

The present study introduces a high-resolution subcortical brain atlas of grey and white matter, as well as a precise and a dedicated workflow to register the atlas to individual brain scans. To create these resources, we developed and extended two open-source software tools. First, it was required to extend functionality of the WarpDrive toolbox^38^ which is used to manually curate nonlinear warps between anatomical spaces, specifically to refine computer generated solutions that are unaligned in parts of the image but can be highly precise in areas of focus. In other words, the tool enables manual alterations to automated registration fields across spaces, which have general anatomical accuracy but are naturally imperfect in some parts of the image. Here, WarpDrive was extended to enable definition of anchor structures across spaces, to invert registration views, to include a ‘shrink’ and ‘expand’ mode, and to include visualization of manual segmentations in both spaces (see *Methods*). Second, to manually define fibre bundles in 3D space, the CurveToBundle software tool was developed, which facilitates anatomically guided tract reconstructions. We made both tools openly available as 3D Slicer modules (https://github.com/netstim/SlicerNetstim).

Using the ex vivo 100 µm MRI dataset provided by Edlow et al. ^39^ as a primary reference, we manually segmented 85 distinct grey and white matter structures per hemisphere and 10 midline structures, resulting in a total of 180 individually delineated regions. These structures are complemented by 77 additional white matter bundles. Our rationale of choosing which structures to include followed a mix of factors which were mainly driven by clinical relevance, target structures of functional neurosurgery themselves as well as their neighbouring structures, and, finally, visibility of structures on the ex-vivo MRI. Namely, particular emphasis was placed on zones of clinical interventions (see *Figure 1*), with special attention to the detailed definition of the mesencephalic-diencephalic transition zone, as a prime area for deep brain stimulation surgery. This region includes multiple DBS targets, such as the subthalamic nucleus, the fields of Forel, the posterior thalamic area, the zona incerta, the ventral intermediate and centromedian/parafascicular nuclei of the thalamus and the internal pallidum (*Figure 1c, Supplementary Materials*).

**Figure 1.**
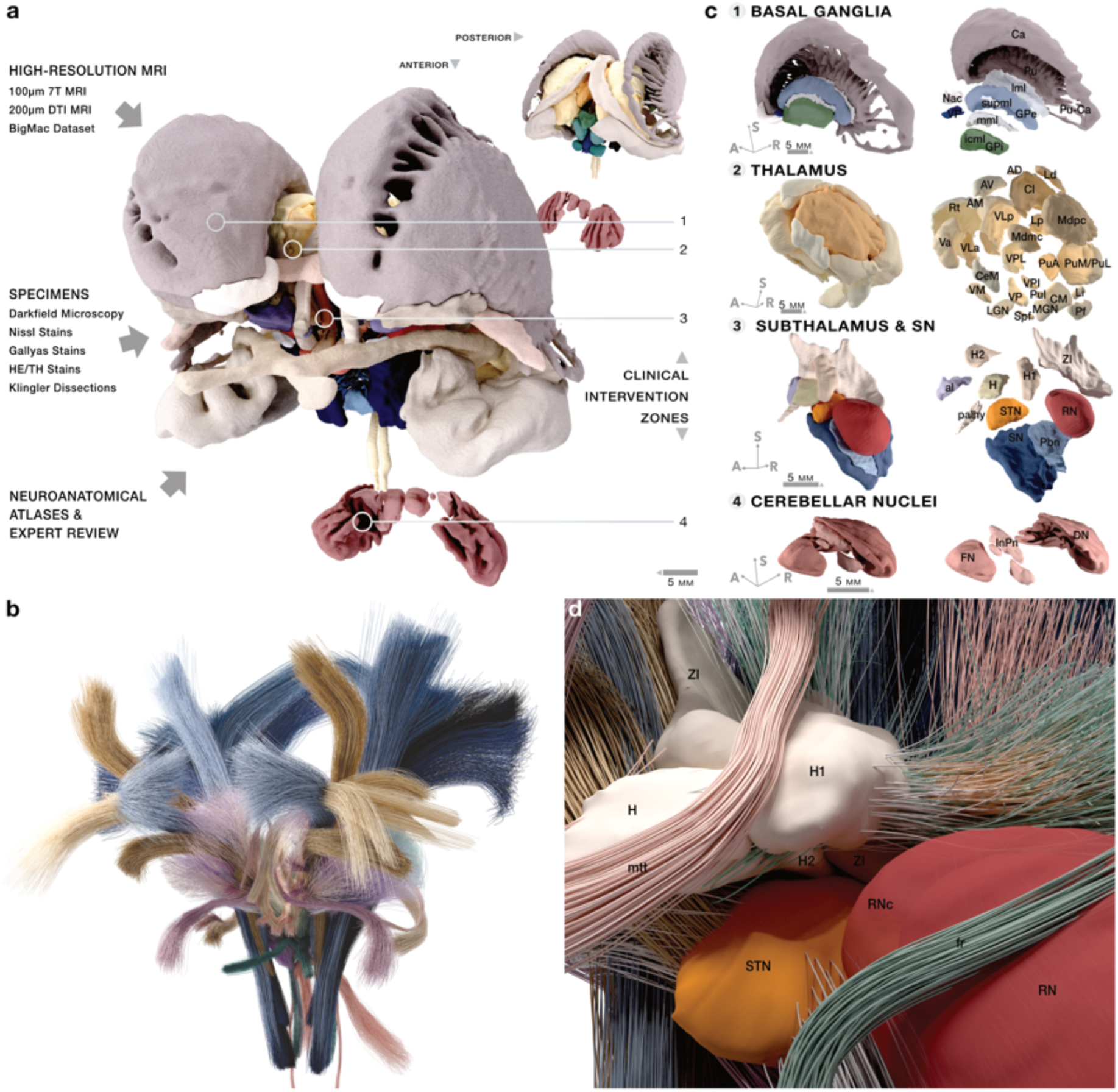
FOCUS atlas: Overview. **a & b**) The atlas comprises 180 individual subcortical grey and white matter segmentations, including bilateral structures and midline structures, as well as 77 fibre tracts. An exemplary subset is shown here. As a primary resource, an ex-vivo MRI template acquired at 100 µm resolution was used, which provided rich anatomical detail (see *Supplementary Table 1)*. However, particularly in regions with subtle contrast boundaries, the rich detail of anatomical structures required extensive refinement based on a total of 48 additional resources (see *Supplementary Materials*). **c)** Particular emphasis was placed on the following clinically relevant intervention zones in which anatomical detail is sparse in most presently available deformable atlas resources. **1** Basal ganglia: Fine-grained segmentation allowed for a detailed reconstruction of internal laminae and bridging cell clusters between the putamen and caudate nucleus. For example, the supplementary medullary lamina which is rarely captured on MRI was clearly identifiable (see *Supplementary Materials)*. **2** Thalamus and mesencephalic transition zone: The intricate architecture of the thalamus and the mesencephalic-diencephalic transition zone was delineated and mapped to standard stereotactic space. Clinically relevant target areas such as the **3** Fields of Forel and associated fibre tracts, such as the ansa lenticularis including axon collaterals to intralaminary nuclei of the thalamus could be traced throughout their course (see *Supplementary Materials*). Remarkably, even fine-caliber bundles such as the pallidohypothalamic tract with less than 2 mm in diameter or the fasciculus retroflexus (see *Figure 6*) could be successfully reconstructed. Subfigure **4** shows the cerebellar nuclei (Details see *Supplementary Materials*). **d)** Magnified medial-to-lateral sections showing the interplay between grey matter nuclei and fibres traversing them. As opposed to diffusion-tractography based results, fibres traced using the present method will not stop at grey-/white matter boundaries due to a drop of diffusion signal. This leads to more anatomically plausible tracts that fan out within their grey matter terminal zones, which can be critical, e.g. when modelling deep brain stimulation effects from electrodes placed in grey matter nuclei onto the axons that traverse these nuclei. *Abbreviations*: al, ansa lenticularis; AM, anteromedial nucleus; AV, anteroventral nucleus; Ca, caudate nucleus; CeM, central medial nucleus; CL, central lateral nucleus; CM, centromedian nucleus; DN, dentate nucleus; FN, fastigial nucleus; fr, fasciculus retroflexus; GPe, external segment of the globus pallidus; GPi, internal segment of the globus pallidus; H, field of Forel H; H1, field of Forel H1; H2, field of Forel H2; icml, incomplete medullary lamina; InPn, interposed nucleus; lml, lateral medullary lamina; LD, lateral dorsal nucleus; LGN, lateral geniculate nucleus; Li, limitans nucleus; MDmc, mediodorsal nucleus, magnocellular division; MDpc, mediodorsal nucleus, parvocellular division; MGN, medial geniculate nucleus; mml, medial medullary lamina; mtt, mammillothalamic tract; Nac, nucleus accumbens; palhy, pallidohypothalamic tract; Pbn, parabrachial pigmented nucleus; Pf, parafascicular nucleus; PuA, anterior pulvinar; Pu-Ca, putamen–caudate bridges; PuI, inferior pulvinar; PuM/PuL, medial and lateral pulvinar nuclei; RN, red nucleus; RNc, capsule of the red nucleus; Rt, reticular nucleus; sPf, subparafascicular nucleus; SN, substantia nigra; STN, subthalamic nucleus; supml, supplementary medullary lamina; Va, ventral anterior nucleus; VLa, ventral lateral anterior nucleus; VLp, ventral lateral posterior nucleus; VM, ventral medial nucleus; VP, ventral pallidum; VPI, ventral posterior inferior nucleus; VPL, ventral posterior lateral nucleus; ZI, zona incerta.

Each segmentation was carried out at 100 µm resolution and was guided by a carefully curated, multimodal review process. To define structural boundaries reliably, we drew on a broad range of anatomical references including modern and historical atlases, textbook references, cadaveric fibre dissections, and original histological material (see *Figure 1* and *Supplementary Materials* for an overview).

Multiple neuroanatomists, neurologists, neurosurgeons and MR physicists (H.F., A.K.E.H, E.J.L.A., V.M.H., M.B., E.K., C.N., M.U.F., S.H., B.L.E.) guided the process of selecting specific staining and visualization techniques (e.g. Gallyas silver stains, hematoxylin-eosin (H.E.)/ Nissl stains, dark-field microscopy) to resolve ambiguities in and around regions that exhibited complex MRI signal properties (see *Supplementary Materials*).

A complete list of segmented structures is provided in *Supplementary Table 1*. *Supplementary Materials Section 1* and the detailed *Segmentation Protocol* in the *data repository* document a representative subset of cross-references between individual segmentations and the anatomical sources used to inform them. The result of this process is the FOCUS atlas itself, which was first defined in the native space of the 100 µm ex vivo template^39^. To maximize impact of the atlas and allow for it to be deformable into patient-/subject-specific MRI space, we next created a registration between this native space and a high-resolution MNI template (ICBM 2009b NLIN Asym). Of note, this coregistration between the 100 µm native space and this particular MNI template can be deemed just as critical as the atlas definition itself, since slight inaccuracies of registrations may propagate into erroneous segmentations of patient space.

Hence this transformation was iteratively refined using a multi-stage normalization pipeline, which combined multispectral nonlinear registration with a high-resolution refinement protocol (*Figure 2*). A total of 62 anchor segmentations (31 per hemisphere) served as fiducial regions, guiding over 37 iterative refinement steps (see *Figure 2* and for more details *Supplementary Materials, Section 2*) with a specific focus on subcortical structures that play a prominent role in current clinical practice (*Figure 1c*). To further optimize alignment in selected areas, we used the WarpDrive toolbox to carry out localized adjustments to the deformation field (*Figure 2c*).

**Figure 2.**
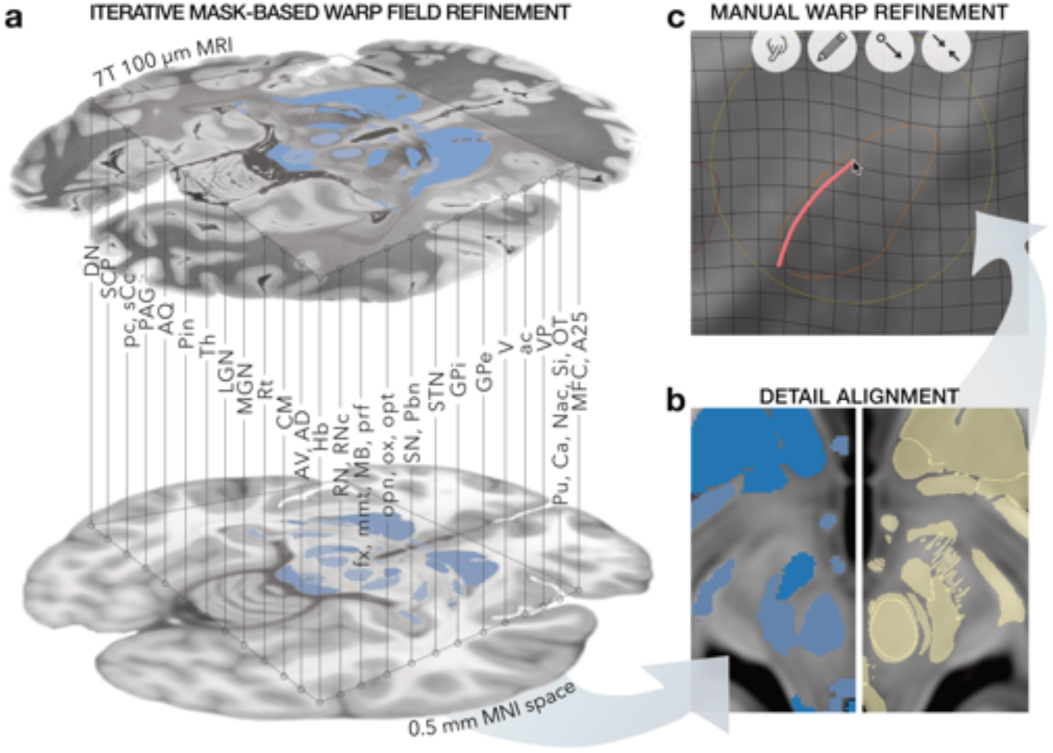
Transfer of High-Resolution FOCUS structures into Standard Stereotactic Space. To maximize precision of high-resolution segmentations and to make them deformable into individualized patient space, a precise registration to standard stereotactic space was required. **a)** First, anchor structures (blue) that were well distinguishable in both the 100 µm and T1, T2 or PD weighted templates of the MNI space were carefully segmented in both spaces. Structures were iteratively included into further refined versions of the warp in a 37-step iterative co-registration process (see *Supplementary Materials*). This process generated an accurate warp field, allowing for the **b)** precise topographic alignment of high-resolution structures even without direct anchors, which instead relied on their reference to neighbouring anchor elements (in yellow; see *detailed protocol at the referenced data repository*). **c)** When required, local adjustments were finally applied using the Warp-Drive software (see e.g. **c** for STN boundary adjustments).

The resulting final nonlinear (and diffeomorphic) transformation made it possible to transport the segmentations from the 100 µm space into MNI space, leading to 100 µm resolution definitions of all structures in MNI space (*Figure 3*). These were visually validated across all orientations (*see Supplementary Materials*). Operating at a mesoscale resolution, the atlas hence combines high anatomical detail with full compatibility to a standardized coordinate space, making it directly deformable into individual patient anatomy while preserving high spatial resolution.

**Figure 3.**
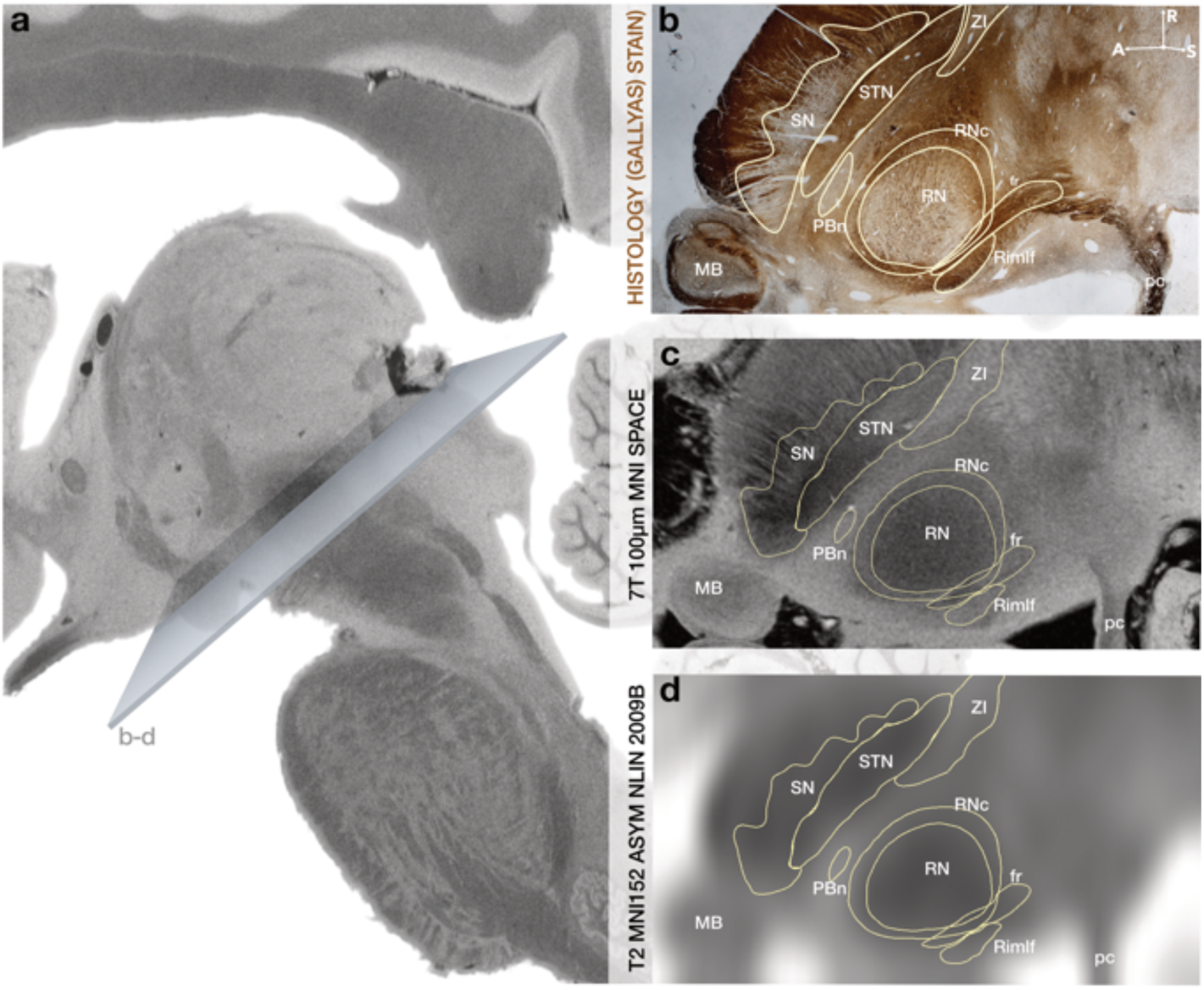
Validation of topographic accuracy for coregistered atlas structures in standard space. To confirm the accuracy of the coregistered atlas definitions, hence the quality of the warp field, we compared them with histological specimens and reference atlases. For this, we matched the **a)** same cross-sectional planes with the **b)** histological references. Refinements of the warp field were made iteratively (see *Supplementary Materials Section 2* and *detailed protocol in the referenced data repository*) until additional iterations no longer yielded meaningful improvement. (**c)** & **d)**).

As a natural secondary outcome of the refined registration between 100 µm ex-vivo and MNI spaces, we provide an updated MNI version of the template itself. While an MNI version of the dataset had been originally released by the original authors^39^ our updated version offers a more accurate fit in subcortical areas, especially in the clinical intervention zones of high interest, such as the thalamus, the subthalamus, and the midbrain (see *Supplementary Materials* for a comparison between original and refined templates).

To facilitate white matter-based analyses, we manually reconstructed a set of unilateral 77 fibre pathways to complement the atlas. These were informed by the previously generated segmentations (*Figures 1b-d, 4*). Fibre tracts were directly constructed in MNI space, since their definition did not require the 100 µm space itself but instead landmark guidance provided by i) structures visible on the MNI templates, ii) mapped FOCUS atlas structures and iii) diffusion MRI reference datasets, gross-dissection, staining results, and text-book results (*Figure 5*). Incorporating the 100 µm segmentations as anatomical guidance priors provided added value: we observed more anatomically constrained tract trajectories, particularly in regions where key structures were only a few millimetres in size and their exact anatomical course could have been easily missed without precise anchor structures (*Figure 5g-j*). Unlike diffusion-based tractography, which can be limited by crossing fibres and partial volume effects, our approach was inspired by the framework of Petersen et al.^40^, which we describe in *Figure 4* (and refer to as ‘tract architectonics’ in the *Methods*). In brief, the framework by Petersen et al. reconstructs fibre pathways manually from convergent anatomical priors and landmark constraints, rather than estimating them directly from diffusion MRI alone. Example structures include the fasciculus retroflexus, the pallidohypothalamic tract and the oculomotor nerve, whose anatomical trajectories would have been difficult if impossible to trace using a conventional diffusion MRI-based tractography approach. Instead, using our stepwise segmentation approach, we were able to model such delicate tracts with high spatial and anatomical accuracy (see *Figures 5-6*).

**Figure 4.**
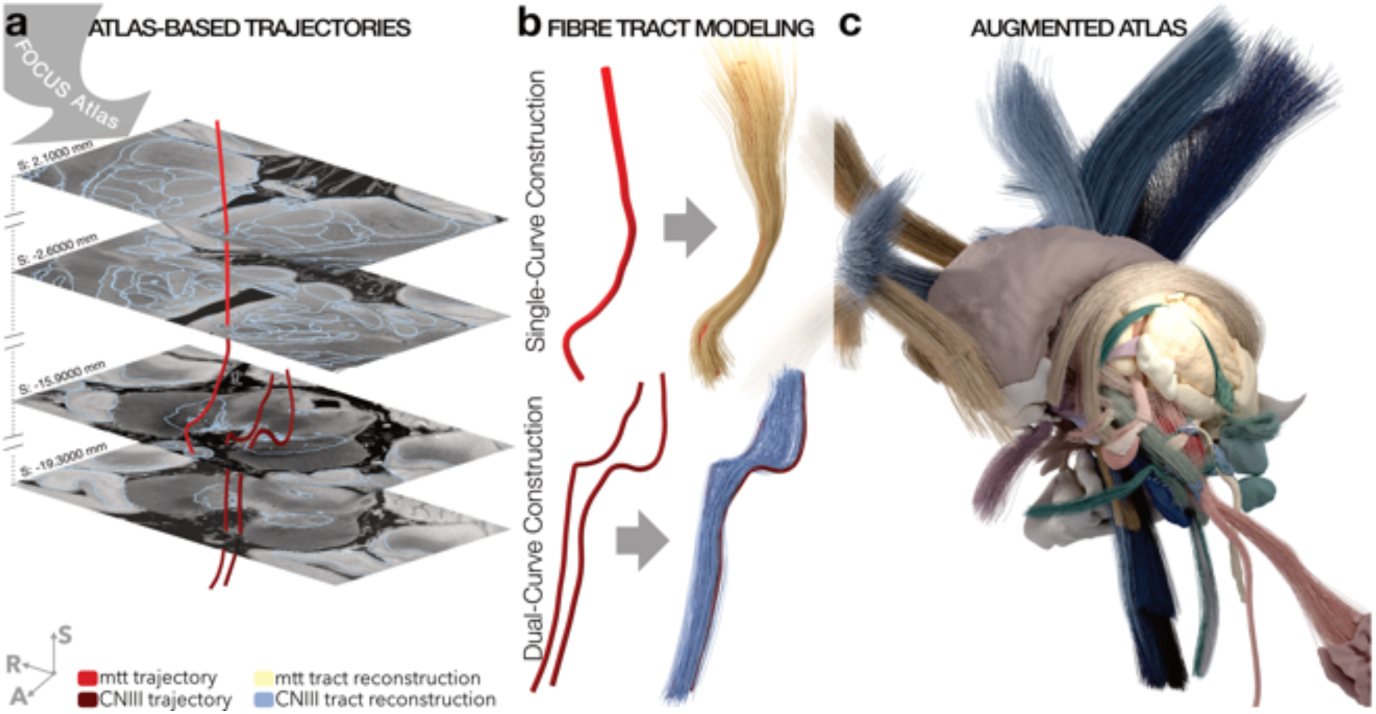
Manual curation of streamline bundles. **a)** Atlas structures as the ones provided by segmentations of the FOCUS atlas helped to trace streamline trajectories based on a variety of resources that included diffusion MRI data, histology, axonal tract tracing and staining results (see *Supplementary Materials*). **b)** The CurveToBundle tool samples uniformly distributed streamlines along a single (top example) or dual (bottom example) curve trajectory. At each point along the curve, a diameter for the distribution can be selected. Termination zones or zones of passage for the streamlines can be restricted by atlas structures. For instance, for tracts known to project to individual thalamic nuclei, the spread of their terminals was restricted by the voxelized definition of the respective nucleus. **c)** Resulting fibre atlas that augments voxelized segmentations.

**Figure 5.**
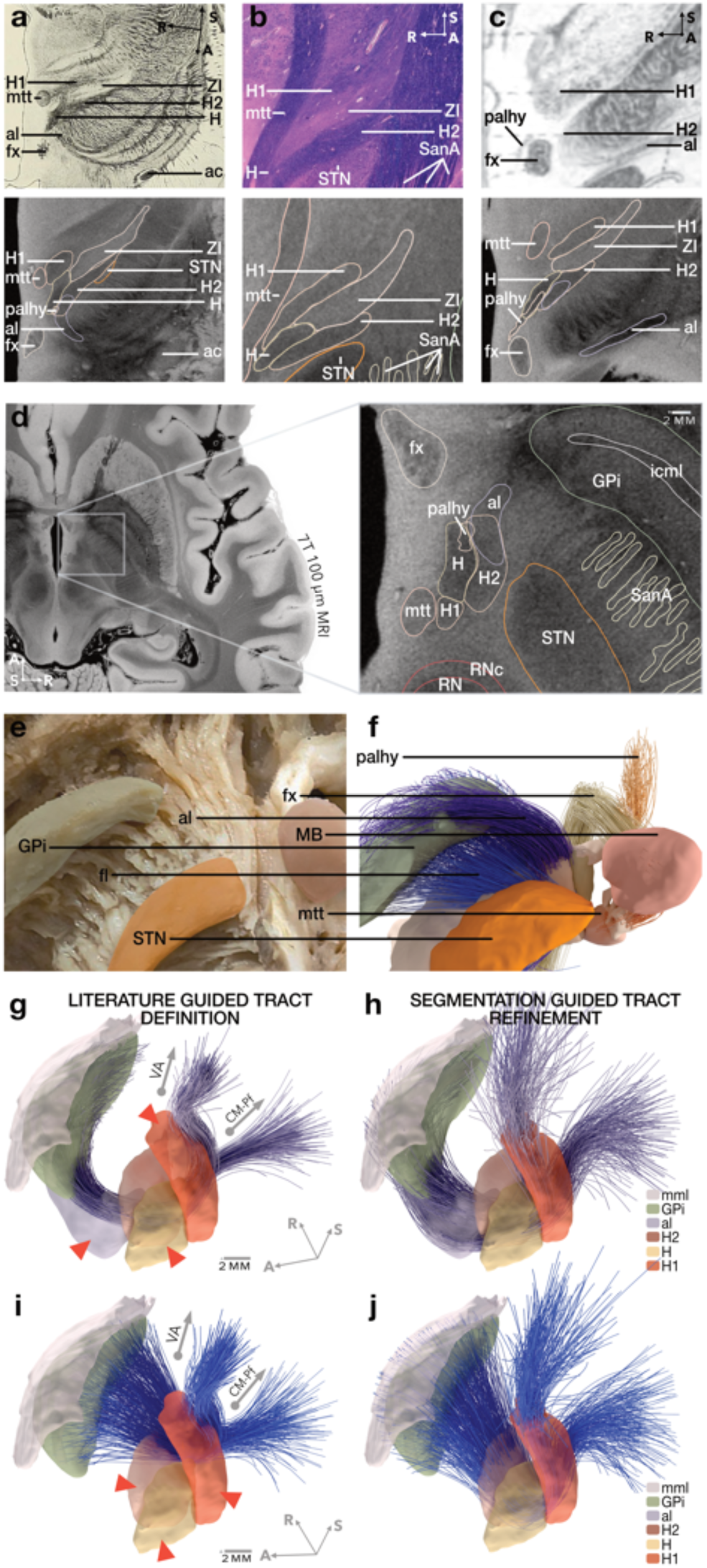
Anatomical workflow for defining subcortical structures: Example of Forel Fields. **a-c)** The ability to reslice high-resolution 3D MRI in any oblique direction was leveraged to directly align structural imaging with anatomical references (including historical depictions. Three key resources (top row), each emphasizing different aspects of the region’s anatomy, are given as illustrative examples. Resources were systematically compared with corresponding MRI slices and our manual segmentations (bottom row) that were sliced in a comparable angle (see *Supplementary Materials*). **a)** An oblique horizontal section from ^47^, clearly depicting the course of the ansa lenticularis and overlapping fibre components. **b)** A histological section by ^48^ providing cellular-level resolution, essential for delineating fine structures and internal boundaries such as the tegmental fields and the Sano A fields (which are not formally part of the Fields of Forel, but included here to illustrate the level of anatomical detail achievable through this method). **c)** The foundational first descriptions by Auguste Forel (1877,^49^). This integrative, multimodal strategy enabled cross-validation across modalities and led to a precise segmentation of the complex subthalamic region. Among the uniquely resolved features are the fusion of the ansa lenticularis with the fasciculus lenticularis (H2; see *Supplementary Figure 2*) and the convergence of the pallidohypothalamic tract within the Haubenfeld H. **d)** Final high-precision segmentation, which was then meticulously registered into standard space (see *Methods*). **e-f)** (ventral-posterior oblique view) Based on these segmentations, we reconstructed and refined fibre tracts including the ansa lenticularis (purple) and the fasciculus lenticularis (blue). Correspondence with white matter dissection specimens supports the anatomical validity of the reconstructions. The advantages of this segmentation-guided approach are further demonstrated in panels **g-j)**: To highlight the limitations of manual pathway definitions that are unconstrained by voxelized segmentations, we compared reconstructions of pallidothalamic fibres in MNI space (0.5 mm resolution) that had been created in earlier work^40,42^ and the ones that were refined by the high-resolution anatomical segmentations created here. **g)** The ansa lenticularis (al) as manually reconstructed using detailed literature and anatomical guidance^40,42^. Not guided by anatomical constraints during registration, the tract missed the natural trajectory of the al, H and H1 (red arrows). **h)** When guided by volumetric segmentations, fibres of the ansa lenticularis matched segmentations of H and H1. **i-j)** shows the same analog analysis for the fasciculus lenticularis. *Abbreviations*: al, ansa lenticularis; GPi, internal pallidum; H, field of Forel H; H1, field of Forel H1; H2, field of Forel H2; mtt, mammillothalamic tract. Permission to reproduce panels that were adapted from original work (as indicated above) are on file.

**Figure 6.**
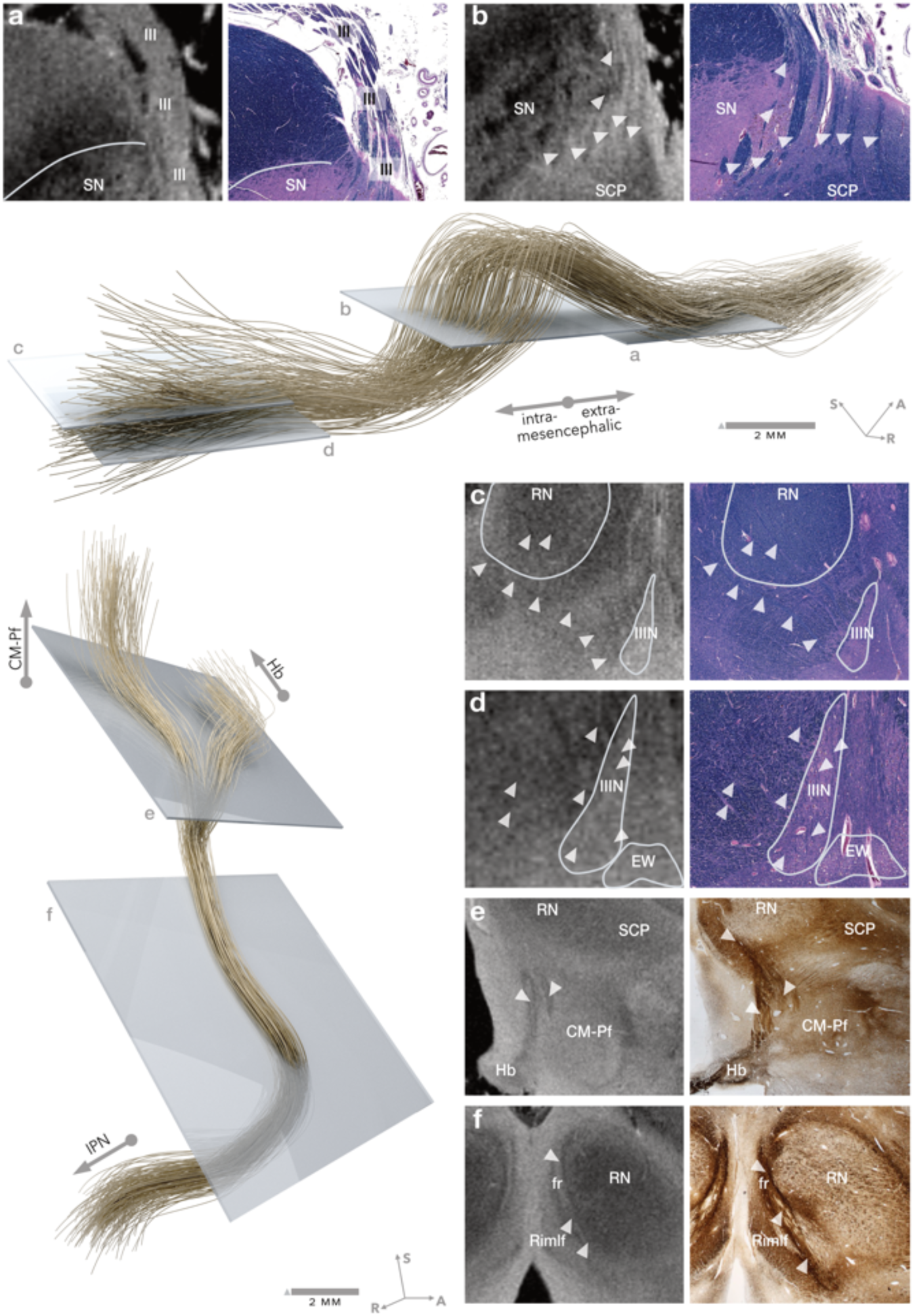
Manual reconstruction of complex fibre tracts based on multimodal guidance structures. The process of manual tract definitions that were constrained based on our high-resolution voxelized atlas definitions made it possible to reconstruct anatomically challenging pathways. Shown here are two representative examples that illustrate the level of anatomical detail achievable with this approach. The oculomotor fibres (top) could be traced from their exit the midbrain in form of a compact extramesencephalic bundle (panel **a**), their partial trajectory through the substantia nigra (**b**), while they course through the capsule of the red nucleus (**c**) and originate from the oculomotor nucleus (IIIN, **d**). The fasciculus retroflexus (syn.: habenular-interpeduncular tract or bundle of Meynert, bottom), the main efferent pathway projecting from the habenular nucleus (Hb) to the interpeduncular nucleus (IPN), was reconstructed along its characteristic steep descent and eponymous retroflexion. Importantly, we were able to resolve a distinct bifurcation of the tract (**e**) in the mesencephalic-diencephalic transition zone, well above the level of the red nucleus. One fibre component ascended toward the habenula, while a second branch coursed medially through the region of the parafascicular (Pf) and centromedian nucleus (CM) of the thalamus. While such a medial trajectory has occasionally been reported in histological material (e.g.^50^), it has not been explicitly described in prior literature, likely due to the limitations of two-dimensional sectioning and resulting anatomical uncertainty. Here, the 3D nature of our MRI-based reconstruction allowed this complex configuration to be visualized with high clarity. Panel **f** shows its trajectory medially to the red nucleus.

To illustrate the translational utility of this anatomical framework, we next carried out two proof-of-concept analyses that applied the FOCUS atlas to a clinical context. First, we related it to three ablative lesion cases that targeted the pallidothalamic tracts in dystonia. Second, we repeated a published analysis that identified tracts associated with optimal motor response in a large multicentre cohort following STN-DBS in Parkinson’s disease. Instead of the connectome used in the original report, tract definitions from the FOCUS atlas were used to define candidate tracts. These analyses are intended as illustrative applications of the atlas rather than as formal clinical validation studies.

In the first analysis, lesions carried out by stereotactic radiofrequency surgery in three patients with dystonia were overlaid with structures defined by the FOCUS atlas. Lesions targeted field H1 of Forel. Cases 1–3 showed 89.8%, 66.7%, and 0% improvement in BFMDRS-M, respectively (*Figure 7b–d*). Matching clinical outcomes, the lesion that led to no improvement was predominantly located within the zona incerta (ZI; 61.1% of lesion volume), followed by field H2 (20.9%), with smaller involvement of the subthalamic nucleus (STN; 6.6%) and reticular nucleus territories (Rt; 0.4%). Instead, both lesions that led to positive clinical response had strongest overlaps with H1. The one leading to 66.7% improvement showed the greatest overlap with H1 (38.7%), followed by the ventromedial thalamic region (VM; 32.4%), ZI (9.8%), VLa (5.5%), and H (1.3%). The one that led to 89.8% response also most prominently overlapped with H1 (32.8%), followed by ZI (18.9%), H2 (6.2%), STN (4.5%), and H (3.7%).

**Figure 7.**
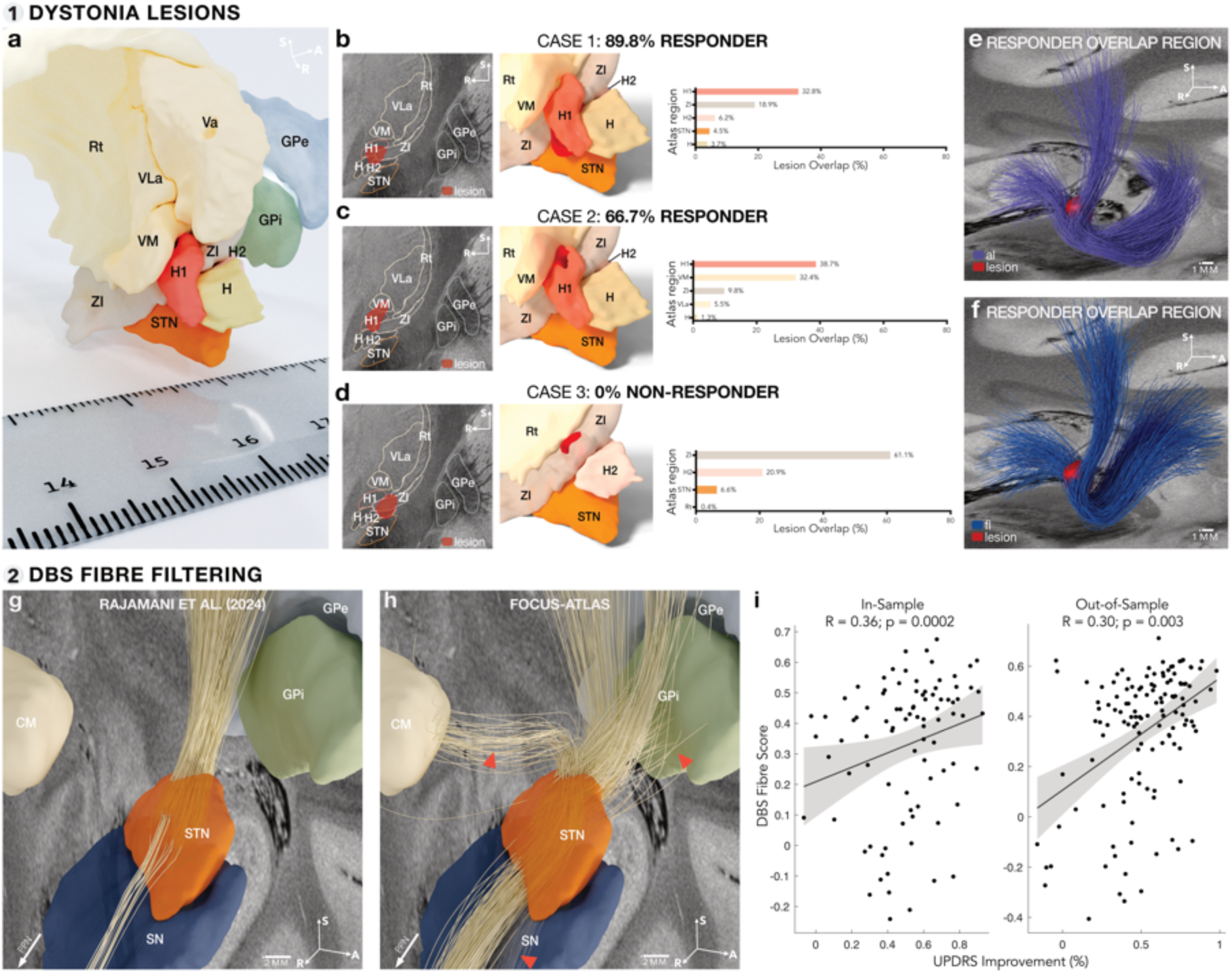
Illustrative applications of the FOCUS atlas. Lesion-based application in dystonia: **a**) Clinically targeted structures for lesion-based treatment of dystonia occupy only a few millimetres. To convey the small anatomical scale, the relevant high-resolution FOCUS atlas regions are shown alongside a standard ruler as a familiar object with an intuitive haptic sense of scale. **b)** Case 1, a responder after unilateral pallidothalamic tractotomy (PTT; 89.8% improvement in BFMDRS-M): lesion shown in red in 7 T MNI space in relation to the local subthalamic anatomy (left), 3D rendering of the lesion within the atlas context (middle), and atlas colocalization showing prominent H1 engagement (32.8% of lesion volume; right). **c)** Case 2, a responder (66.7% improvement): lesion shown in red in 7 T MNI space in relation to the local subthalamic anatomy (left), 3D rendering (middle), and atlas colocalization showing H1 engagement (38.7%; right). **d)** Case 3, a non-responder (0% improvement): lesion shown in red in 7 T MNI space in relation to the local subthalamic anatomy (left), 3D rendering (middle), and atlas colocalization showing predominant involvement of the zona incerta (ZI; 61.1%) and field H2 (20.9%), with no overlap with H1 (right). In all bar plots, values indicate the percentage of lesion volume overlapping each atlas-defined region. Despite their distinct anatomical profiles, lesion centroids were separated by only 2.75-3.23 mm, highlighting the millimetre-scale spatial resolution required to distinguish response-associated anatomy within the pallidothalamic territory. **e**-**f)** The spatial overlap between the two responder lesions localized predominantly to Forel’s field H1 (55.8% of the responder-overlap region), a compact region traversed by major pallidothalamic fibre pathways. The responder-overlap region is shown in relation to **e)** the ansa lenticularis and **f)** the fasciculus lenticularis. **DBS fibre-filtering application in Parkinson’s disease: g)** Streamlines associated with clinical improvement in Parkinson’s disease (PD), identified using DBS fibre filtering with the Netstim Atlas v2.2 normative connectome. **h)** Corresponding streamlines identified using DBS fibre filtering with the FOCUS atlas. Compared with g), the more extensive anatomical delineation of fibre tracts in the FOCUS atlas provides a richer representation of local fibre architecture, with exemplary additional trajectories indicated by red arrows. **i)** Validation of the FOCUS-derived fibre-filtering model. Greater similarity between each patient’s stimulation-associated streamline profile and the optimal model shown in **h)** was associated with greater clinical improvement in both the discovery cohort (Spearman’s *ρ* = 0.36, *p* = 0.0002) and an independent test cohort (Spearman’s *ρ* = 0.30, *p* = 0.003).

The conjunction region shared by both responder lesions in template space comprised 197 voxels (corresponding to approximately 0.2 mm^3^ at 100 µm isotropic resolution) and was predominantly located within H1 (55.8% of the overlap), followed by ZI (21.8%), H (3.6%), and H2 (1.5%). This shared responder region localized to a portion of H1 traversed by major pallidothalamic fibre path-ways, including the ansa lenticularis and fasciculus lenticularis (*Figure 7e,f*). Despite these distinct anatomical profiles, lesion centroids were separated by only 2.75–3.23 mm, indicating that out-come-related differences occurred within a highly focal pallidothalamic territory and illustrating the potential utility of high-resolution atlas resources such as FOCUS.

In the second analysis, we repeated a previously published DBS fiber filtering analysis using FOCUS white matter definitions instead of the originally used ‘Netstim atlas’^41,42^. The analysis tested across patients which tracts associated most strongly with optimal response in the motor part of the Unified Parkinson’s Disease Rating Scale (UPDRS-III). It was informed by 129 patients that underwent DBS surgery to the subthalamic nucleus (STN-DBS). Exactly replicating the original analysis in^41^, results were validated by estimating improvements in an independent test cohort of 89 patients operated at different centres.

Using the Netstim Atlas (as used in the original report), motor improvement was primarily associated with modulating streamlines that projected from Brodmann areas 4, 6, and 8 to the STN, which, functionally, partly correspond to the ‘hyperdirect pathway’ (*Figure 7g*). Stimulating these tracts – which were exclusively informed by the discovery cohort – was associated with greater UPDRS-III improvement in the discovery cohort (circular analysis; Spearman ρ = 0.33, p = 0.0004), when subjecting the analysis to ten-fold cross-validation (ρ = 0.29, p = 0.001), and, most critically, in the entirely independent test cohort (ρ = 0.30, p = 0.006).

Repeating the same analysis after replacing fibre definitions from the Netstim Atlas with the ones from the FOCUS atlas identified similar corticosubthalamic projections additionally highlighting importance of subcortical projections such as the anatomical surrogates of the ‘indirect pathway’ (interconnecting internal and external globus pallidus; GPi and GPe; with STN and thalamus), connections with the substantia nigra (SN), pedunculopontine nucleus (PPN,^43^), and centromedian nucleus (CM) (*Figure 7h*). This may demonstrate that original assumptions of exclusive importance of the hyperdirect pathway may have originated since tested tractograms did not include tract definitions of subcortical anatomy ^40,44–46^. While the FOCUS atlas provided a more comprehensive definition of the tracts that correlated with clinical response, its capability of explaining variance in clinical outcomes remained unchanged (discovery cohort circular: *ρ* = 0.36, *p* = 0.0002; 10-fold CV: *ρ* = 0.30, *p* = 0.0008; test cohort: *ρ* = 0.30, *p* = 0.003; *Figure 7i*). In our view, this is not surprising. For reasons of brevity, we point the reader to a recently published extensive treatise on precisely the reasons behind this matter^41^.

## Discussion

Here, we introduce a subcortical atlas that delineates subcortical structures with high accuracy and resolution. The atlas consists of 85 bilateral and 10 midline grey and white matter structures as well as 77 fibre bundles, anchored around detailed 3D definitions derived from an ex-vivo, 100 µm, high resolution brain MRI dataset. These structures were validated against 48 core sources of information, including histology, fibre dissections, and classical anatomical references. Further, these segmentations were registered into standard stereotactic space for precise brain mapping. This process involved a labour-intensive, iterative multi-stage approach that relied on key anatomical landmarks as fiducial anchor structures and software-enhanced manual refinements. The resulting FOCUS atlas provides anatomical definitions at the submillimetre scale. It can be precisely registered to individual subject anatomy. This approach overcomes common issues found in purely histology-driven atlases, such as the challenges of registering two-dimensional histological stacks to MRI scans for deformable mapping^29,30^. By tightly integrating multi-layered anatomical references, the FOCUS atlas largely preserves anatomical definitions at a near-histological level of detail, possibly setting a new standard for precision in neuroanatomical mapping.

Brain atlasing has a long history in the field of neuroscience. In the early days of brain anatomy, such as the times of Auguste Forel (*Figure 5c*), a common procedure was to manually draw (i.e., reproduce) structures seen on stained histology slices through the microscope. While the advent of neuroimaging and MRI brought such classical anatomy work to a temporary decline, impactful reference atlases such as the Schaltenbrandt-Wahren^51^ or Mai^52^ atlases remain important to define surgical targets of intervention, such as in ablative neurosurgery or DBS. With increasing magnetic field strengths and advanced MRI sequences, the concept of ‘direct targeting’ was fostered, which claimed that surgical targets could be seen clearly enough on MRI alone^53–55^. On the one hand, this framework is elegant because it does not require any atlas registration steps, which can introduce bias. On the other hand, surgical interventions in the subcortex fall into highly complex terrains in which numerous structures – which are hardly discernible on MRI – reside in close proximity and complex spatial relationships to one another. The inability to see these structures makes it impossible to study their potential impact on surgical success.

The following examples illustrate that relatively obscure structures need to ‘become visible’ before we can truly model and understand which precise structure is responsible for mediating clinical improvements following surgery. For instance, it remains unclear whether surgical targeting in movement disorders such as Parkinson’s disease, dystonia, and Tourette’s syndrome should optimally fall onto the medial or lateral parts of the internal pallidum^56^, or, as suggested by others^57^, onto the lamina interna which divides the internal and external pallidum, or the lamina accessoria, which subdivides the internal pallidum. Delineating these structures (as is possible with the present atlas) may help shed light on such relationships, as the clinical application examples in the present article already hint at. Similarly, an optimal surgical target region to treat obsessive compulsive disorder has been described in close proximity to the bed nucleus of stria terminalis^58^. As extensively discussed by ^17^, efficacy of this precise location could also be interpreted as the medial forebrain bundle (mfb), the ventral amygdalofugal pathway (including a different DBS target for OCD, the inferior thalamic peduncle) or the substantia innominata. Direct visualization on MRI or the use of tractography does not come close to disentangle such ambiguities, which halts progress in mechanistically understanding the efficacy of surgical applications. As a third example, surgical targeting for tremor has included the ‘ventral intermediate nucleus’ (VIM) of the thalamus (which itself has been questioned by some researchers, who considered it more of a transitional zone between afferent pallidal and cerebellar fibres rather than a distinct entity^59^), as well as the area ventral to it, which has been referred to as the zona incerta by some^60^ or pragmatically dubbed the ‘posterior subthalamic area’ (PSA) by others (for a review, see^61^). This precise location is filled with potential candidate structures, such as incoming cerebellar projection fibres^59,61,62^, pallidal projections of passage that merge in Forel fields^63^, the subthalamic nucleus proper^42^ or the zona incerta^60^, which have all been proposed to mediate tremor relief, independently. Before we can identify the structures that are truly associated with tremor, we need to make them visible, in the first place.

As DBS targets continue to expand and become more refined across a growing range of neurological and psychiatric indications, precise anatomical mapping of both intended and off-target stimulation effects is becoming increasingly critical for clinical interpretation and translation. In established applications such as subthalamic nucleus stimulation for Parkinson’s disease – where stimulation volumes often extend into neighbouring structures – the high spatial resolution of our atlas in this region may offer new insights into the substrates of desired and unintended effects. The exemplary DBS fiber filtering analysis presented here already gives an example for extended anatomical insight of the structures that mediate DBS effects. Moreover, for emerging targets located near the Fields of Forel^64,65,66^, our detailed anatomical delineation and precise normalization may help clarify structural-functional relationships and contribute to future clinical discoveries, as exemplified by the case series of dystonia patients that underwent ablative neurosurgery. DBS leads to spill-over to neighboring structures, which often leads to side-effects of various forms. One example are DBS-induced ocular motor phenomena. Precise anatomical mapping of these “off-target” effects in the midbrain are even used to guide optimal electrode placement and programming in depression^67^, and have led to serendipitous findings in the context of PD^68^ or Tourette syndrome^69^. By providing a detailed map of relevant subcortical zones implicated in neuromodulation, the FOCUS atlas may supply the anatomical groundwork needed to relate stimulation sites both to therapeutic benefit and to potential off-target effects.

Beyond the impact on the field of neuromodulation, applications of the FOCUS atlas may extend to functional neuroimaging studies that focus on subcortical activations or brain connectivity. In recent years, the subcortex has become a new frontier in the field of functional neuroimaging^1,70^, particularly with concepts such as precision functional mapping^3,71^ (PFM) on the rise. Here, extensive scan times enhance the typically low subcortical signal to noise to map subject-specific functional territories^3,72^ – even in ways that may one day become relevant for brain surgery^73^. Similarly, it has been proposed to increasingly use ultra-highfield MRI to map subcortical structures to further our mechanistic understanding of the subcortex^74^ and to gain enhanced surgical precision^75^. At first glance, these concepts may seem orthogonal to the use of refined brain atlases, since their aim is to uncover subject-specific regions precisely without using atlases.

However, no currently available form of in-vivo imaging – be it structural or functional – comes close to the resolution of the presented atlas, which is a synthesis of numerous resources that will almost certainly never become available for a single human living brain. Instead, we argue for a combined use of i) high-resolution individualized imaging such as PFM and ultra-high-field imaging together with ii) high-resolution atlases such as the present one to truly move the fields of neurosurgery and subcortical neuroimaging forward. Deep learning based applications such as the promising Next-Brain platform already allow seamless integrations between high-resolution atlas-based priors and individual neuroanatomy^8^.

Our atlas includes high resolution definitions of grey and white matter structures in voxelized format. In addition, we included 77 fibre tracts that represent pathways of the subcortical wiring diagram in streamline format, which resembles the output structure of common diffusion-MRI based tractography experiments^76^. However, instead of relying on diffusion weighted imaging, we deliberately chose to follow a novel concept introduced by Petersen et al.^40^, which aims at reconstructing tracts based on various sources of information (such as tracing studies, gross-dissection and literature results as used here). To contrast this method from diffusion-based tractography and avoid confusion due to its similar appearance, we propose to term this method ‘tract architectonics’, since the aim is to create tract models in the computer similar to how architects plan buildings. In essence, the method is supported by software tools that allow delineating polynomial curves relative to landmarks. Based on these and parameters that control fanning and thickness of tracts, and, at times, regions of inclusion, the computer then samples streamlines along these curves in such a way that they best fit the anatomical description of tracts. The rationale to choose this method for tract reconstructions in the subcortex has been well outlined in the pioneering paper by

Petersen and colleagues^40^. In brief, diffusion-MRI based tractography suffers from numerous problems that make the method poorly suitable to reconstruct thin and short tracts with complex trajectories that cross with each other and fan out after entering grey matter^77,78^. Diffusion-tractography is inherently limited by a high-rate of false-positive fibres^78^, coarser spatial resolution compared to the underlying axonal architecture^79–81^, and the difficulty in resolving crossing fibres^78^, especially in regions where the diffusion signal is biased towards highly myelinated fibres. These limitations are especially prominent for the subcortical connections, which are thin, dispersed and often cross the highly myelinated internal capsule. In the most prominent study that analyzed shortcomings of the method to date^78^, the authors concluded that ‘tractography should increasingly employ reliable anatomical priors from ex vivo histology, high-resolution postmortem DWI, or complementary electrophysiology for optimal guidance’ – which is precisely the approach taken here.

Several limitations are important when interpreting our findings and when using the atlas in future neuroscientific studies. First, it is anchored onto the anatomy of a single post-mortem brain and may not reflect interindividual variability including gender– and age-related differences or postmortem tissue changes. Despite our efforts to mitigate these effects through deformable registration into standard space and cross-validation to numerous different anatomical resources, this limitation could not be completely averted. Future work could incorporate additional specimens and complementary in vivo imaging to systematically quantify anatomical variability across populations. Second, we deliberately chose to map several zones of clinical interventions in greater detail than other subcortical regions, and the atlas does not represent cortical anatomy beyond cortico-subcortical white matter projections. This choice was deliberate since mapping the entire brain with the precision and care introduced here would go beyond the scope of any single scientific project. However, we openly release the atlas, associated software tools and protocols in a version-controlled fashion, which will allow contributions, refinements and extensions by others, in the future. Third, while the atlas is intended to be used for precision functional mapping studies of the subcortex^3,73^, as well as analyses of functional neurosurgery^17,82^, the clinical analyses presented here are intended as illustrative proof-of-concept applications rather than formal clinical validation studies. The lesion analysis comprises only three selected cases and is intended to demonstrate the anatomical resolution with which small neighbouring treatment sites can be distinguished. Similarly, the DBS fiber-filtering analysis illustrates how an expanded subcortical tract repertoire can alter anatomical interpretation while yielding predictive performance comparable to the previously published model.

Importantly, the present first release of FOCUS is focused on subcortical anatomy and does not constitute a uniformly refined whole-brain atlas. While the ex vivo dataset was registered to MNI space at the whole-brain level, iterative manual refinement of the transformation was deliberately concentrated on the subcortical regions prioritized in this work. Accordingly, anatomical correspondence of this template is expected to be highest in these targeted subcortical zones, whereas regions outside them reflect the underlying automated whole-brain normalization but were not subjected to the same degree of dedicated refinement. Despite this, all areas covered by the FOCUS atlas segmentations should be considered equally precise.

Cortical anatomy is therefore not within the scope of the present release, except for selected cortical seed regions that were defined directly in MNI space for tract modelling. Possibly, in the future, the atlas will be extended to also support cortical segmentations. However, numerous excellent cortical atlases exist, such as the Julich Brain atlas^25^ that represents decades of work in a modern cytoarchitectonic anatomy centre.

Finally, our segmentations are currently limited to structural anatomy and do not incorporate ‘functional’ zones, such as motor, associative and limbic divisions of thalamus and basal ganglia. Such information is present in other resources such as the DISTAL^15^ or Melbourne Subcortex Atlases^83^ and could be included as an additional ‘layer’ in future versions of FOCUS.

In summary, by combining unprecedented anatomical detail with a dedicated tool that allows precise deformability to individual subject space, interactive refinement, and tract-generation tools, we provide a foundational atlasing platform for precision neuroimaging and neuromodulation. This atlas not only bridges the resolution between ex-vivo 7T research and routine clinical imaging, but also empowers investigators to map and manipulate subcortical circuits with submillimetre fidelity.

## STAR★Methods

Detailed methods are provided in the online version of this paper and include the following:

• Key Resources Table
• Method Detail
  ○ Manual segmentation of grey and white matter brain structures
  ○ Registration to MNI space using advanced concepts
  ○ Manually curated Pathway Atlas
  ○ Lesion Study
  ○ DBS Fiber Filtering
• Ethics Statement
• Data and Code Availability
• Materials Availability

## Acknowledgements

The authors would like to thank Drs. Martin Parent and Abbas Sadikot for their helpful guidance on tract results in comparison to original and unpublished tract tracing data as well as for sharing their long-standing experience in variability of pallidothalamic tract tracing results across individuals. H.F. was supported by the Carl-Duisberg Stipend of the Bayer Foundation and the Graduate School of Life Sciences Wuerzburg (SFB Retune). M. U. F. is supported by the Jung Stiftung für Forschung und Wissenschaft (Career Advancement Prize 2024), the Manfred und Ursula Müller-Stiftung and the German Society of Neurology (NeuroTech Innovationspreis 2024) and the “Wissenschaftliches Oberarztprogramm” at the University of Ulm. A.H. was supported by the Schilling Foundation, the German Research Foundation (Deutsche Forschungsgemeinschaft, CRC-1451, 431549029 and CRC-1270 ELAINE, 3–299150580), and the National Institutes of Health (R01MH130666, 1R01NS127892-01, 2R01 MH113929 & UM1NS132358). B.L.E. was supported by the Chen Institute MGH Research Scholar Award, Jean Perkins Foundation, NIH Director’s Office (DP2HD101400), NINDS (U01NS137484, R21NS109627), and Department of Defense (W81XWH2210999). N.L. is supported by the Fundamental Research Funds for the Central Universities, the National Natural Science Foundation of China (Grant No. 5113250169).

## Author contributions

H.F. conceptualized the atlas framework and warping strategy underlying this study, contributed to the conceptual development of the WarpDrive tools, created the voxelized high-resolution atlas and tracts, performed the primary data analyses, created the visualizations and renderings, wrote the manuscript, and contributed to funding acquisition. A.H. created the atlas, performed data analyses, conceptualized and supervised the study, conceptualized WarpDrive tools, and the CurveToBundle tool, wrote the manuscript, and provided funding for the study. I.A.S. contributed to the conceptual development of the CurveToBundle tool, maintained the tract database, created tracts, and assisted with dataset curation. S.O. conceptualized and implemented the software necessary for warping and the CurveToBundle tool. A.K.E.H., E.J.L.A., V.M.H., and B.L.E. provided anatomical data and helped with the interpretation of anatomical structures. E.J.L.A. additionally contributed to the interrater analysis. M.B. and E.K. assisted in MR signal analysis and anatomical delineation of substantia nigra boundaries. N.R., G.M., and L.G. created white matter structures of the atlas. L.K. and R.D. contributed manual segmentations for the interrater analysis. N.R., J.P., C.H., and A.G. assisted with data analysis. M.M. collected clinical data and material for an illustrative application of the atlas and performed associated analyses. P.Z. contributed to the DBS fiber-filtering application of the atlas. N.L. assisted with technical support. M.U.F. contributed code, supported anatomical interpretation, and performed analysis. S.H. provided data and helped to analyse and interpret the data. J.V., C.M., C.N., B.L.E., and M.D.F. provided supervision.

## Declaration of interests

A.H. reports lecture fees for Boston Scientific, is a consultant for and holds stock options of Modulight.bio, was a consultant for FxNeuromodulation in recent years and serves as a co-inventor on a patent granted to Charité University Medicine Berlin that covers multisymptom DBS fibrefiltering and an automated DBS parameter suggestion algorithm unrelated to this work (patent #LU103178). M.U.F. has served as a consultant for CereGate GmbH and Tovly LLC, and as an associate editor for NPJ Parkinson’s disease, unrelated to the submitted work. The remaining authors declare no competing interest.

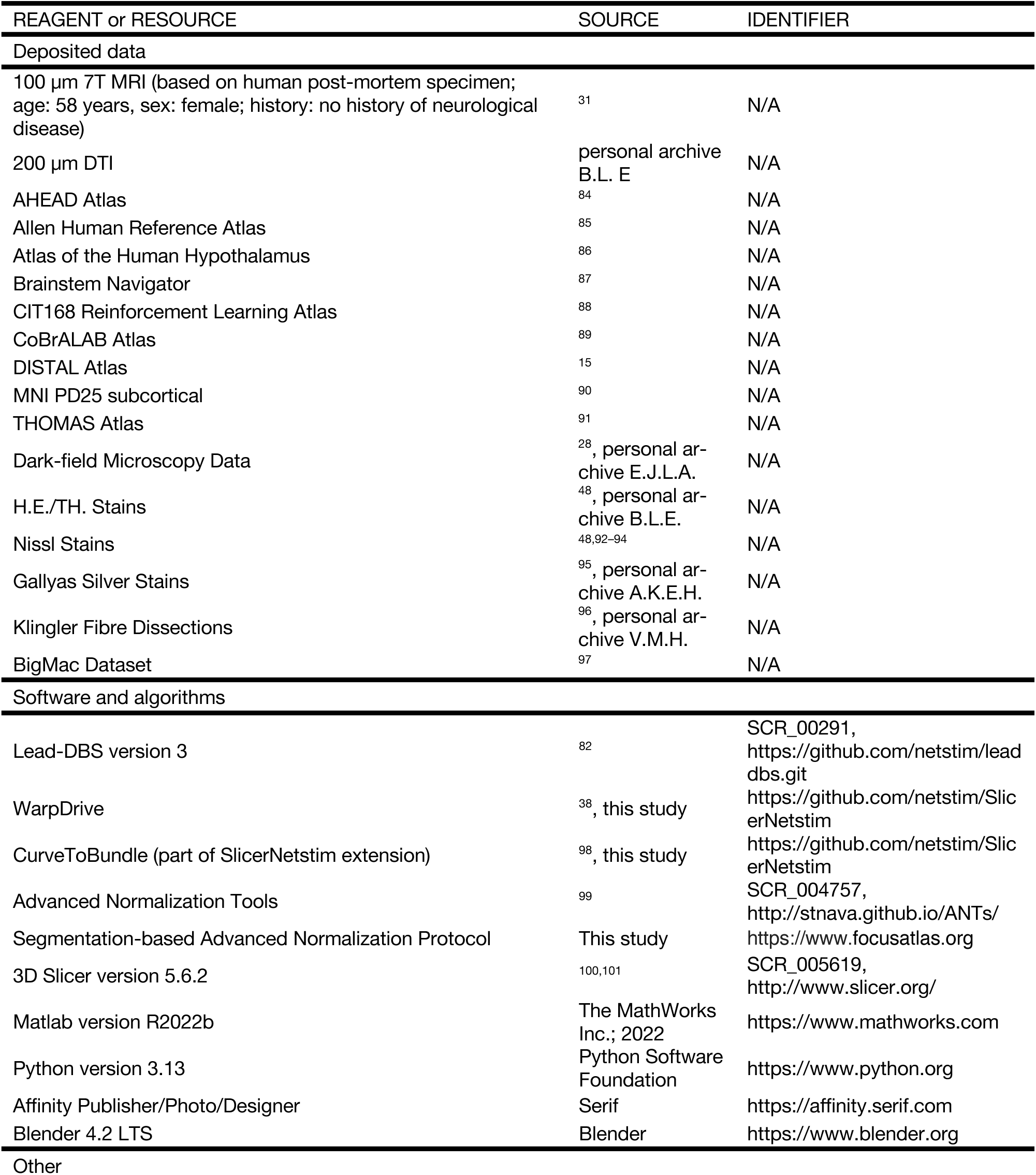
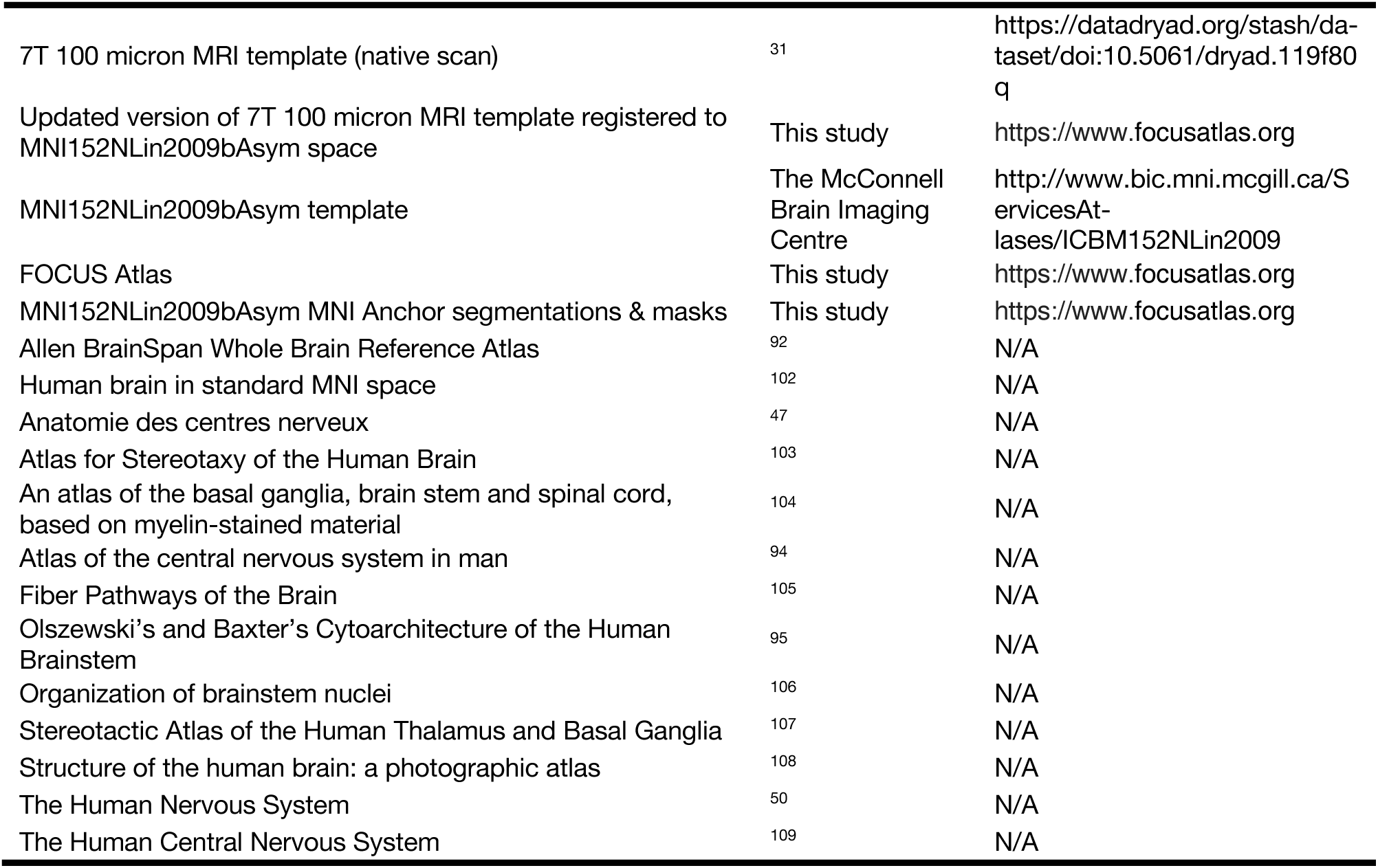
Key Resources Table.

Key Resources Table
| REAGENT or RESOURCE | SOURCE | IDENTIFIER |
| --- | --- | --- |
| Deposited data |  |  |
| 100 μm 7T MRI (based on human post-mortem specimen; age: 58 years, sex: female; history: no history of neurological disease) | <sup>31</sup> | N/A |
| 200 μm DTI | personal archive B.L. E | N/A |
| AHEAD Atlas | <sup>84</sup> | N/A |
| Allen Human Reference Atlas | <sup>85</sup> | N/A |
| Atlas of the Human Hypothalamus | <sup>86</sup> | N/A |
| Brainstem Navigator | <sup>87</sup> | N/A |
| CIT168 Reinforcement Learning Atlas | <sup>88</sup> | N/A |
| CoBrALAB Atlas | <sup>89</sup> | N/A |
| DISTAL Atlas | <sup>15</sup> | N/A |
| MNI PD25 subcortical | <sup>90</sup> | N/A |
| THOMAS Atlas | <sup>91</sup> | N/A |
| Dark-field Microscopy Data | <sup>28</sup>, personal archive E.J.L.A. | N/A |
| H.E./TH. Stains | <sup>48</sup>, personal archive B.L.E. | N/A |
| Nissl Stains | <sup>48,92–94</sup> | N/A |
| Gallyas Silver Stains | <sup>95</sup>, personal archive A.K.E.H. | N/A |
| Klingler Fibre Dissections | <sup>96</sup>, personal archive V.M.H. | N/A |
| BigMac Dataset | <sup>97</sup> | N/A |
| Software and algorithms |  |  |
| Lead-DBS version 3 | <sup>82</sup> | SCR_00291, https://github.com/netstim/lead-dbs.git |
| WarpDrive | <sup>38</sup>, this study | https://github.com/netstim/SlicerNetstim |
| CurveToBundle (part of SlicerNetstim extension) | <sup>98</sup>, this study | https://github.com/netstim/SlicerNetstim |
| Advanced Normalization Tools | <sup>99</sup> | SCR_004757, http://stnava.github.io/ANTs/ |
| Segmentation-based Advanced Normalization Protocol | This study | https://www.focusatlas.org |
| 3D Slicer version 5.6.2 | <sup>100,101</sup> | SCR_005619, http://www.slicer.org/ |
| Matlab version R2022b | The MathWorks Inc.; 2022 | https://www.mathworks.com |
| Python version 3.13 | Python Software Foundation | https://www.python.org |
| Affinity Publisher/Photo/Designer | Serif | https://affinity.serif.com |
| Blender 4.2 LTS | Blender | https://www.blender.org |
| Other |  |  |
| 7T 100 micron MRI template (native scan) | 31 | https://datadryad.org/stash/dataset/doi:10.5061/dryad.119f80q |
| Updated version of 7T 100 micron MRI template registered to MNI152Nlin2009bAsym space | This study | https://www.focusatlas.org |
| MNI152Nlin2009bAsym template | The McConnell Brain Imaging Centre | http://www.bic.mni.mcgill.ca/Services/At-lases/ICBM152Nlin2009 |
| FOCUS Atlas | This study | https://www.focusatlas.org |
| MNI152Nlin2009bAsym MNI Anchor segmentations & masks | This study | https://www.focusatlas.org |
| Allen BrainSpan Whole Brain Reference Atlas | 92 | N/A |
| Human brain in standard MNI space | 102 | N/A |
| Anatomie des centres nerveux | 47 | N/A |
| Atlas for Stereotaxy of the Human Brain | 103 | N/A |
| An atlas of the basal ganglia, brain stem and spinal cord, based on myelin-stained material | 104 | N/A |
| Atlas of the central nervous system in man | 94 | N/A |
| Fiber Pathways of the Brain | 105 | N/A |
| Olszewski's and Baxter's Cytoarchitecture of the Human Brainstem | 95 | N/A |
| Organization of brainstem nuclei | 106 | N/A |
| Stereotactic Atlas of the Human Thalamus and Basal Ganglia | 107 | N/A |
| Structure of the human brain: a photographic atlas | 108 | N/A |
| The Human Nervous System | 50 | N/A |
| The Human Central Nervous System | 109 | N/A |

For key resources, N, age, sex and clinical history are detailed in *Supplementary Table 2*.

## Manual segmentation of grey and white matter brain structures

All manual atlas segmentations, as well as the iterative visual assessment and refinement of the native-to-MNI registration, were performed by H.F. to maintain consistency throughout the delineation and registration process. Segmentations were primarily performed on the 100-µm ex vivo MRI. Where anatomical boundaries were ambiguous, complementary anatomical resources, including histological stacks, were consulted by matching the corresponding sectioning plane. Example segmentation protocols for several structures are provided in the Supplement to illustrate the general delineation approach and the use of complementary anatomical information. A subset of structures was additionally segmented by three additional raters to assess interrater reproducibility.

The anchor image used to create the high-resolution subcortical brain atlas was based on the open-source 100-micron ex-vivo MRI brain scan of a 58-year old woman with no history of neurological disease^110^. Segmentations were primarily guided by this synthesized Fast Low Angle Shot (FLASH) volume in native space. Additionally, when needed, individual volumes acquired at flip angles of FA15°, FA20°, FA25°, and FA30° were used to inform specific boundaries, for example when delineating nigrosomes.

Spatial normalization was performed to the asymmetrical ICBM152 2009 release b (MNI152NLin2009bAsym,^111,112^), which also serves as the default reference template in Lead-DBS (^113^; www.lead-dbs.org). For most structures, the T1-weighted version was used as the primary reference, in addition to the T2-weighted template. Comprehensive procedural details are provided in a detailed protocol available at the data repository (https://www.focusatlas.org). All anatomical structures were manually segmented in 3D-Slicer (Version 5.6.2; https://www.slicer.org,^101^) at 100 µm resolution, based on the native FLASH scan described above. Delineation was performed in axial, coronal, and sagittal planes, and segmentations were iteratively refined in all orientations. Paired structures were segmented bilaterally in both cerebral and cerebellar hemispheres. A standardized anatomical delineation protocol was developed and adapted from previous work^114, 115^ (*see detailed protocol available at the data repository*). In brief, the initial focus was on well-defined structures with distinct contrast boundaries, including the red nucleus (RN), subthalamic nucleus (STN), globus pallidus (GP), and striatum. Throughout the process, boundary precision was iteratively improved across planes.

Given the signal complexity and resolution of the FLASH sequence, structures typically considered straightforward – such as the putamen or substantia nigra – required critical re-evaluation. To ensure anatomical validity, all segmentations were reviewed by a multidisciplinary team of neuroanatomists (A.K.E.H., V.M.H.), neurosurgeons (E.J.L.A., V.M.H.), and MR physicists (E.K., M.B.), who assessed contrast profiles and structural definitions collaboratively (see *Supplementary Materials* and *detailed protocol at data repository*).

To refine the segmented structures while preserving fine anatomical details, a multi-step iterative smoothing approach was applied using the Segment Editor module in 3D Slicer (^116^). First, median filtering (kernel size: 0.2 mm ± 0.1 mm) was applied to reduce voxel-level noise while preserving edge sharpness. Next, joint smoothing (smoothing factor 0.4–1) was applied to selected regions, allowing fine adjustment of structural boundaries without over-regularization. When needed, a final Gaussian smoothing step (σ = 0.2 mm ± 0.1 mm) provided subtle surface refinement. All boundaries were manually reviewed and adjusted in each slice orientation. This process was repeated iteratively until a balance was achieved between sharp anatomical definition and overall surface smoothness.

To establish a robust anatomical framework, we initially employed widely used neuroanatomical atlases (see *Supplementary Materials*) as references to delineate large, well-characterized structures. This provided a coarse anatomical scaffold that was subsequently refined in a stepwise manner, moving from general boundaries to precise segmentations. Once major structures were defined, each region was individually reassessed using a combination of neuroanatomical expertise, classical reference atlases, anatomical textbooks, fibre dissection data, and histological material. A full list of segmented structures and delineation methods is provided in the *Supplementary Materials*.

For each structure, a tailored approach was taken to achieve the highest possible anatomical accuracy. Histological preparations with specific staining and processing techniques (e.g. Gallyas/ Nissl staining, dark-field microscopy) were selected according to the anatomical characteristic of the target region. To ensure precise cross-modality correspondence, the volume was sliced in oblique angles using 3D-Slicer to match the slice orientation with the one of reference sections. Multiple anatomical landmarks were used across slice orientations to ensure accurate topographic mapping and alignment. Particular attention was given to the mesencephalic–diencephalic transition zone, a region of high anatomical complexity and clinical relevance. Conventional histological sectioning often introduces local distortions in this area, leading to blind spots in many existing resources. We therefore applied a systematic reconstruction strategy and iterative annotation process to produce a complete, continuous, and topologically coherent description of this region. Special care was also taken to delineate adjacent structures relevant to current and emerging DBS targets, to provide continuous anatomical coverage without segmentation gaps.

### Registration to MNI space using advanced concepts

To enable application of the high-resolution atlas in clinical neuroimaging and patient-specific DBS research, the native ex vivo dataset had to be nonlinearly registered to standard stereotactic (ICBM 2009b NLIN Asym; ‘MNI’) space. For later applications to be meaningful, this process had to be as precise as the original delineations of anatomical structures itself.

A previously published version of this nonlinear warp has been introduced^110^ but is not precise in various subcortical regions (see *Supplementary Materials*). Our primary focus was on improving the transformation in clinically relevant brain regions, such as the thalamus, subthalamus, mesencephalon, and cerebellum. The following outlines the procedure used to generate the transformation.

First, the FLASH volume was downsampled to isotropic voxel sizes of 0.5 mm. A linear transformation was then applied using the built-in 3D-Slicer transform tool to align the high-resolution image to the MNI reference space. This transformation adjusted the spatial coordinates of the image to match the MNI standard. The MR image was then normalized into template space using a multispectral and multi-step approach with Advanced Normalization Tools (ANTs,^99^). This process included five stages: After two linear (rigid followed by affine) steps, a nonlinear (whole brain) SyN-registration stage was followed by two nonlinear SyN-registrations that consecutively focused on the area of interest as defined by subcortical masks in ^117^. The registration was conducted with the “Effective: Low Variance, Default” preset.

Once this automated warp was generated, it was manually improved to maximize precision: First, a total of n = 62 corresponding anchor segmentations were manually created (n = 31 for each hemisphere) in both the high-resolution native space and the MNI template. Critically, segmentations in native space were done to match the visible components delineated in MNI space, i.e. these were additional segmentations carried out on top of the anatomical segmentations discussed above. These paired segmentations were carefully selected based on anatomical structures that were clearly identifiable in both templates and served as fiducials throughout an iterative normalization process consisting of n = 37 steps. After each iteration, alignment between native and MNI space was assessed by expert visual inspection in axial, coronal, and sagittal views, with particular attention to the correspondence of anchor structures, visible anatomical landmarks, and the topographic relationships of neighbouring regions. Where residual mismatch was identified, the transformation was further refined. This process was repeated until additional iterations no longer yielded meaningful improvement in the brain regions of interest (for details see *Supplementary Materials* and *detailed protocol at data repository*).

Given the difficulty of distinguishing smaller nuclei, such as differentiating parabrachial pigmented nucleus and paranigral nucleus vs. substantia nigra in MNI space, we created larger, coarser segmentations that grouped these regions together. These masks (see *detailed protocol at data repository*) were manually defined in both the native and MNI spaces, ensuring that even small nuclei, which are not visible in the MNI space, could accurately be transformed. Following this, we visually inspected the alignment of the single structures to confirm that the topography and arrangement of the structures were accurately represented.

To further address residual misalignments between the source and target spaces, which sometimes persisted despite the aforementioned procedure, we extended and employed the WarpDrive toolbox (^38^) as shown in *Figure 2c*. This tool allows for manual, localized corrections of the transformation. For the current study, we extended the capabilities of the toolbox by incorporating additional functions, such as the ‘shrink and expand’ module.

### Manually curated Pathway Atlas

Once the high-resolution brain structures were precisely mapped in standard space, we manually constructed anatomically accurate tracts intended for use in studies that require precise definitions of subcortical white matter pathways^40,42^. Instead of relying on commonly used fibre tractography based on diffusion MRI, we followed the pioneering concept by the McIntyre group, which consists of manually reconstructing pathways based on anatomical priors and anatomical knowledge^40^. To contrast this method from diffusion-based tractography and avoid confusion due to the similar appearance of these manually created streamlines, we propose to term this method ‘tract architectonics’, since the aim is to create tract models in the computer similar to how architects plan buildings.

To facilitate such manual tract reconstructions, we developed an open-source 3D-Slicer extension, *CurveToBundle* (https://github.com/netstim/SlicerNetstim). This module generates fibre tracts from user-defined curve trajectories by uniformly distributing streamlines along the curve with adjustable radii and jitter. After delineation of the curve trajectories, histology-based atlases were used to further define the start, end, pass-through and avoidance regions. For instance, for fasciculus lenticularis models, any streamline that doesn’t pass through Field of Forel H2 was discarded, and a new streamline was sampled until the criteria is met. The resulting bundle was trimmed to start at internal globus pallidus. For each bundle, a standardized streamline count was sampled along the curve: 1000 for large cortico-cortical and cortico-subcortical tracts, 500 for major subcortical projections, and 250 for smaller bundles. Bundles were generated on the left hemisphere and were flipped using nonlinear transformation.

Tracts were first based on direct white matter segmentations, leveraging the exceptional contrast of the 7T high-resolution template to its fullest potential. Additionally, the dense representation of core regions facilitated the construction of tracts in accordance with established literature.

In the tract creation process, we also relied on Klingler fibre dissections that were deliberately created for the present project (V.M.H.), histological data (B.L.E., A.K.E.H, E.J.L.A), and textbook references to ensure anatomical precision and validation. In addition, we cross-referenced our data to openly available high-resolution 600-micron DTI data ^97^. The advantage of this manual approach lies in its ability to identify and delineate very small subcortical tracts, particularly those near the midline, such as for example the pallidohypothalamic tract (with a diameter of less than 2 mm)^40^. Moreover, this approach enables the precise reconstruction of small branching tracts, such as the fasciculus retroflexus, as well as pathways with complex trajectories, like the oculomotor nerve *(Figure 6)*. Finally, the technique allows to recreate tracts that enter and diffusely project within grey matter zones, which are impossible to reconstruct based on diffusion MRI given low anisotropy in these regions. The resulting pathways exemplify cases in which conventional diffusion MRI-based tractography struggles due to crossing and dispersing fibres, leading to signal loss – such as when the oculomotor nerve traverses the capsule of the red nucleus. Even in the 100-micron 7T template, such pathways were challenging to trace. However, by segmenting their visible portions step by step and extending the knowledge using the additional resources mentioned above, we were able to reconstruct even such fine tracts using our manual approach.

### Lesion study

We retrospectively analysed three patients with dystonia who underwent unilateral radiofrequency pallidothalamic tractotomy (PTT; target: field H1 of Forel) at Tokyo Women’s Medical University, selected to span the range of clinical outcomes. Motor status was assessed using the Burke-Fahn-Marsden Dystonia Rating Scale movement subscale (BFMDRS-M): Case 1 (34-year-old woman, left-sided treatment, segmental dystonia), 59 to 6 (89.8% improvement); Case 2 (48-year-old woman, left-sided treatment, segmental dystonia), 9 to 3 (66.7% improvement); and Case 3 (66-year-old man, right-sided treatment, focal cervical dystonia), 8 to 8 (0% improvement).

Tractotomy lesions were manually segmented as T1-hyperintense regions on immediate-postoperative T1-weighted MRI using MRIcroGL and normalized to MNI152 NLIN 2009b asymmetric space using the default parameters for ANTs^99^ based normalization implemented in Lead-DBS. Binary lesion masks were superimposed with FOCUS atlas structures. To quantify lesion overlaps, individual atlas structures were resampled into lesion space using nearest-neighbour interpolation. For each lesion, overlap with individual FOCUS atlas structures was quantified as the number and proportion of lesioned voxels contained within each structure. In addition, the distance between each lesion centroid and field H1 was calculated. Responder overlap was defined by voxel-wise summation of the two responder lesion masks followed by thresholding at >1, thereby retaining only voxels shared by both lesions. The resulting overlap mask was intersected with each atlas structure, and we quantified the number of intersecting voxels, the percentage of the responder-overlap region contained within each structure, and the percentage of each atlas structure covered by the overlap.

To contextualize the spatial scale and relative location of the lesions, lesion volumes, bounding-box extents, centroids, and pairwise centroid-to-centroid distances were calculated. Centroids were defined as the mean coordinates of all lesioned voxels transformed into millimetre space using the NIfTI affine. Pairwise Euclidean centroid distances were summarized descriptively for the responder-responder and responder-non-responder lesion pairs.

### DBS Fiber Filtering

We repeated a DBS fiber filtering analysis in a previously described dataset of patients with Parkinson’s disease who underwent subthalamic nucleus deep brain stimulation (STN-DBS; ^41,42^). The discovery cohort comprised 129 patients from three international centres (Berlin, Würzburg, and Amsterdam; mean age, 57.4 ± 9.6 years), and the independent test cohort comprised 89 patients from two centres (Beijing and Würzburg; mean age, 60.5 ± 8.9 years). Clinical improvements were defined as relative improvements in the motor part of the Unified Parkinson’s Disease Rating Scale (UPDRS-III), calculated as (preoperative score – postoperative score) / preoperative score.

DBS fiber filtering followed the preprocessing, parameter settings, and validation strategy described in ^41^. The analysis was performed separately using the Netstim Atlas v2.2 normative connectome (as in the original study) and after replacing streamline definitions with the ones of the FOCUS atlas – i.e. with the connectome constituting the only methodological difference between the two analyses.

Methods were identical with those in the original report ^41^. In brief, electrode locations were reconstructed using Lead-DBS v3.1 in MATLAB R2022b (The MathWorks Inc., Natick, MA) according to a standardized protocol (^82^) and normalized to ICBM 2009b Nonlinear Asymmetric (“MNI”) space for group-level analysis. Stimulation volumes were modelled using Open-Source Simulation (OSS)-DBS v2 as continuous electric-field (E-field) distributions.

For DBS fibre filtering, E-fields were thresholded at 100 V/m and associated with streamlines from the respective connectome. At the group level, only streamlines reaching a peak activation of 500 V/m in at least 3% of patients in the discovery cohort were retained. For each retained streamline, peak E-field magnitude across patients was correlated with clinical improvement using Pearson correlation, and the resulting correlation coefficient was assigned to that streamline. The 1,500 streamlines with the strongest positive and 1,000 with the strongest negative associations were retained to construct the final model.

Separate models were generated for the Netstim Atlas v2.2 and FOCUS atlas and evaluated in the discovery cohort, using ten-fold cross-validation, and in the independent test cohort. For consistency across connectomes, model performance was summarized using Spearman correlation coefficients between model-derived fibre scores and clinical improvement.

### Ethics Statement

The retrospective lesion analysis was approved by the Ethics Committee of Tokyo Women’s Medical University (vote 2024-0084). In accordance with the Japanese Ethical Guidelines for Medical and Biological Research Involving Human Subjects, the requirement for written informed consent was waived and an opt-out procedure was implemented. All imaging data were anonymized prior to analysis.

The DBS fiber filtering analysis used previously collected and published clinical datasets. Ethical approval and informed-consent procedures for the participating cohorts were obtained at the respective contributing centres as described in the original studies (^41,42^).

Ethical approval for reanalysis (secondary use of deidentified research data) was obtained from the Ethics Committee of University Hospital Cologne (vote 25-1342).

## Data and code availability

All original code has been deposited to https://www.focusatlas.org and will be made publicly available on the day of the peer-reviewed publication. Additional code used in the preparation of this study is publicly available at https://github.com/netstim; especially https://github.com/netstim/SlicerNetstim and https://github.com/netstim/leaddbs. DOIs are listed in the key resources table.

## Materials availability

Atlas data as well as a meticulously curated protocol of the segmentation process and microscopic, macroscopic, and textbook correlations is made openly available at https://www.focusatlas.org on the day of the peer-reviewed publication. The **F**lexible **O**pen atlas for **C**linical **U**se in the **S**ubcortex (FOCUS) has been primarily implemented for use in Lead-DBS (^82^) but can be broadly applied in any tool that reads in the commonly used NIfTI format. An updated and refined MNI version of the ex vivo 7T MRI 100-micron template^110^ is also available for download within Lead-DBS and at https://www.focusatlas.org.

## Declaration of generative AI and AI-assisted technologies in the manuscript preparation process

During the preparation and revision of this work, the authors used ChatGPT (OpenAI) for language editing. The authors reviewed and edited the output as needed and take full responsibility for the content of the published article.

## Notes

### Competing Interest Statement

M.U.F. has served as a consultant for CereGate GmbH and Tovly LLC, unrelated to the submitted work. A.H. reports lecture fees for Boston Scientific, is a consultant for and holds stock options of Modulight.bio, was a consultant for FxNeuromodulation in recent years and serves as a co-inventor on a patent granted to Charite University Medicine Berlin that covers multisymptom DBS fibrefiltering and an automated DBS parameter suggestion algorithm unrelated to this work (patent #LU103178). The remaining authors declare no competing interest.

### Summary of Updates

Most importantly, we have expanded the quantitative validation and benchmarking of the atlas, including detailed inter-rater analyses, a comprehensive segmentation protocol, and additional comparisons of our delineations with histological and other independent anatomical reference sources. We further illustrate the application of the atlas across heterogeneous in vivo brain anatomies spanning different ages, diagnoses, and acquisition settings, and provide quantitative descriptors of atlas fit and inter-individual variability after patient-specific registration. Finally, we include two applications of the atlas in a clinical context. First, we present a lesion study of three patients with dystonia showing markedly different clinical outcomes and demonstrate how their lesions relate to the targeted pallidothalamic anatomy, including field of Forel H1. Second, we repeat a previously published multi-center DBS fiber-filtering analysis and show how the white matter definitions of the FOCUS atlas provide a more anatomically differentiated interpretation of pathways associated with clinical improvement in Parkinson disease.

