## Supplementary Material for "A precise atlas of the human subcortex"

Friedrich et al., 2026

### Supplementary Materials

#### Supplementary Figures

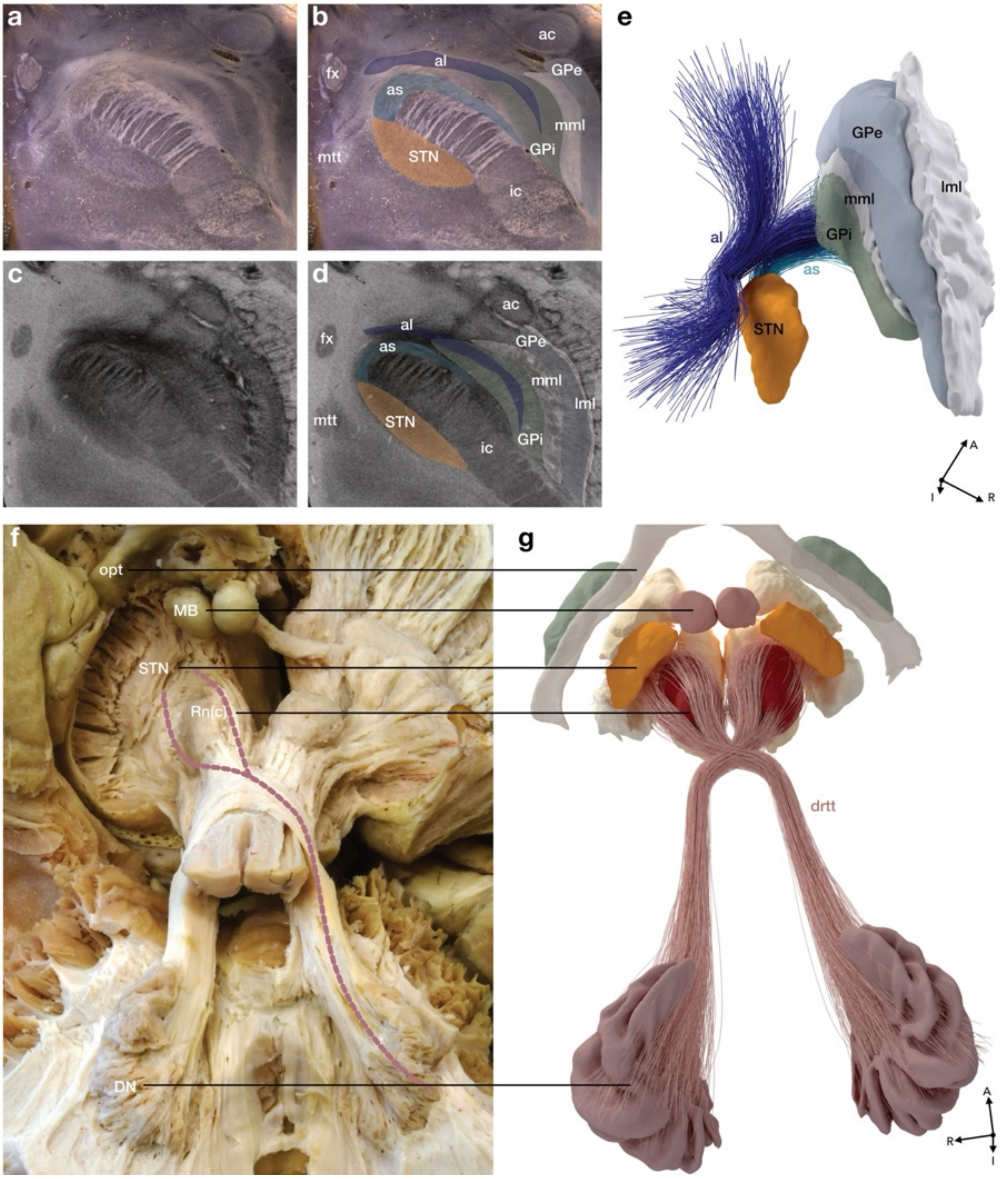

***Supplementary Figure 1.*** **Manual tract reconstruction based on high-resolution atlas segmentations.** Using anatomical ground-truth data and the resources listed in the *Supplementary Notes*, grey and white matter segmentations were delineated in the high-resolution MRI template. After transferring these segmentations into MNI space, anatomical landmarks served as anchors for manual tract generation. **a** Axial section of a dark-field microscopy image used to **b** identify regions of interest, which guided their localization in the MRI. **c**-**d** Corresponding MRI sections were cut to match the histological plane precisely. **e** Three-dimensional rendering of high-resolution atlas structures showing the anatomically faithful reconstructions of the ansa lenticularis and ansa subthalamica. **f** Example of the integration of white matter fibre dissections: reconstruction of the decussating dentato-rubrothalamic tract (drtt), originating in the dentate nucleus (DN), crossing at the level of the midbrain, and passing through the capsule of the red nucleus (RNc) toward its thalamic targets (not dissected here). **g** Accurate segmentation of reference structures such as the DN and RNc enabled an unprecedented level of anatomical precision. Histological reference data from E.J.L. Alho; fibre dissection data from V. Milanese Holanda. *Abbreviations*: ac, anterior commissure; al, ansa lenticularis; as, ansa subthalamica; DN, dentate nucleus; fx, fornix; GPe, globus pallidus externus; GPi, globus pallidus internus; ic, internal capsule; lml, lateral medullary lamina; MB, mammillary body; mml, medial medullary lamina; mtt, mammillothalamic tract; opt, optic tract; RN, red nucleus; RNc, capsule of the red nucleus; STN, subthalamic nucleus.

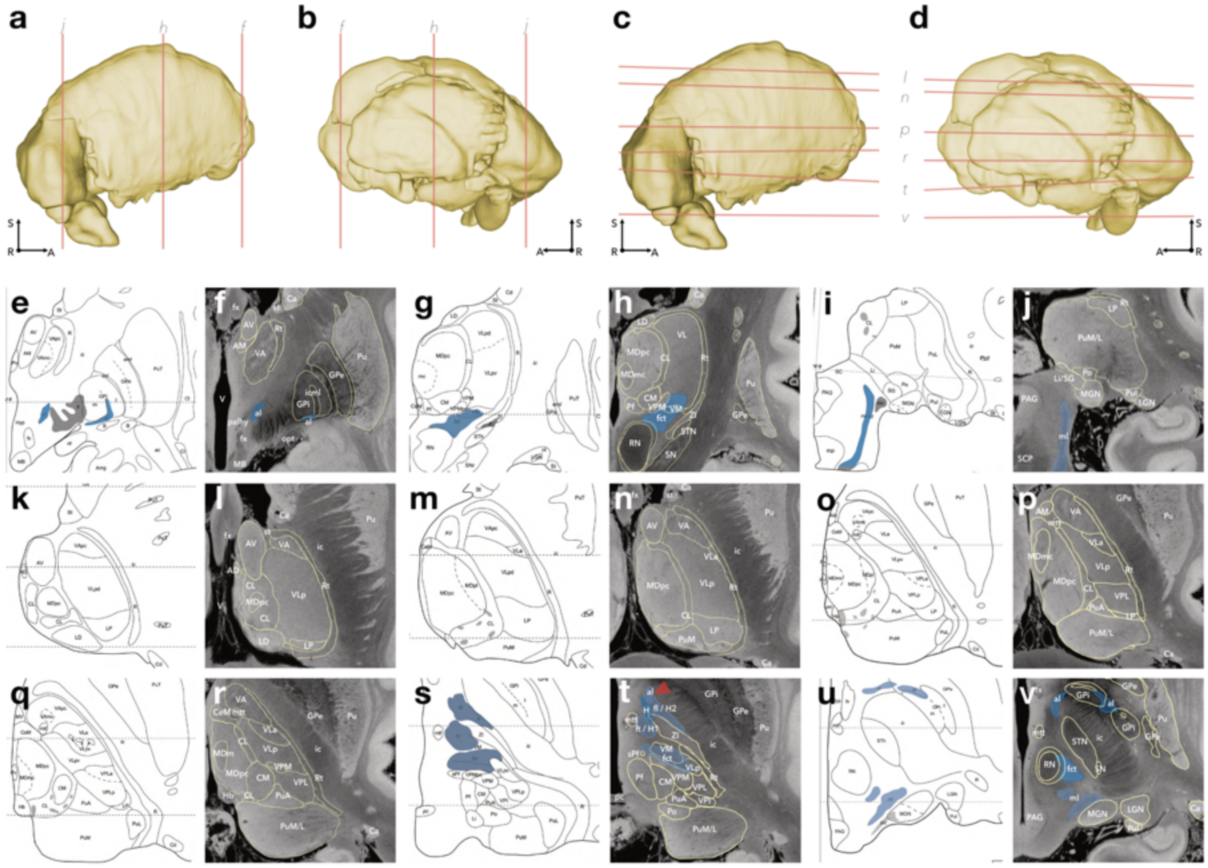

***Supplementary Figure 2.*** **Delineation of thalamic nuclei. a**–**d** Illustration of section planes corresponding to panels f, h, j, l, n, p, r, t, v. **e** Coronal section at the level of the mammillary body and anterior thalamus (adapted from [1]). **f** Corresponding 7T MRI section showing strong concordance in the delineation of nuclei and white-matter structures (e.g., ansa lenticularis, al) between [1] and the present study. **g**–**h** Similar agreement at more posterior levels near the red nucleus. **i**–**j**, Consistency extends to the level of the superior cerebellar peduncle (SCP). Please note, that the differentiation between the limitans nucleus (Li) and the suprageniculate nucleus (SG), as well as between medial and lateral pulvinar, was not feasible in the present study. **k** Axial section at the level of the anteroventral nucleus (AV) in a superior portion of the thalamus ([1]). **l** corresponding 7T MRI section showing close agreement in nuclear boundaries. **m**–**p** Similar concordance in more inferior thalamic sections. **q**–**v** Progressively more caudal sections through the thalamus and upper mesencephalon, with white-matter tracts highlighted in blue for clarity (adapted from [1]). The anatomy at the mesencephalic–diencephalic junction is particularly complex, and two-dimensional histological atlases often leave blind spots due to slicing-plane limitations. By integrating multiple reference atlases (details see *Supplementary Notes*) and segmenting within a three-dimensional volume, this study overcomes these constraints. Definitions of thalamic nuclei and fibre tracts closely align with [1], with even finer delineation of pathways such as the confluence of the ansa lenticularis (al) into the field of Forel H2 / fasciculus lenticularis (fl, red arrowhead) and the tegmental fields ("Haubenfelder") first described by [2]. For further details on the fields of Forel, see *Figure 5* and the detailed documentation available at the referenced *data repository*. Illustrations in e, g, I, s and u include blue highlights of selected white-matter tracts for improved visualization. *Abbreviations:* al, ansa lenticularis; AM, anteromedial nucleus; AV, anteroventral nucleus; Ca, caudate nucleus; CeM, central medial nucleus; CL, central lateral nucleus; CM, centromedian nucleus; fct, fasciculus cerebellothalamicus; ft, fasciculus thalamicus; fx, fornix; GPe, external segment of globus pallidus; GPi, internal segment of globus pallidus; H, field of Forel H; H1, field of Forel H1; H2, field of Forel H2; Hb, habenula; ic, internal capsule; icml, incomplete medullary lamina; LD, lateral dorsal nucleus; LGN, lateral geniculate nucleus; Li/SG, limitans and suprageniculate nuclei; Lp, lateral posterior nucleus; MB, mammillary body; MDm, mediodorsal nucleus, magnocellular division; MDpc, mediodorsal nucleus, parvocellular division; MGN, medial geniculate nucleus; ml, medial lemniscus; mtt, mammillothalamic tract; PAG, periaqueductal gray substance; palhy, pallidohypothalamic tract; pc, posterior commissure; Pf, parafascicular nucleus; Po, posterior nucleus; Pu, putamen; PuA, anterior pulvinar nucleus; PuI, inferior pulvinar nucleus; PuM/L, medial and lateral pulvinar nuclei; Rt, reticular nucleus; RN, red nucleus; SCP, superior cerebellar peduncle; SN, substantia nigra; STN, subthalamic nucleus; st, stria terminalis; V, ventricle; VA, ventral anterior nucleus; VLa, ventral lateral anterior nucleus; VLp, ventral lateral posterior nucleus; VM, ventral medial nucleus; VPI, ventral posterior inferior nucleus; VPL, ventral posterior lateral nucleus; VPM, ventral posterior medial nucleus; ZI, zona incerta. Permissions to reproduce panels that were adapted from original work (as indicated above) are on file.

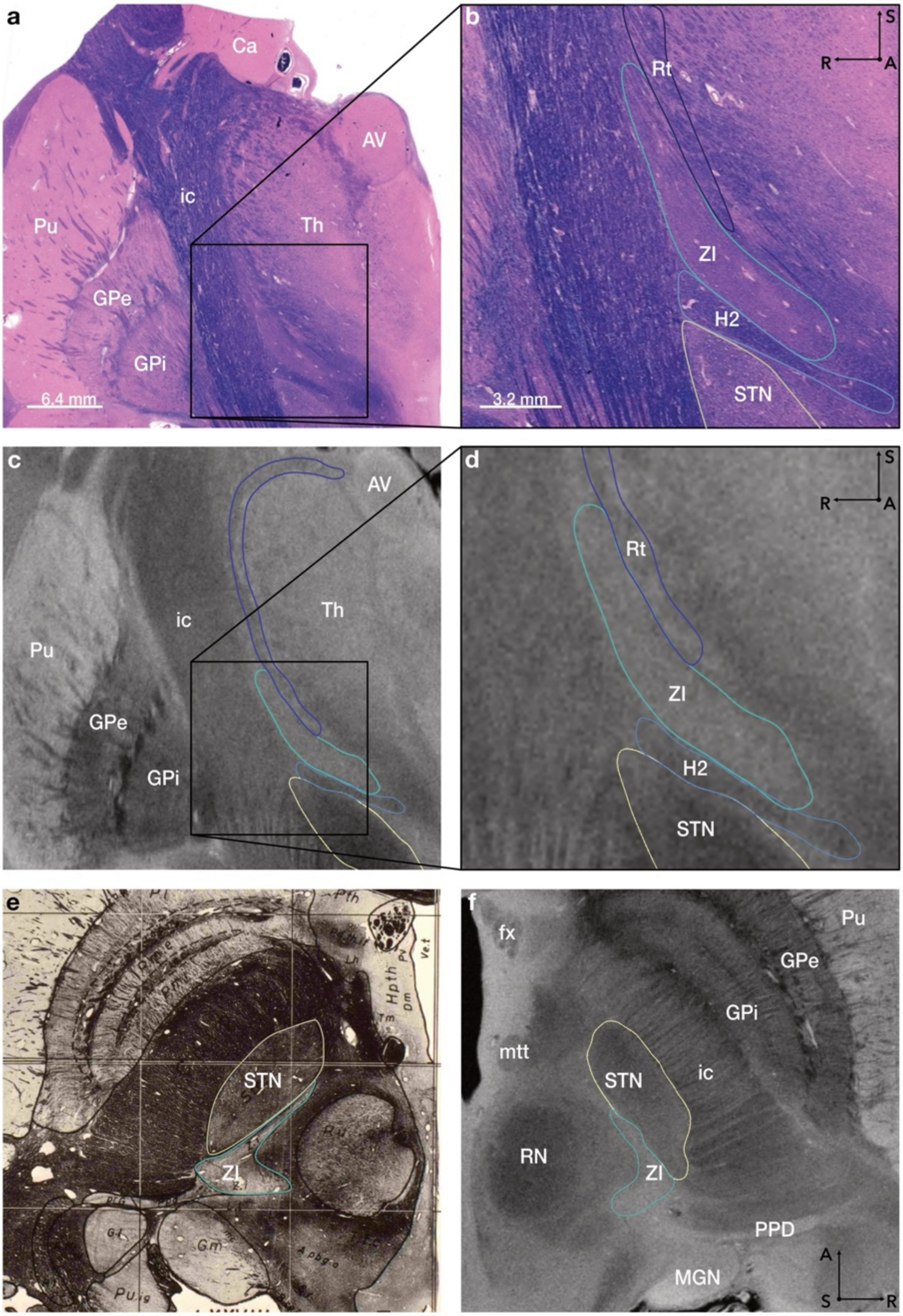

***Supplementary Figure 3.*** **Integrative delineation workflow for the zona incerta (ZI) and reticular nucleus (Rt).** This figure illustrates the multi-modal delineation approach combining MRI, histology, and atlas references to define subthalamic structures with high precision. The ZI and Rt were delineated with reference to [1,3–7] and refined using histological preparations kindly provided by Brian L. Edlow ([8]), as shown in **a** (magnified in **b**). **c**–**d** Corresponding 7T MRI sections depict the ZI as a hyperintense band adjacent to the more hypointense Rt. Histological comparison guided the definition of their shared boundary and ensured consistent segmentation across planes. While most atlases leave this interface ambiguous, the 7T scan reveals distinct signal contrasts corroborated by histology. As shown in **a**–**d**, caudal portions of the Rt extend medially to the ZI, visible in both modalities. **e** Axial section through the caudal ZI (adapted from [1]); **f** corresponding 7T MRI with delineated ZI boundaries. *Abbreviations*: AV, anteroventral nucleus; Ca, caudate nucleus; fx, Fornix; GPe, external segment of globus pallidus; GPi, internal segment of globus pallidus; H2, field of Forel H2; ic, internal capsule; MGN, medial geniculate nucleus; mtt, mammillothalamic tract; PPD, peripeduncular nucleus; Pu, putamen; RN, red nucleus; Rt, reticular nucleus; STN, subthalamic nucleus; Th, thalamus; ZI, zona incerta. Permissions to reproduce panels that were adapted from original work (as indicated above) are on file.

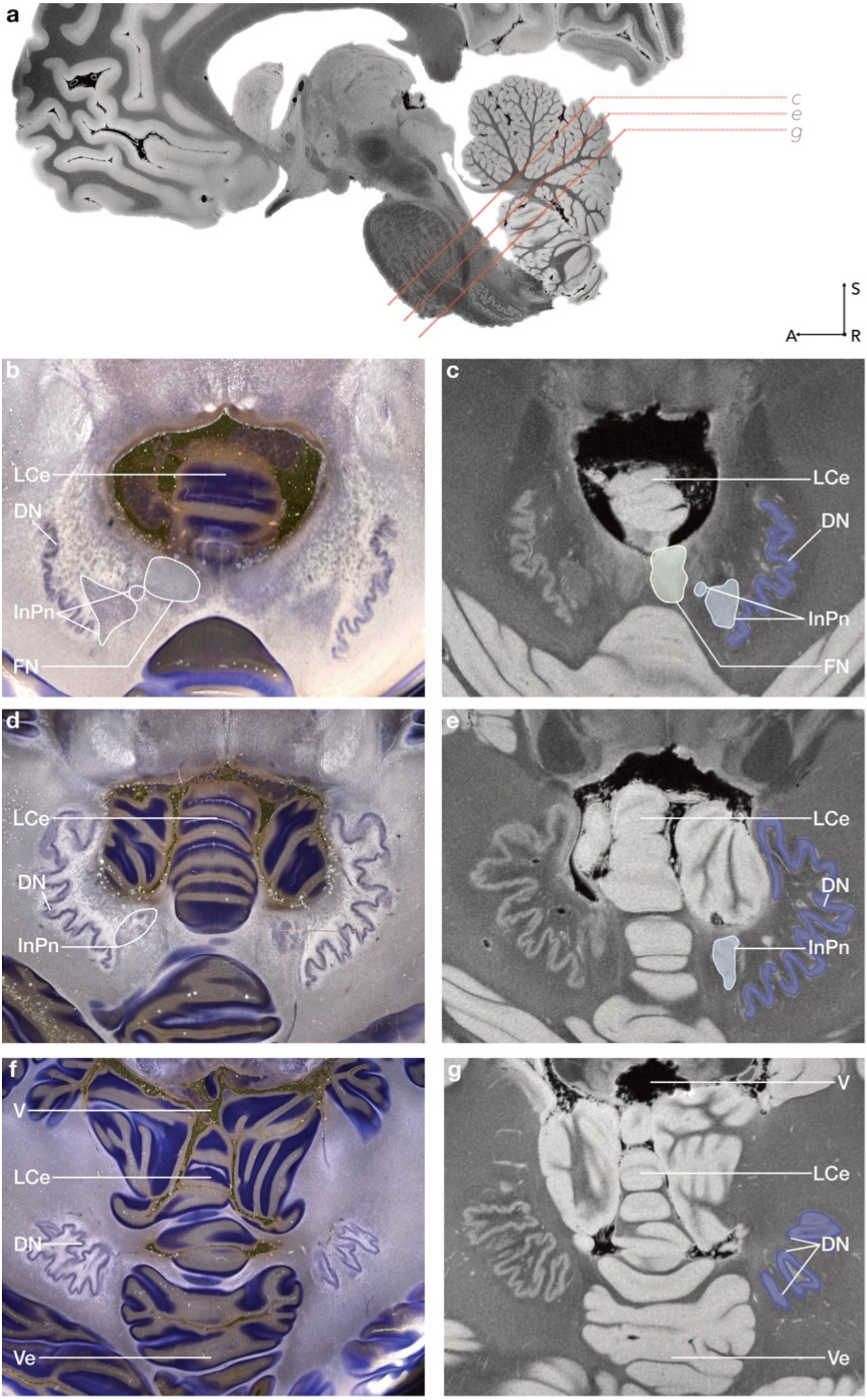
a
c
e
g
S
A
R
b
LCe
DN
InPn
FN
c
LCe
DN
InPn
FN
d
LCe
DN
InPn
e
LCe
DN
InPn
f
V
LCe
DN
Ve
g
V
LCe
DN
Ve

***Supplementary Figure 4.*** **Integration of MRI and histology for delineation of deep cerebellar nuclei**. **a** Sagittal section (left view) illustrating slice planes (red) corresponding to panels **c**–**g** in the 7T MRI, aligned with dark-field microscopy data. The dentate nucleus (DN), interposed nuclei (InPn), and fastigial nucleus (FN) were segmented based on [9,10],with additional guidance from histological preparations kindly provided by Eduardo J. Alho. Dark-field microscopy (examples in b, d, f) provided exceptional clarity for defining nuclear boundaries, which informed segmentation in the 7T MRI. In the MRI template, the deep cerebellar nuclei appear as hyperintense structures embedded within the hypointense medulla cerebelli. Panels b–g are displayed in histological convention. *Abbreviations*: DN, dentate nucleus; FN, fastigial nucleus; InPn, interposed nuclei; LCe, lingula cerebelli; Ve, vermis cerebelli; V, ventricle.

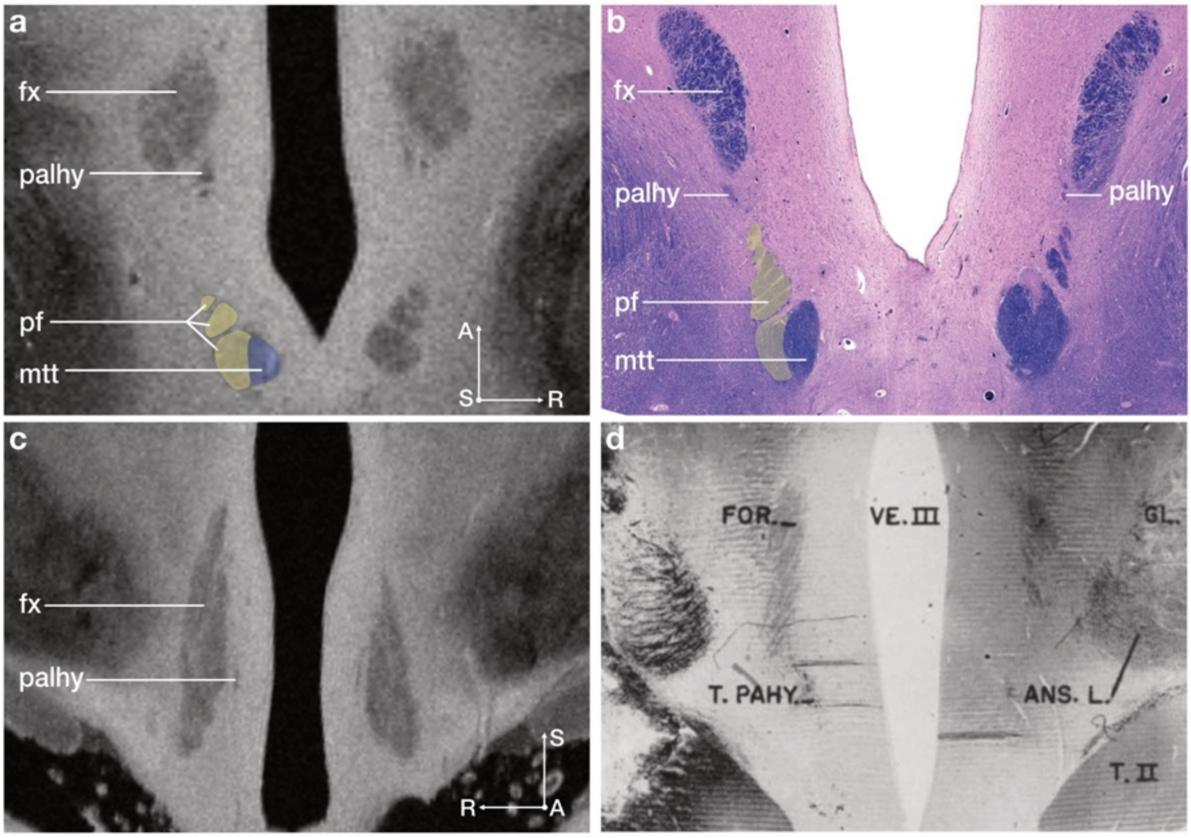

***Supplementary Figure 5.*** **High-resolution reconstruction of fine pallidofugal and hypothalamic fibre pathways.** This figure illustrates how the applied multimodal workflow enables accurate 3D reconstruction of small and anatomically complex white-matter tracts. Axial sections through the upper midbrain show distinct fibre bundles near the fornix (fx) in the **a** 7 T MRI, corresponding to the pallidohypothalamic tract (palhy). **b** Corresponding histological section (H.E. stain). The palhy diverges from pallidofugal fibres within the fields of Forel and proceeds ventromedially toward the hypothalamus, consistent with early histological descriptions in animal material ([11]). The histological section in b is slightly oblique, causing the left palhy to appear lateral to the fornix, accurately reflecting its anatomical course. In the present work, careful inspection of axial sections revealed distinct fibre bundles corresponding to the bifurcation of the principal fasciculus (pf, displayed in yellow in a & b) into the mammillothalamic tract (mtt, displayed in purple in a & b) and the mammillotegmental tract (mtg) of Gudden ([12]). While the latter could not be traced beyond its proximal origin in the available MRI specimen, the bifurcation at the level of the mammillary body (MB) could be delineated with high anatomical precision. This detailed correspondence between MRI and histology underscores the resolving power of the present 3D segmentation approach, allowing differentiation of pathways that have been subsumed under a single mtt label in most existing atlases. **c** illustrates the medial course of the palhy toward the fornix (fx), consistent with the trajectory observed in non-human primate preparations in **d** by [13]. c-d displays a more anterior coronal section, demonstrating the continuation of the palhy trajectory toward the hypothalamus. Given that the fornix is being explored as a potential DBS target for cognitive modulation (e.g.[14]), a detailed understanding of the surrounding anatomy including neighbouring fibre systems such as the pallidohypothalamic tract and fields of Forel may hold clinical relevance for future interventions in this region. *Abbreviations*: fx, fornix; MB, mammillary body; mtt, mammillothalamic tract; palhy, pallidohypothalamic tract; pf, principal fasciculus; Th, Thalamus. Permissions to reproduce panels that were adapted from original work (as indicated above) are on file.

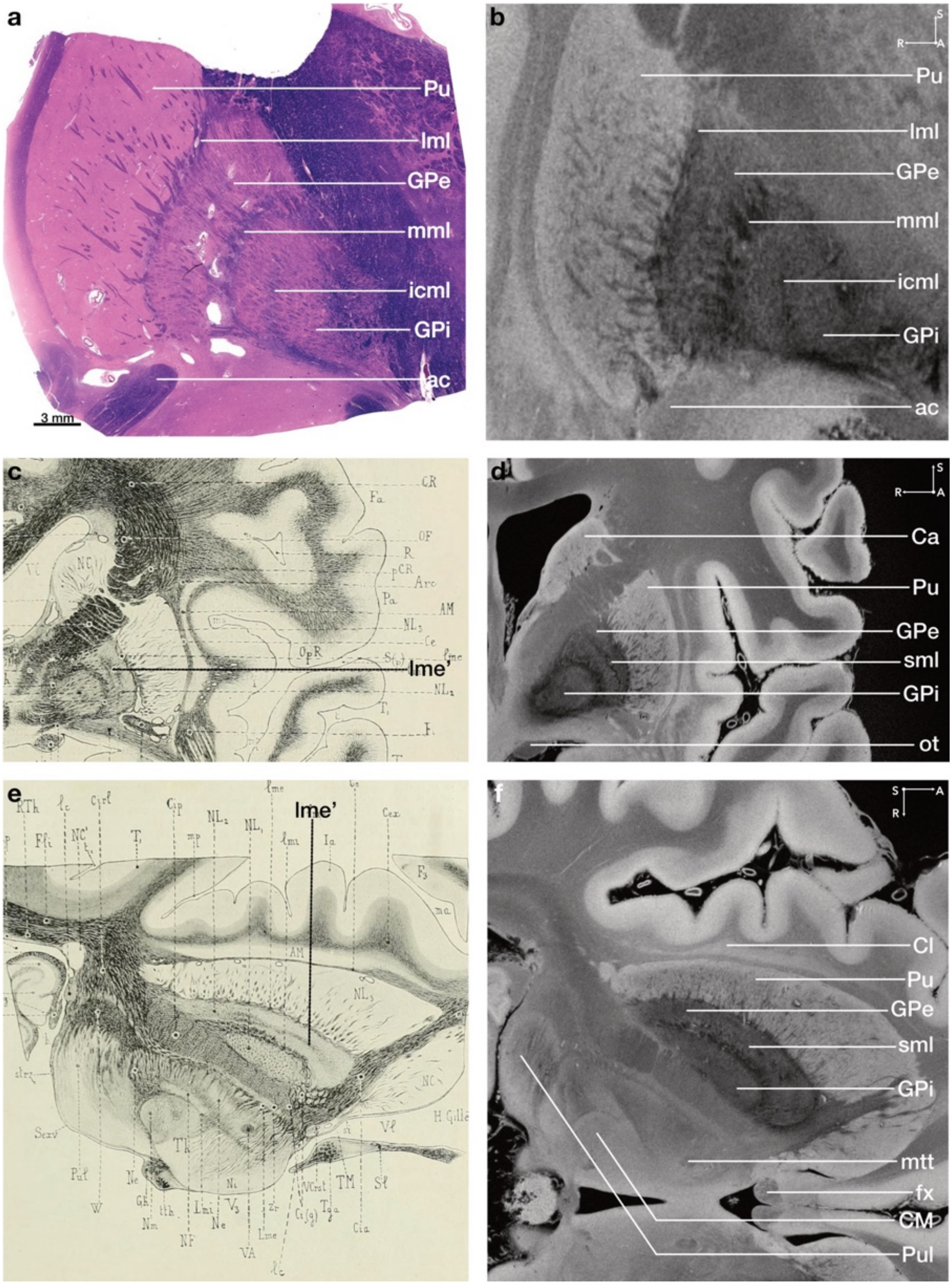

***Supplementary Figure 6.*** **Integration of histological and historical references for delineating fine pallidal laminae. a** Histological section stained with H.E. in a coronally oriented plane (kindly provided by B.L. Edlow,[8]). The medullary laminae are clearly distinguishable and can also be identified as hypointense structures in the corresponding MRI section shown in **b**. **c** Historical anatomical drawing by [15] in a coronal view, depicting the supplementary medullary lamina (french: lame médullaire supplémentaire du deuxième segment du noyau lenticulaire; lme'), **d** which we were able to delineate in the corresponding MRI section. **e**-**f** Comparison of the supplementary medullary lamina within the external segment of the globus pallidus, shown in an obliquely angulated axial section. *Abbreviations*: ac, anterior commissure; Ca, caudate nucleus; Cl, claustrum; CM, centromedian nucleus; fx, fornix; GPe, external segment of the globus pallidus; GPi, internal segment of the globus pallidus; icml, incomplete medullary lamina; lml, lateral medullary lamina; mml, medial medullary lamina; mtt, mammillothalamic tract; ot, optical tract; Pu, putamen; Pul, lateral pulvinar; sml/Lme', supplementary medullary lamina. Permissions to reproduce panels that were adapted from original work (as indicated above) are on file.

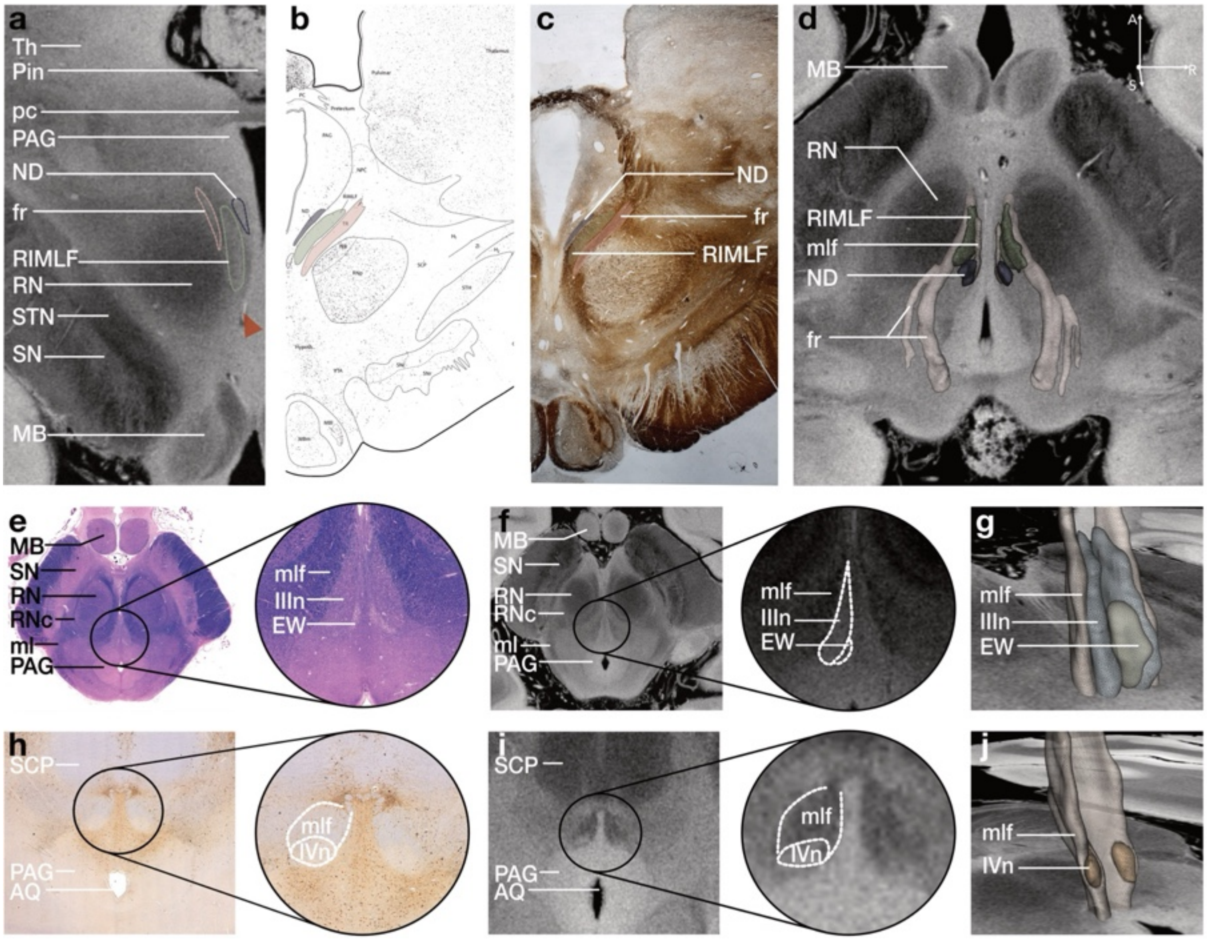

***Supplementary Figure 7.*** **Submillimetric delineation of ocular motor nuclei in the human midbrain. a** Fine signal contrasts in the rostral midbrain at 7T MRI allow delineation of key ocular motor nuclei. The arteria subthalamica posterior (red arrowhead) and fasciculus retroflexus (fr, rose colour) serve as reliable anatomical landmarks (see [16,17] ) for identifying the rostral interstitial nucleus of the medial longitudinal fasciculus (RIMLF, green colour). **b**–**c** In all three planes, these landmarks further aid the definition of the nucleus of Darkschewitsch (ND, blue colour), guided by histological reference sections kindly provided by A. K. E. Horn and corresponding atlas information. **d** Three-dimensional reconstruction of these nuclei, located medial to the red nucleus (RN). **e** In H.E.-stained histological sections, the medial longitudinal fasciculus (mlf), oculomotor nucleus (IIIn), and Edinger–Westphal nucleus (EW) can be clearly distinguished. **f** In the corresponding 7T MRI, the mlf appears as a hypointense structure, the IIIn as a relatively hyperintense cluster, and the EW as a more hypointense group dorsomedial to the IIIn. **g** Three-dimensional reconstruction of the oculomotor complex. **h** Immunohistochemistry (TH) was chosen as it provides clear delineation of dopaminergic midbrain nuclei and adjacent non-dopaminergic structures, such as the trochlear nucleus (IVn) and mlf, which appear as TH-negative regions within a densely stained background. **i** In the 7T MRI, this trochlear nucleus corresponds to a discrete hyperintense interruption within the hypointense mlf. **j** Three-dimensional reconstruction at the level of the midbrain tegmentum and inferior colliculi, illustrating the relative positions of the IIIn, EW, IVn, and RIMLF. *Abbreviations*: EW, Edinger–Westphal nucleus; fr, fasciculus retroflexus; IIIn, oculomotor nucleus; IVn, trochlear nucleus; MB, mammillary body; ml, medial lemniscus; mlf, medial longitudinal fasciculus; PAG, periaqueductal gray; pc, posterior commissure; Pin, pineal body; RN, red nucleus; RNc, capsule of the red nucleus; SCP, superior cerebellar peduncle; SN, substantia nigra; STN, subthalamic nucleus; Th, thalamus; RIMLF, rostral interstitial nucleus of the medial longitudinal fasciculus. Permissions to reproduce panels that were adapted from original work (as indicated above) are on file.

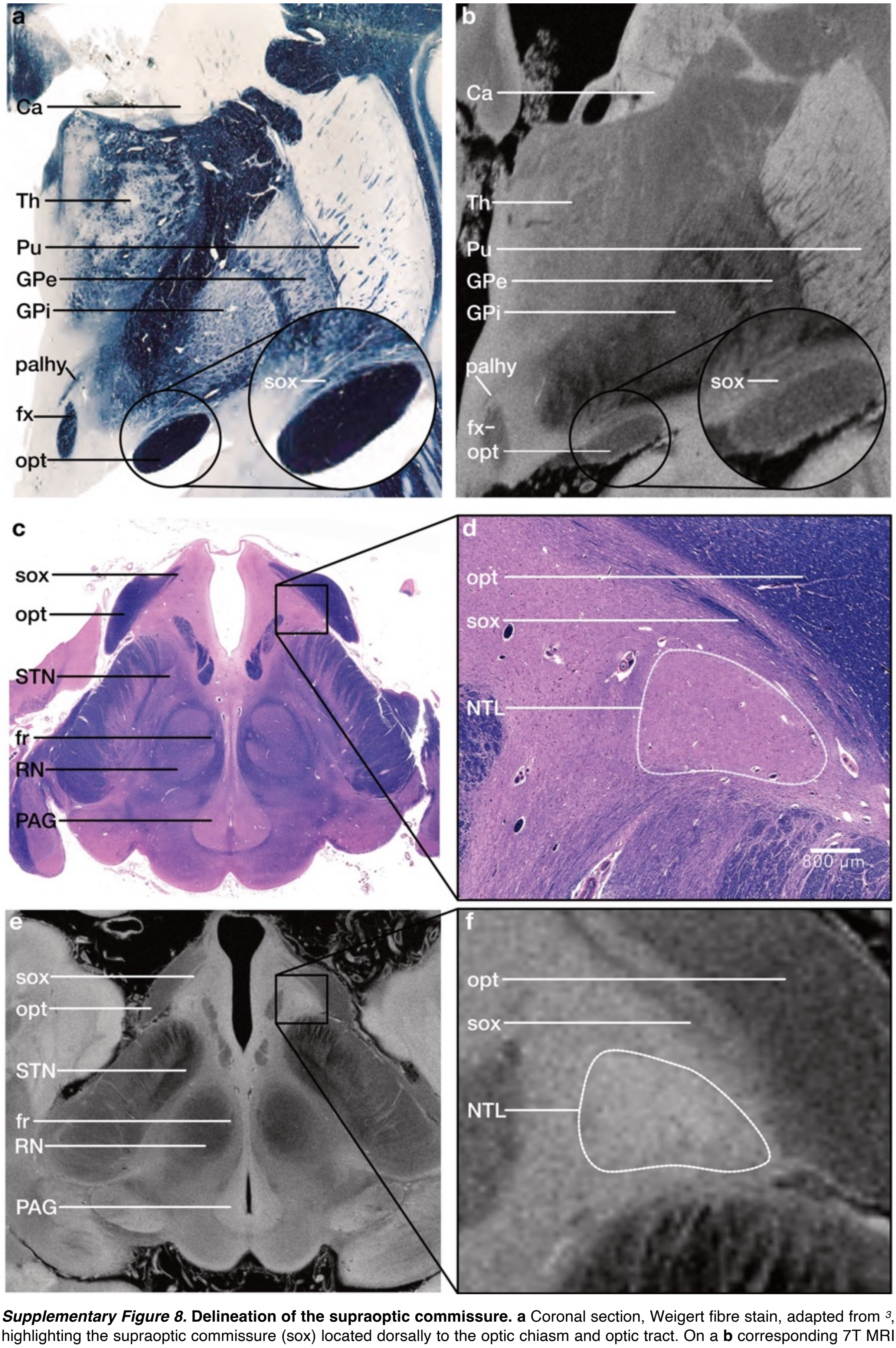

***Supplementary Figure 8.*** **Delineation of the supraoptic commissure. a** Coronal section, Weigert fibre stain, adapted from [3], highlighting the supraoptic commissure (sox) located dorsally to the optic chiasm and optic tract. On a **b** corresponding 7T MRI slice, the sox is visible as a hypointense structure. Delineation of the sox was performed according to [3] and with reference to histological preparations kindly provided by B. L. Edlow ([8]). **c** Axial histological section (H.E. stain) with **d** a magnified view of the region of interest. **e**-**f** High-resolution template representations further support the identification of the sox, which measures a mere 0.5 mm in diameter. The sox contains thin fibre bundles that cross the midline ([18]): The ventral supraoptic commissure of Gudden, the anterior hypothalamic commissure of Ganser and the dorsal supraoptic commissure of Meynert (which interconnects the subthalamic nuclei, the globus pallidus, the lateral geniculate bodies, and the SC ([19]). Additionally, the lateral tuberal nucleus (NTL,[15,20–22]), appearing as a hyperintense signal, could be identified. *Abbreviations*: Ca, caudate nucleus; fx, fornix; fr, fasciculus retroflexus; GPe, external segment of the globus pallidus; GPi, internal segment of the globus pallidus; NTL, lateral tuberal nucleus; opt, optic tract; PAG, periaqueductal gray; palhy, pallidohypothalamic tract; Pu, putamen; RN, red nucleus; sox, supraoptic commissure; Th, thalamus. Permissions to reproduce panels that were adapted from original work (as indicated above) are on file.

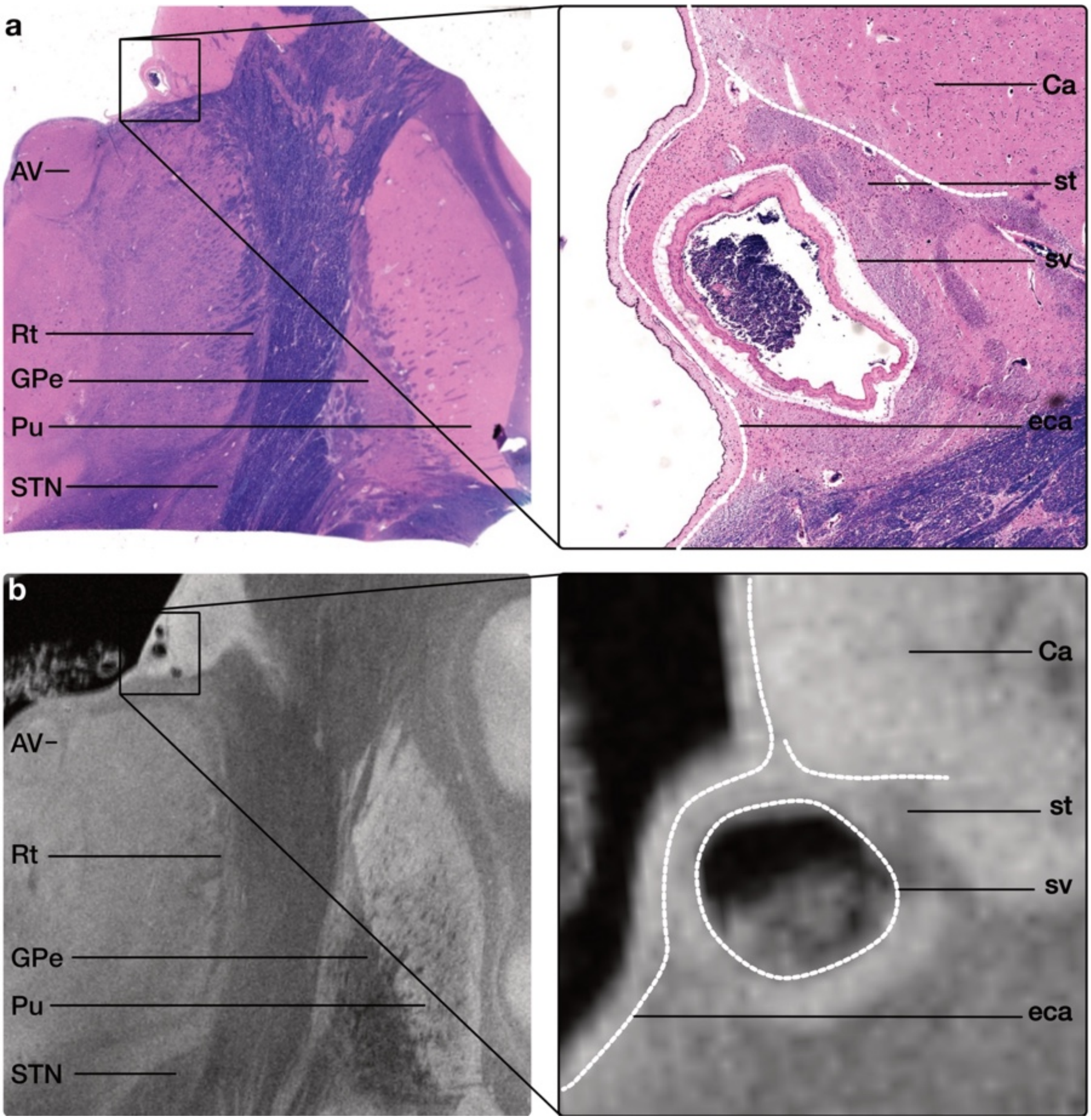

***Supplementary Figure 9.*** **Anatomical delineation of the caudate nucleus. a** Histological section illustrating the relationship between the caudate nucleus, the ependymal lining of the lateral ventricle (eca), the adjacent stria terminalis (st), and the terminal vein (sv). The terminal vein appears artificially shrunken as a consequence of tissue processing. Histological preparations kindly provided by B. L. Edlow ([8]). **b** Corresponding high-resolution MRI section. The caudate nucleus is readily identifiable by its relatively hyperintense signal, while the adjacent stria terminalis appears comparatively hypointense. Notably, the ependymal lining and terminal vein are also clearly resolved on MRI, providing additional anatomical landmarks for precise delineation of the ventricular and inferomedial boundary of the caudate nucleus.

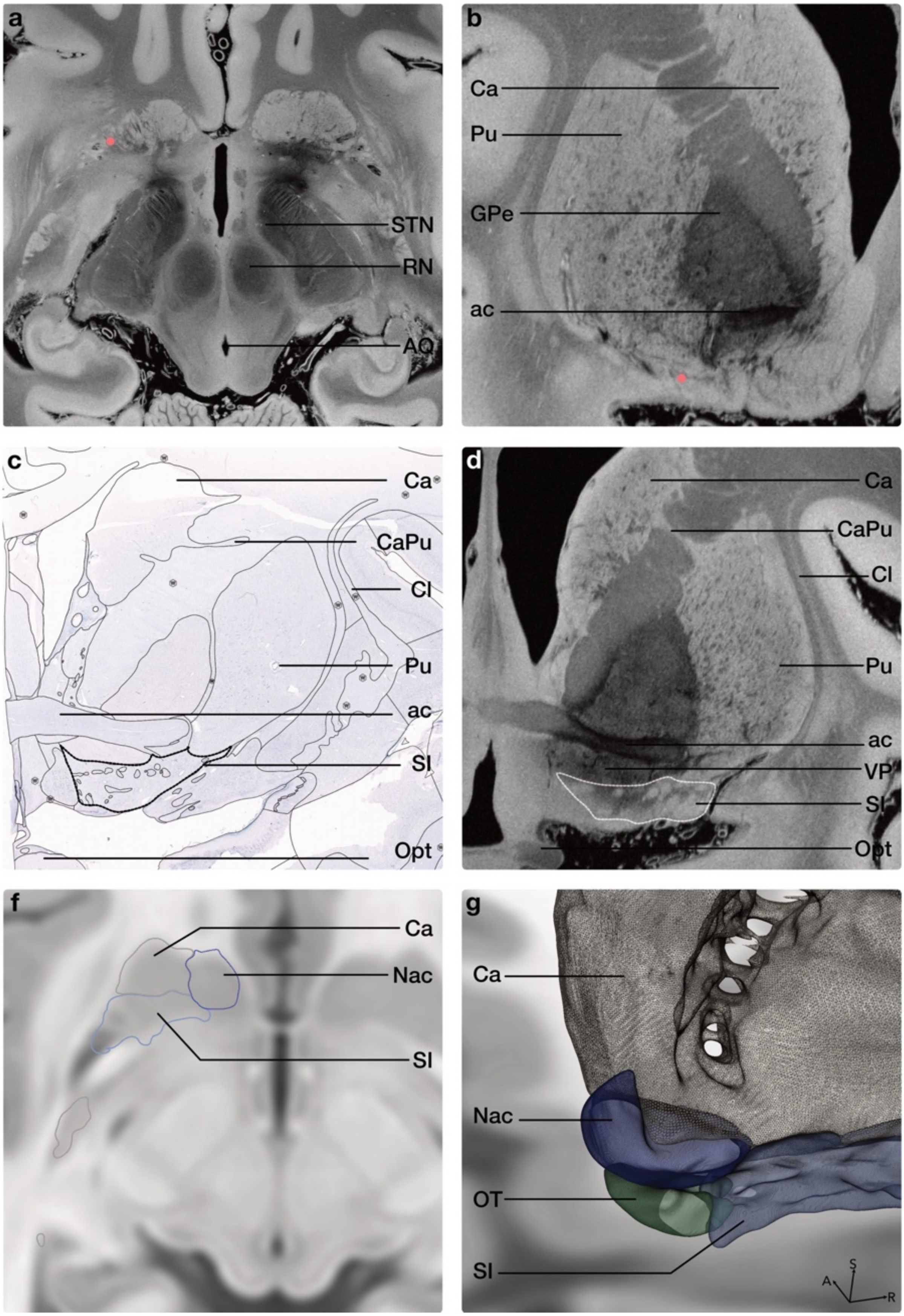

***Supplementary Figure 10.*** **Multiplanar delineation of the ventral striatum and substantia innominata. a** In the axial plane, the ventral striatal region appears as a relatively diffuse hyperintense signal, with individual anatomical boundaries not readily discernible. This region is likewise only incompletely resolved in many conventional anatomical atlases, particularly in axial representations. **b**-**d** By identifying the corresponding anatomical location across imaging planes (red points) and tracing it in the coronal plane, where the complex anatomy of the ventral striatum and basal forebrain is more clearly represented in established histological atlases and literature ([3,9,23–27]; c coronal histological section by [9]), the boundaries of the caudate nucleus (Ca), putamen (Pu), nucleus accumbens (Nac), olfactory tubercle (OT), and substantia innominata (SI) could be reconstructed in a stepwise manner. These initial coronal delineations were then iteratively refined across axial and sagittal planes, allowing the substantia innominata to be carefully separated from the adjacent putamen and other ventral striatal structures even where local MRI contrast alone was insufficient for an unambiguous boundary assignment. **e**-**f** The resulting segmentations were subsequently transformed into MNI space using careful registration, thereby transferring anatomical detail established in the high-resolution native dataset into standard space, where these boundaries would not be directly resolvable from image contrast alone. **g** Three-dimensional reconstruction illustrating the resulting spatial relationships between the nucleus accumbens, olfactory tubercle, substantia innominata, and striatum.

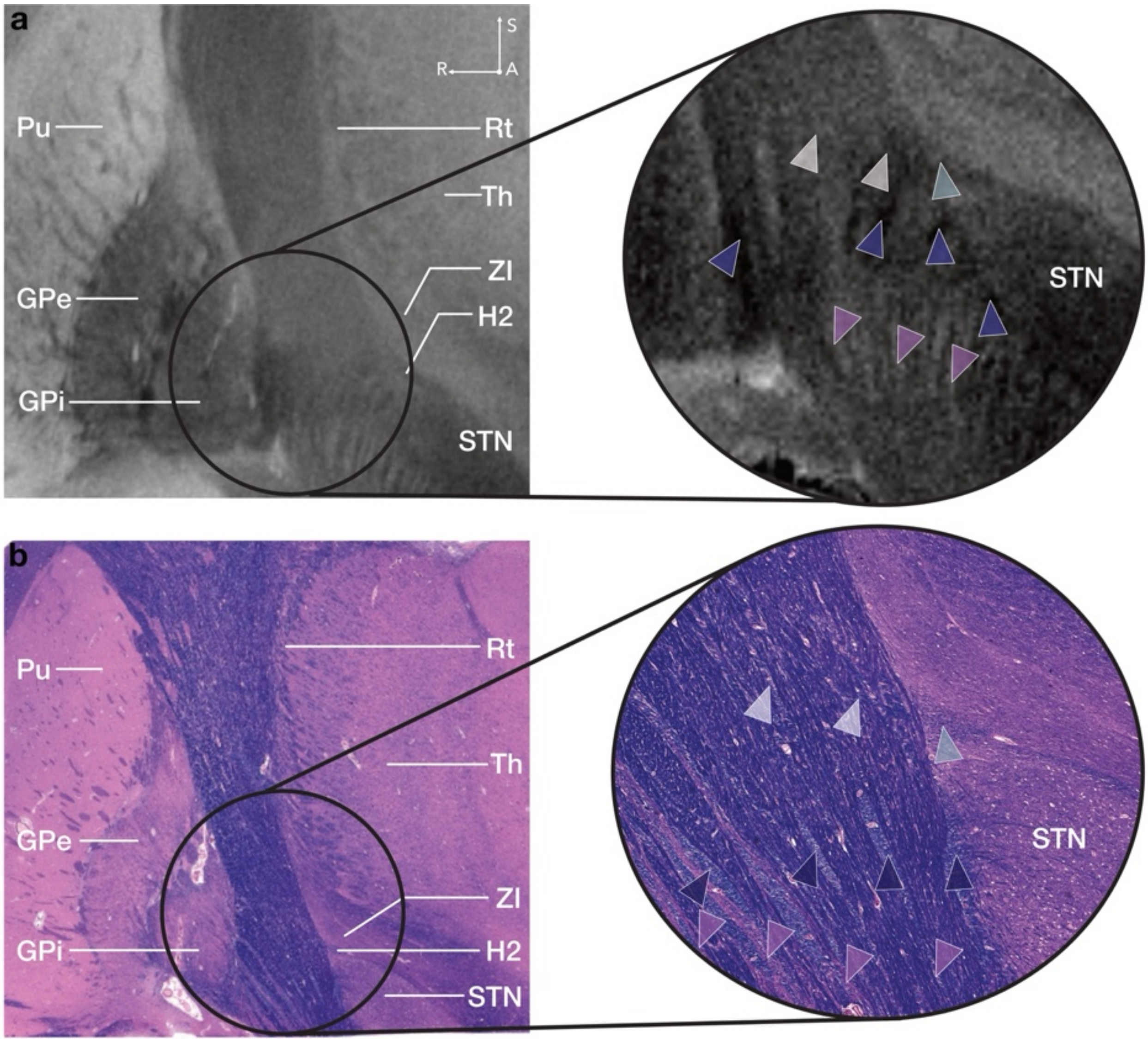

***Supplementary Figure 11*. Histology-guided delineation of rich MRI signal patterns in the cerebral peduncle. a** Coronal section of the 7T MRI template illustrating the complex internal signal architecture of the cerebral peduncular and adjacent pallidal region. The rich signal architecture of the 7T MRI template provides substantial anatomical detail that can be resolved through careful, histology-informed visual inspection, including fine-grained features that may remain challenging for automated segmentation. **b** Corresponding coronal histological section of a human brain specimen (H.-E.), adapted from [8]. Purple arrows indicate striatopallidal fibres forming part of the Edinger comb system ([26]), whereas white arrows indicate fibre bundles originating from the region of field H2 of Forel (turquoise arrow), as well as portions of the fasciculus lenticularis. These fibre-rich structures exhibit relatively hypointense signal on MRI and show close spatial correspondence with their histological counterparts, indicated by arrows of matching colour. In contrast, the fields of Sano A, representing islands and bridges of gray matter along the continuum between the substantia nigra pars reticulata (SNr) and globus pallidus internus (GPi; [28,29]), appear relatively hyperintense on the 7T MRI. Their corresponding histological location is indicated by pink arrows.

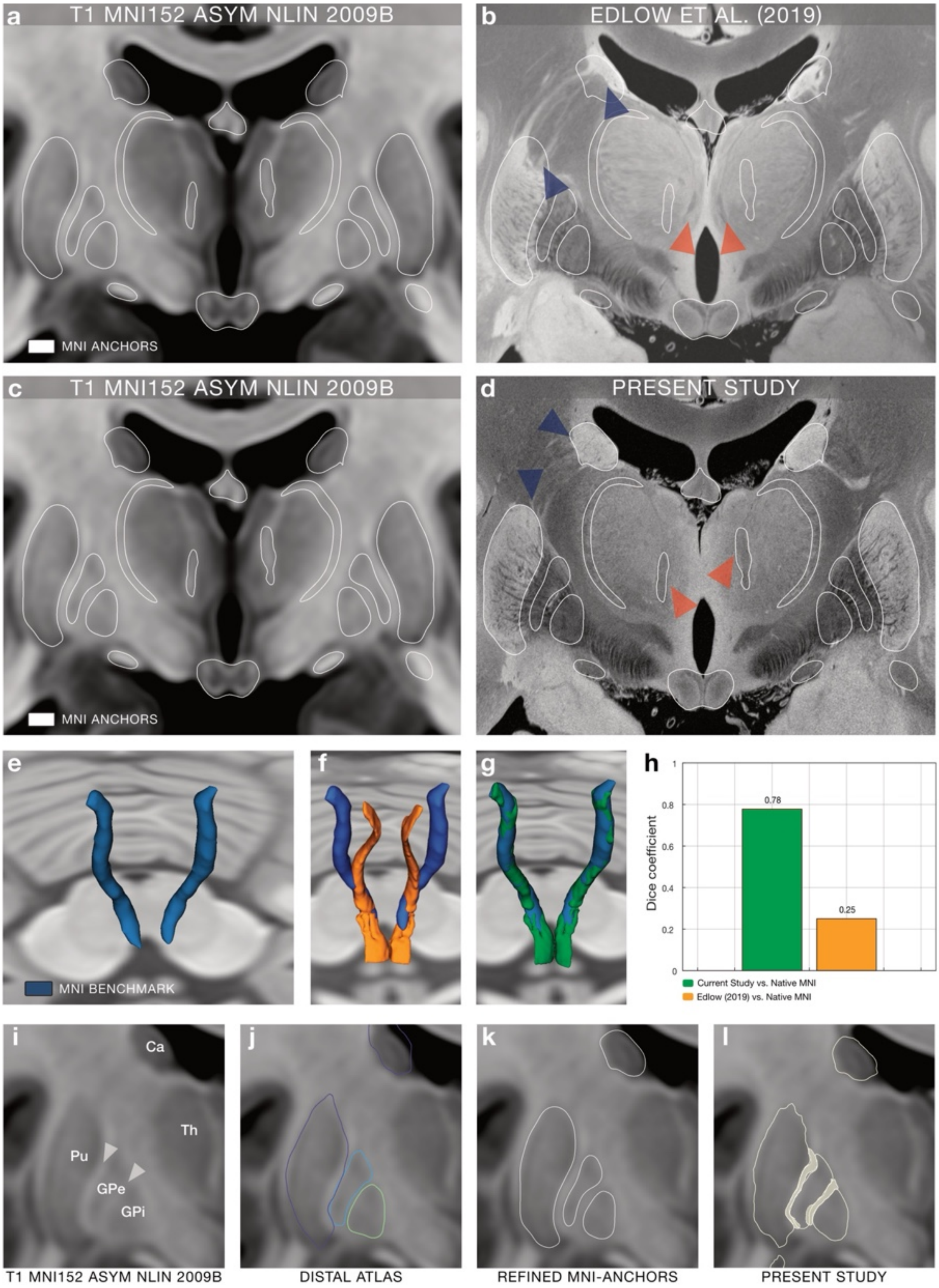

***Supplementary Figure 12.*** **Example of enhanced alignment achieved through refinement of native-MNI space transformation. a** & **c** coronal section at the level of the mammillary bodies in original T1 MNI space. MNI ground truth anchor structures (white borders), were selected based on reliably identifiable boundaries in native contrast and manually delineated bilaterally in this original MNI space. **b** Upon projection, the fit between the ground truth MNI anchor structures and their corresponding structures in the MNI version of the high-resolution 100-micron MRI as of 2019 shows suboptimal overlap (e. g. medial deviation of the mtt (red arrows), or misalignment at the boundaries of the putamen/caudate (blue arrows)), highlighting the need for an improved transformation in this subcortical region, encompassing important clinical intervention zones such as thalamus, subthalamus, and the mesencephalic-diencephalic transition zone. **d** Through an optimized registration process involving 62 corresponding anchor segmentations (31 per hemisphere), readily identifiable in both MNI and native contrast of the high-resolution template, and 37 iterations (for details see referenced *data repository*), the refined transformation demonstrates a stronger anatomical correspondence, with aforementioned deviations being clearly minimized (red and blue arrows). **e** Blue: Manually segmented mtt in MNI space ("native MNI"), defined directly in the T1-weighted MNI template and used as ground truth. **f** Overlay of the mtt segmentation transformed from 7T native space using the method by [30]. **g** Overlay of the same native mtt segmentation transformed using the updated pipeline developed in the current study. **h** Quantitative comparison of the two transformations. The Dice coefficients (current study: 0.78, Edlow et al.: 0.25) indicate a more than threefold improvement in anatomical overlap. It is important to note that ventral portions of the tract, namely the principal fasciculus (prf), from which the mtt emerges, are not captured in the T1-based ground truth segmentation due to limited visibility. This potentially leads to an underestimation of both Dice and precision scores. Nevertheless, the consistent and substantial gain across all metrics clearly reflects the improved anatomical accuracy of the updated normalization pipeline. For precise coregistration of high-resolution segmentations, it was crucial to ensure that corresponding segmentations were present in the reference space. This step was necessary to guarantee that native structures without an anchor structure, were correctly positioned relative to neighbouring structures (see also *Figures 2 & 3*). As an example, **i** shows a coronal slice of the T1 MNI template. Hypointense structures such as Ca, Pu, GPe, and GPi are clearly visible. Between the Pu, GPe, and GPi, medullary laminae (mml, lml; see *Supplementary Figure 6*) can be partially distinguished as hyperintense structures in the T1 image (indicated by white arrowheads). Many atlases, such as the DISTAL Atlas shown in **j**, do not provide strict boundaries for these laminae, necessitating **k** manual re-delineation of these structures in MNI space. **l** demonstrates that the Pu, GPe, GPi, and Ca are well-registered and now serve as anchor elements. The medullary laminae with their overlapping nature are correctly positioned. In contrast to the MNI anchor segmentations, the high-resolution atlas reveals finer anatomical detail, yielding a less smoothed but more precise and faithful representation of the structures' morphology. *Abbreviations*: Ca, caudate nucleus; GPe, external segment of the globus pallidus; GPi, internal segment of the globus pallidus; Pu, putamen; Th, thalamus.

### Supplementary Notes

*Introduction*

This work had four main objectives, each forming a cornerstone of the high-resolution atlas framework. First, we delineated subcortical structures in a 100 µm isotropic 7 T MRI dataset. These segmentations provided the anatomical foundation for subsequent tract reconstructions in MNI space, serving as reference points alongside established anatomical resources and literature. To assess the reproducibility of the manual delineations, we additionally performed an inter-rater agreement analysis on a representative subset of native-space segmentations (see *Supplementary Section 1*). Second, to achieve anatomically precise normalization, we developed a dedicated transformation protocol and accompanying software optimized for subcortical alignment. This customized warp yielded markedly improved correspondence within the thalamus, the mesencephalo-diencephalic transition, and the mesencephalon compared with the original 2019 normalization framework (see *Supplementary Section 2*). We further quantified spatial agreement for the anatomical anchor structures, characterized the iterative refinement process and local transformations, and provide a step-by-step protocol to facilitate reproducible application of the workflow (see *Supplementary Section 2*). Third, building directly on this refined transformation, we generated an updated 100 µm MNI template with enhanced subcortical alignment (see *Methods*). We further assessed cross-atlas variability of subcortical definitions in MNI space, both qualitatively and quantitatively, to contextualize the importance of anatomical delineation and spatial normalization (see *Supplementary Section 3*). Fourth, leveraging the high-resolution segmentations, we manually reconstructed major subcortical fibre tracts in MNI space. Finally, we demonstrated selected illustrative clinical applications of the atlas (see *Results*) and examined its applicability across individual healthy and pathophysiological brain anatomies (see *Supplementary Section 4*).

1. Manual delineation of subcortical brain structures and supporting anatomical resources

The first step of this work was the precise delineation of subcortical structures using a 100 µm isotropic 7 T MRI dataset by [30]. A total of 180 grey and white matter structures were manually segmented across both hemispheres, corresponding to 85 bilateral and 10 midline anatomical structures (see *Supplementary Table 1*). To ensure anatomical fidelity at submillimeter precision, each boundary was iteratively refined under expert supervision using multiple contrast angles and orthogonal slice views.
To accurately capture clinically relevant yet often underrepresented regions, such as the Fields of Forel, ventral tegmental area, and the mesencephalo-diencephalic transition zone, including anatomically complex areas with challenging MRI contrast, we employed a multimodal comparative approach. This involved systematic cross-referencing with classical anatomical atlases, histological preparations, fibre dissection studies, and complementary high-resolution MRI datasets. Representative examples of this process are shown in *Supplementary Figures 1-11* and in the *Segmentation Protocol* (see *Data Repository*).

After transferring all segmentations into MNI space (see *Methods* and *Data Repository* for the normalization protocol), we manually reconstructed unilateral fibre tracts in a collaborative effort, guided primarily by the delineated atlas structures and further supported by established anatomical resources. The reconstructed tracts were subsequently mirrored to the contralateral hemisphere to generate bilateral representations. *Supplementary Table 1* provides detailed information on the individual segmentations and fibre tracts, along with key references used for each structure.

Although the 100 $\mu$m FLASH volume served as the primary geometric reference for segmentation, not all structures were delineated from FLASH contrast alone. In a number of regions, especially where boundaries were subtle or only partially visible on any single slice, delineation relied on the combined use of additional resources, including alternative flip-angle images, histological material from the same specimen, external histological and dissection references, and the topographic relationships to neighbouring structures. Importantly, many boundaries became more interpretable when inspected interactively across axial, coronal, sagittal, and oblique planes and under adjusted contrast settings, rather than in a single static view. Thus, while contrast was not equally strong for all structures on the underlying FLASH image alone, anatomically reasonable delineation was achieved through multimodal integration rather than single-image contrast in isolation.

In total, 48 histological, anatomical, and imaging resources were particularly influential in guiding the construction of the atlas, supporting both manual segmentations and tract reconstructions (also see *Methods*). The principal reference categories include: *Atlases*: Allen BrainSpan Whole Brain Reference Atlas ([9]); Anatomie des centres nerveux, v1([15]); Anatomie des centres nerveux, v2 ([31]); Atlas for Stereotaxy of the Human Brain ([4]); An atlas of the basal ganglia, brain stem and spinal cord, based on myelin-stained material ([32]); Atlas of the central nervous system in man ([5]); Atlas of the Human Brain ([33]); Duvernoy's Atlas of the Brain Stem and Cerebellum ([34]); Fiber Pathways of the Brain ([35]); Human brain in standard MNI space ([36]); Olszewski's and Baxter's Cytoarchitecture of the Human Brainstem ([17]); Organization of brainstem nuclei ([37]); Stereotactic Atlas of the Human Thalamus and Basal Ganglia ([1]); Structure of the human brain: a photographic atlas ([38]); The Human Nervous System ([39]); The Human Central Nervous System ([26]). *Digital Atlases:* AHEAD Atlas ([40]); Allen Human Reference Atlas ([41]); Atlas of the Human Hypothalamus ([42]); Brainstem Navigator ([43]); CIT168 Reinforcement Learning Atlas ([44]); CoBrALAB Atlas ([45]); DISTAL Atlas ([46]); MNI PD25 subcortical ([47]); THOMAS Atlas ([48]). *Seminal neuroanatomical studies*: Beitrag zur Kenntniss des Corpus mammillare und der sogenannten Schenkel des Fornix ([12]); Beitrag zur vergleichenden Anatomie der Substantia nigra, des Corpus Luysii und der Zona incerta ([49]); Connections of the red nucleus ([50]); Efferent pathways of the nucleus of the optic tract in monkey and their role in eye movements ([51]); Experimentelle und pathologisch-anatomische Untersuchungen über die Haubenregion, den Sehhügel und die Regio subthalamica ([52]); Human pallidothalamic and cerebellothalamic tracts: anatomical basis for functional stereotactic neurosurgery ([53]); Neuroanatomical background and functional considerations for stereotactic interventions in the H fields of Forel ([54]); Pallidohypothalamic tract, or X bundle of Meynert, in the rhesus monkey ([11]); Perspective on basal ganglia connections as described by Nauta and Mehler in 1966 ([55]); Projections of the lentiform nucleus in the monkey ([56]); The substantia nigra of the human brain. I. Nigrosomes and the nigral matrix, a compartmental organization based on calbindin D28K immunohistochemistry ([57]); Ueber einen bisher nicht beschriebenen Nervenfasernstrang im Gehirn der Säugetiere und des Menschen ([58]); Untersuchungen über die Haubenregion und ihre oberen Verknüpfungen im Gehirn des Menschen und einiger Säugethiere ([2]); Zur Normalanatomie der Substantia nigra: Versuch einer architektonischen Gliederung ([28]); 9.4 T MR microscopy of the substantia nigra with pathological validation in controls and disease ([59]). *Histology:* Darkfield Microscopy Data ([60], personal archive E.J.L.A.); H.E./TH. Stains ([8], personal archive B.L.E.); Nissl Stains ([9]); Gallyas Silver Stains ([17], personal archive A.K.E.H.). *Fibre dissections*: Klingler Fibre Dissections ([61], personal archive V.M.H.). *High-Resolution MRI Resources*: 100 $\mu$m 7T MRI ([30]); 200 $\mu$m DTI (personal archive B.L.E); BigMac Dataset ([62]). *Supplementary Table 1* further lists segmentation- and fibre tract-specific key sources.

***Supplementary Table 1.* Ontological overview of high-resolution subcortical atlas structures and accompanying manually curated white matter tracts.** In total, n = 180 individual atlas structures were meticulously delineated in native space and transformed into MNI reference space. Additionally, n = 77 anatomical fibre tracts were manually reconstructed unilaterally and subsequently mirrored to the contralateral hemisphere.

| Anatomical name | Abbreviation of atlas definition | (Additional) References |
|---|---|---|
| | Grey matter segmentations | |
| Cerebrum | | |
| Prosencephalon | | |
| Telencephalon | | |
| Cerebral cortex | | |
| Hippocampus | HIP | [62,32,61] |
| Olfactory tubercle | OT | [65,33,5] |
| Cerebral nuclei | | |
| Basal ganglia | | |
| Striatum | | |
| Caudate nucleus (including cell bridges) | ST (Ca) | [39,65], histological preparations (provided by B.L.E., E.J.L.A.) |

| | | |
|---|---|---|
| Putamen (including cell bridges) | ST (Pu) | [39,65], histological preparations (provided by B.L.E., E.J.L.A.) |
| Nucleus accumbens | Nac | [65,32,33,39,64], histological preparations (provided by B.L.E., E.J.L.A.) |
| Globus Pallidus | | |
| External Segment of Globus Pallidus | GPe | [5,8,9,33], histological preparations (provided by B.L.E., E.J.L.A) |
| Internal Segment of Globus Pallidus | GPi | [5,8,33,65], histological preparations (provided by B.L.E., E.J.L.A.) |
| Ventral Pallidum | VP | [65,32,5,33], histological preparations (provided by B.L.E., E.J.L.A.) |
| Basal Forebrain | | |
| Substantia innominata | Si | [65,33,64,23,66,24,25,27], histological preparations (provided by B.L.E., E.J.L.A.) |
| Basal nucleus of Meynert | BM | [65,33,39,27,67–70], histological preparations (provided by B.L.E., E.J.L.A) |
| Diencephalon | | |
| Thalamus | Th | [1] |
| Reticular nucleus | Rt | [4,1,6,7], histological preparations (provided by B.L.E., E.J.L.A.) |
| *Lateral Group* | | |
| *Ventroposterior complex (VPL, VPM, VPI)* | | |
| Ventral posterior medial nucleus (incl. parvocellular division) | VPM | [1] |
| Ventral posterior lateral nucleus (posterior & anterior division) | VPL | [1] |
| Ventral posterior inferior nucleus | VPI | [1] |
| *Ventral lateral posterior & ventral lateral anterior (VLp, VLa)* | | |
| Ventral lateral posterior nucleus (dorsal & ventral division) | VLp | [1] |
| Ventral lateral anterior nucleus | VLa | [1] |
| Ventral anterior nucleus (parvocellular & magnocellular divisions) | VA | [1] |
| Ventral medial nucleus (VM) | VM | [1] |
| *Medial Group* | | |
| Mediodorsal nucleus, magnocellular division | MDmc | [1] |
| Mediodorsal nucleus, parvocellular division | MDpc | [1] |
| Central medial nucleus | CM | [33,1,39], histological preparations (provided by B.L.E., E.J.L.A., A.K.E.H.) |
| Parafascicular nucleus | Pf | [33,1,39],histological preparations (provided by B.L.E., E.J.L.A., A.K.E.H.) |
| Subparafascicular | sPf | [1] |
| Central lateral nucleus | CL | [1] |
| *Midline* | | |
| Central medial nucleus | CeM | [1] |
| *Posterior Group* | | |
| Medial geniculate nucleus | MGN | [1] |

| | | |
|---|---|---|
| Lateral geniculate nucleus | LGN | [1] |
| Posterior nucleus | Po | [1] |
| Limitans nucleus | Li | [1] |
| Lateral posterior | LP | [1] |
| *Pulvinar nuclei (PuM, PuI, PuL, PuA)* | | |
| Medial Pulvinar & Lateral Pulvinar | PuMPuL | [1] |
| Inferior Pulvinar | PuI | [1] |
| Anterior Pulvinar | PuA | [1] |
| *Anterior Group* | | |
| Anteroventral nucleus | AV | [1] |
| Anteromedial nucleus | AM | [1] |
| Anterodorsal nucleus | AD | [1] |
| Lateral dorsal nucleus | LD | [1] |
| Epithalamus | | |
| Pineal body | Pin | [1,64,71,72] |
| Habenula | Hb | [33,1,73,74] |
| Lateral habenular nucleus | Hbl | [33,1,73,74] |
| Medial habenular nucleus | Hbm | [33,1,73,74] |
| Subthalamus | | |
| Subthalamic nucleus | STN | [32,33,1,8] |
| Zona incerta | ZI | [4,5,33,1],histological preparations (provided by B.L.E., E.J.L.A.) |
| Field of Forel H1 | H1 | [15,31,5,33,1,54,2], histological preparations (provided by B.L.E., E.J.L.A |
| Field of Forel H2 | H2 | [15,31,5,33,1,54,2], histological preparations (provided by B.L.E., E.J.L.A.) |
| Field of Forel H | H | [15,31,5,33,1,54,2], histological preparations (provided by B.L.E., E.J.L.A) |
| Hypothalamus | | |
| Mammillary body | MB | [65,33] |
| Lateral tuberal nucleus | NTL | [15,20–22,31] |
| Mesencephalon | | |
| Tegmentum | | |
| Periaqueductal gray substance | PAG | [33,17,39,75], histological preparations (provided by A.K.E.H.) |
| Red nucleus | RN | [33,17,37,39,50], histological preparations (provided by B.L.E., A.K.E.H) |
| Capsule of the Red nucleus | RNc | [33,17,37,39,50], histological preparations (provided by B.L.E., A.K.E.H) |
| Rostral interstitial nucleus of the medial longitudinal fasciculus | Rimlf | [7,16,17,37,76,77], histological preparations (provided by A.K.E.H) |
| Substantia nigra | SN | [65,33,17,78,28,79], histological preparations (provided by B.L.E., A.K.E.H) |
| Nigrosome 1 | N1 | [57,59] |
| Nigrosome 2 | N2 | [57,59] |
| Nigrosome 3 | N3 | [57,59] |
| Nigrosome 4 | N4 | [57,59] |

| | | |
|---|---|---|
| Sano Field A | SanA | [28,78] |
| Ventral Tegmental Area of Tsai | | |
| Parabrachial pigmented nucleus | Pbn | [17,65], histological preparations (provided by A.K.E.H) |
| Paranigral nucleus | PNn | [17,37,65], histological preparations (provided by B.L.E., A.K.E.H) |
| Caudal linear nucleus | CLin | [17,37,65], histological preparations (provided by B.L.E., A.K.E.H) |
| Interpeduncular nucleus | IPn | [17,37,65], histological preparations (provided by B.L.E.) |
| Oculomotor nucleus | IIIn | [17,37], histological preparations (provided by B.L.E., A.K.E.H) |
| Edinger-Westphal nucleus | EW | [17,37], histological preparations (provided by B.L.E., A.K.E.H) |
| Trochlear nucleus | Ivn | [17,37], histological preparations (provided by B.L.E.) |
| Tectum | | |
| Superior colliculus | SC | [29,37] |
| Inferior colliculus | IC | [17,37] |
| Rhombencephalon | | |
| Cerebellum | | |
| Dentate nucleus | DN | [65,64], histological preparations (provided by E.J.L.A.) |
| Fastigial nucleus | FN | [65,64], histological preparations (provided by E.J.L.A) |
| Interposed nucleus | InPn | [65,64], histological preparations (provided by E.J.L.A.) |

White matter segmentations

| | | |
|---|---|---|
| Cerebrum | | |
| Prosencephalon | | |
| Lateral medullary lamina | lml | [65,15,31,33], histological preparations (provided by B.L.E.) |
| Medial medullary lamina | mml | [65,15,31,33], histological preparations (provided by B.L.E.) |
| Incomplete medullary lamina | icml | [65,15,31,33], histological preparations (provided by B.L.E.) |
| Supplementary medullary lamina | supml | [15,31,65,33], histological preparations (provided by B.L.E.) |
| Anterior commissure | ac | [65,33,35,80,81], histological preparations (provided by B.L.E.) |
| Optic nerve, chiasm, tract | onxt | [65,33,64] |
| Supraoptic pathway commissure | sox | [65,33,18,19], histological preparations (provided by B.L.E.) |
| Habenular commissure | Hbc | [33,29,64],histological preparations (provided by A.K.E.H) |
| Ansa lenticularis | al | [65,15,31,5,33,34,1,39,56,55,64], histological preparations (provided by B.L.E., E.J.L.A) |
| Fasciculus thalamicus | fct | [15,31,5,33,34,1,39,64], histological preparations (provided by B.L.E., E.J.L.A.) |
| Pallidohypothalamic tract | palhy | [33,11,13], histological preparations (provided by B.L.E., E.J.L.A) |
| Fornix | fx | [65,33] |
| Mammillothalamic tract | mtt | [33,39] |
| Principal fasciculus | pf | [33,12,64], histological preparations (provided by B.L.E.) |
| Stria medullaris | sm | [5,33,34,1], histological preparations (provided by B.L.E.) |
| Stria terminalis | st | [5,33,34,1], histological preparations (provided by B.L.E.) |

| | | |
|---|---|---|
| Mesencephalon | | |
| Posterior commissure | pc | [17,35,64],histological preparations (provided by A.K.E.H) |
| Commissure of the Superior colliculus | SCc | [17,64],histological preparations (provided by A.K.E.H) |
| Oculomotor nerve | CNIII | [5,17,34], histological preparations (provided by B.L.E., A.K.E.H) |
| Trochlear nerve | CNIV | [5,17,34], histological preparations (provided by B.L.E., A.K.E.H) |
| Fasciculus retroflexus | fr | [5,33,34,17,39,82], histological preparations (provided by B.L.E., A.K.E.H) |
| Rhombencephalon | | |
| Medial longitudinal fasciculus | mlf | [17,32,65], histological preparations (provided by B.L.E., A.K.E.H) |
| Medial lemniscus | ml | [32,5,33,34,64], histological preparations (provided by B.L.E., A.K.E.H) |
| Lateral lemniscus | ll | [32,5,33,34,64], histological preparations (provided by B.L.E., A.K.E.H) |

Manual curated white matter tracts

| Anatomical description | Abbreviation | References | Atlases |
|---|---|---|---|
| Basal Ganglia Pathways | | | |
| Pallidothalamic Pathways | | | |
| Ansa lenticularis | | Klingler fibre dissections (provided by V.M.H.) | FOCUS atlas |
| Branch to Centromedian-parafascicular complex of thalamus | al_to_cmpf | [56,64,83–90] | |
| Branch to ventral anterior thalamic nucleus; corresponding to thalamic fasciculus | al_to_va | [33,54,56,64,83–91] | |
| Fasciculus lenticularis | | Klingler fibre dissections (provided by V.M.H.) | FOCUS atlas |
| Branch to Centromedian-Para fascicular complex of thalamus | fl_to_cmpf | [56,83–85,88,90] | |
| Branch to ventral anterior thalamic nucleus; corresponding to thalamic fasciculus | fl_to_va | [33,56,83–85,88,90] | |
| Pallidum to MD connection | gpi_md | [92,93] | FOCUS atlas, [1] |
| Pallidohypothalamic Pathway | | | |
| Pallidohypothalamic tract | palhy | [33,11,13] | FOCUS atlas, [42] |
| Pallidosubthalamic Pathways | | | |
| Ansa subthalamica | as | [56,94,95], histological preparations (provided by E.J.L.A.) | FOCUS atlas |
| GPe – STN connection | | [83,96,97,64] | FOCUS atlas |
| GPe – STN connection, caudal portion | gpe_stn_cau | | |
| GPe – STN connection, rostral portion | gpe_stn_ros | | |
| GPi – PPN connection | gpi_to_ppn | [98–100] | FOCUS atlas, [101] |
| GPi – STN connection | | [64,83,97] | FOCUS atlas |
| GPi – STN caudal part | gpi_stn_cau | | |
| GPi – STN rostral part | gpi_stn_ros | | |

| | | | |
|---|---|---|---|
| Striatopallidofugal Pathways | | | |
| Caudate – GPi connection | | [64,102–105] | FOCUS atlas, [44,46] |
| Caudate – GPi caudal portion | cd_to_gpi_cau | | |
| Caudate – GPi rostral portion | cd_to_gpi_ros | | |
| Caudate-SN connection | | [64,102–107] | FOCUS atlas, [42,44,46] |
| Caudate – SN caudal portion | cd_to_sn_cau | | |
| Caudate – SN rostral portion | cd_to_sn_ros | | |
| Putamen – GPi | | [64,102–105,56] | FOCUS atlas, [44,46] |
| Putamen – GPi caudal part | put_to_gpi_cau | | |
| Putamen – GPi rostral part | put_to_gpi_ros | | |
| Putamen – SN connection | | [64,102–104,106,107,105] | FOCUS atlas, [42,44,46] |
| Putamen – SN caudal part | put_to_sn_cau | | |
| Putamen – SN rostral part | put_to_sn_ros | | |
| Ventral Striatum - SN/VTA | | [64,102,106–110] | FOCUS atlas, [42,44,46] |
| Supracommisural protion | venStr_to_snvta_overAC | | |
| Infracommisural portion | venStr_to_snvta_underAC | | |
| Nigrothalamic Pathways | | | |
| Substantia Nigra to Mediodorsal thalamic nucleus | | | |
| *Parvocellular part* | snr_to_mdpc | [111–115] | FOCUS atlas, [1,44] |
| *Magnocellular part* | snr_to_mdmc | [111–115] | FOCUS atlas, [1,44] |
| Substantia Nigra to Ventral Anterior thalamic nucleus | | | |
| *Magnocellular part* | snr_to_VAmc | [111–115] | FOCUS atlas, [1,44] |
| Subthalamic Efferent Pathways | | | |
| STN-Pedunculopontine nucleus connection | stn_to_ppn | [116–118] | FOCUS atlas |
| STN-Substantia nigra connection | stn_to_sn | [117–119] | FOCUS atlas |
| Hyperdirect Pathways | | | |
| Brodmann area 4 – STN | BA04_stn | [97,120–122] | FOCUS atlas, [123] |
| Brodmann area 6 – STN | BA06_stn | [97,120–122] | FOCUS atlas, [123] |
| Brodmann area 8 – STN | BA08_stn | [64,97,120,124–126] | FOCUS atlas, [123] |
| Brodmann area 9/10 – STN | BA0910_stn | [97,120] | FOCUS atlas, [123] |
| Brodmann area 11 – STN | BA11_stn | [120,127,128] | FOCUS atlas, [123] |
| Brodmann area 24 & 32 – STN | BA2432_stn | [97,120,128] | FOCUS atlas, [123] |
| Brodmann area 46 – STN | BA46_stn | [97,120,128] | FOCUS atlas, [123] |
| Descending Pathways | | | |
| Corticofugal Pathways | | | |
| Brodmann area 1,2, and 3 to Internal Capsule | BA123_ic | [64,121,129–131] | FOCUS atlas, [123] |
| Brodmann area 1,2, and 3 (face area) – thalamus (VPM) | BA123_vpm | [64,132] | FOCUS atlas, [123] |
| Brodmann area 1,2, and 3 (upper and lower extremity areas) – thalamus (VPL) | BA123_vpl | [64,132] | FOCUS atlas, [123] |
| Brodmann area 4 to Internal Capsule | BA04_ic | [97,120] | FOCUS atlas, [123] |

| | | | |
|---|---|---|---|
| Brodmann area 6 to Internal Capsule | BA06_ic | [97,120,133] | FOCUS atlas, [123] |
| Brodmann area 8 to Internal Capsule | BA08_ic | [97,120,124,125,133] | FOCUS atlas, [123] |
| Brodmann area 9/10 to Internal Capsule | BA0910_ic | [97,120,127,133,134] | FOCUS atlas, [123] |
| Brodmann area 11 (medial) to Internal Capsule | BA11m_ic | [97,120,127,128,134,135] | FOCUS atlas, [123] |
| Brodmann area 11 (lateral) to Internal Capsule | BA11L_ic | [97,120,127,128,135] | FOCUS atlas, [123] |
| Brodmann area 24 & 32 to Internal Capsule | BA2432_ic | [97,120,127,128,134] | FOCUS atlas, [123] |
| Brodmann area 45 to Internal Capsule | BA45_ic | [128,134] | FOCUS atlas, [123] |
| Brodmann area 46 to Internal Capsule | BA46_ic | [128,133,134] | FOCUS atlas, [123] |
| Brodmann area 47 to Internal Capsule | BA47_ic | [127,128,133–135] | FOCUS atlas, [123] |
| Rubrospinal Pathway | | | |
| Rubrospinal tract | rst | [32,34,37,64] | |
| Thalamic Affarent Pathways | | | |
| Cerebellothalamic Pathways | | | |
| Dentatorubrothalamic Tract | | Klingler fibre dissections (provided by V.M.H.), | FOCUS atlas |
| Decussating Dentorubrothalamic Tract (dorsal dentate to SMA projecting zones of VL) | dDRTT_VL_cau | [33,64,54,89,91,90,136–139] | |
| Decussating Dentorubrothalamic Tract (dorsal dentate to M1 projecting zones of VL) | dDRTT_VL_ros | [33,64,54,89,91,90,136–138] | |
| Decussating Dentorubrothalamic Tract (ventral dentate to VL-MD) | dDRTT_ven | [33,64,54,89,136–138,140–142] | |
| Somatosensory Thalamic Afferents | | | |
| Medial lemniscus | medl | [64,54,143–145] | FOCUS atlas |
| Trigeminal lemniscus | tril | [64,54,143–145] | FOCUS atlas |
| Thalamic Efferent Pathways | | | |
| Centromedian to dorsolateral postcomissural Putamen | cm_to_put | [146–151] | FOCUS atlas |
| Centromedian to STN | cm_to_stn | [147,148,150] | FOCUS atlas, [46] |
| Parafascicular nucleus – Striatum | | [148–151] | FOCUS atlas, [42] |
| Pf to Caudate Nucleus (body and tail) | pf_to_cau_bodytail | | |
| Pf to Caudate Nucleus (head) | pf_to_cau_ic | | |
| Pf to Putamen (head) | pf_to_put_ic | | |
| Pf to Pallidum (collateral) | pf_to_pall | | |
| Pf to Ventral Striatum | pf_to_put_venStr | | |
| Parafasciculus nucleus to STN | pf_to_stn | [148,150] | FOCUS atlas |
| Parafasciculus nucleus to Ventral Tegmental Area | pf_to_vta | [148,150] | FOCUS atlas |

| | | | |
|---|---|---|---|
| Limbic Pathways | | | |
| Amygdalofugal Pathways | | | |
| Stria terminalis | | [33,152–154] | FOCUS atlas, [42,44,155,156] |
| | st_cmAmyg_to_bnst | | |
| | st_cmAmyg_to_hypothal | | |
| | st_cmAmyg_to_NAc | | |
| Ventral amygdalofugal pathway | | [33,153,157–160] | FOCUS atlas, [1,155,156] |
| Basolateral Amygdala to Hypothalamus | vafp_blAmyg_to_hypothal | | |
| Basolateral Amygdala to Mediodorsal Thalamic Nucleus, magnocellular part | vafp_blAmyg_to_mdmc | | |
| Basolateral Amygdala to Septum | vafp_blAmyg_to_septal | | |
| Habenular Pathways | | | |
| Fasciculus retroflexus | | [5,33,34,17,39,82], Klingler fibre dissections (provided by V.M.H.) | FOCUS atlas |
| Branch to centromedian thalamic nucleus | frf_to_cm | | |
| Branch to Habenula | frf_to_Hb | | |
| Stria medullaris | sm | [5,33,34,1] | FOCUS atlas |
| Hippocampal System Pathways | | | |
| Fornix | | [65,33,161], Klingler fibre dissections (provided by V.M.H.) | FOCUS atlas |
| Postcommisural Part | fx_postcomm | | |
| Precommisural Part | fx_precomm | | |
| Mammillothalamic tract including the parafascicular fasciculus | mtt | [33,39,162], Klingler fibre dissections (provided by V.M.H.) | FOCUS atlas |
| Mesolimbic Pathways | | | |
| Medial Forebrain Bundle, Ventral Tegmental Area projection pathway to Nucleus Accumbens | mfb_vta_to_nac | [33,64,111,163–167] | FOCUS atlas, [44] |
| Pathways related to Visual System | | | |
| Commissure of the superior colliculus | scc | [29,64] | FOCUS atlas |
| Oculomotor nerve | CNIII | [5,29,34] | FOCUS Atlas |

***Supplementary Table 2.*** **Demographic and clinical characteristics of key anatomical reference resources used in this study.**

| Resource | | n | Sex | Age | Clinical History |
|---|---|---|---|---|---|
| Postmortem 100 $\mu$m 7T MRI | [30] | 1 | f | 58 years | History of lymphoma and stem cell transplantation; no neurological or psychiatric disease; died of hypoxic respiratory failure due to viral pneumonia |

| | | | | | |
|---|---|---|---|---|---|
| MNI152NLin2009bAsym template | [168,169] | 152 | f (n=66)<br>m (n=86) | 25.0 ± 4.9 years | Normative young adults; individual clinical histories not reported |
| Klingler Fibre Dissections | [61], personal archive V.M.H. | 20 | Not available | Not available | No history of neurological, neuropsychiatric, or infectious diseases |
| H.E./TH. Stains | [8], personal archive B.L.E. | 2 | f (n=1)<br>m (n=1) | 53 years<br>49 years | No neurological disease; deaths related to systemic malignancy/sepsis in two cases; only mild incidental neuropathological changes and no tractography-relevant lesions |
| Gallyas Silver Stains | [29], personal archive A.K.E.H. | 3 | f (n=1)<br>m (n=2) | 53.7 ± 30.3 years | No known neurological disease or neuropathological abnormalities; causes of death were myocardial infarction, cardiogenic shock, and drug overdose |
| Dark-field Microscopy Data | [60] , personal archive E.J.L.A. | 3 | f (n=1)<br>m (n=2) | 59.3 ± 5.8 years | No previous neurological disease; causes of death were myocardial infarction (n=2) and blunt trauma (n=1) |
| Allen Brain-Span Whole Brain Reference Atlas | [9] | 1 | f | 34 years | No history of neurological disease or brain abnormality, cause of death not reported |
| Atlas for Stereotaxy of the Human Brain | [1] | 7 | f (n=4)<br>m (n=3) | 61.0 ± 10.6 years | Non-neurological causes of death: carcinoma (n=4; lung n=3, anal n=1) and cardiovascular disease (n=3) |
| Olszewski's and Baxter's Cytoarchitecture of the Human Brainstem | [29] | 16 | f (n=6)<br>m (n=9)<br>not reported (n=1) | 7th gestational month–77 years; 1 not reported | Heterogeneous autopsy series, predominantly systemic, cardiopulmonary, malignant, and vascular pathology; one subarachnoid hemorrhage, n =1 not reported |
| Stereotactic Atlas of the Human Thalamus and Basal Ganglia | [4] | 2 | m | 40 years<br>51 years | Clinical history and cause of death not reported |
| The Human Nervous System | [1] | 1 | m | 24 years | No history of neurological disease, cause of death: hypovolemic shock |

| | | | | | | |
|---|---|---|---|---|---|---|
| Clinical Dystonia cases | Previously unpublished cases; institutional medical records and imaging archive, Tokyo Women's Medical University | | | | | |
| | | Case 1 | 1 | f | 34 years | Segmental dystonia |
| | | Case 2 | 1 | f | 48 years | Segmental dystonia |
| | | Case 3 | 1 | m | 66 years | Focal cervical dystonia |
| Parkinson's Disease Cohort (discovery) | [170,171] | | 129 | f (n=36)<br>m (n=93) | 57.4 ± 9.6 years | Parkinson's Disease |
| Parkinson's Disease Cohort (test) | [170,171] | | 89 | f (n=39)<br>m (n=50) | 60.5 ± 8.9 years | Parkinson's Disease |

*Interrater Agreement Analysis*

HF led the generation of the 180 native-space high-resolution segmentations and performed the comparative neuroanatomical synthesis across the complete set of structures using the anatomical resources described above. For selected structures with anatomically ambiguous or poorly resolved boundaries, delineations were reviewed with co-authors contributing relevant structure-specific expertise and refined where appropriate. To complement this expert-informed development process, we performed a quantitative interrater agreement analysis in four exemplar structures.

Independent re-segmentation of the complete set of 180 structures would have required extensive manual delineation and comparative anatomical synthesis. We therefore selected four exemplar structures that differed in size, geometry, tissue contrast, and boundary definition: the dentate nucleus (DN), globus pallidus internus (GPi), red nucleus (RN), and subthalamic nucleus (STN). Three additional raters (R.D., L.K., E.J.L.A.) independently delineated each structure using the same anatomical definitions, reference materials, and detailed *Segmentation Protocol* provided in the *Data Repository*. Together with the original segmentation, this yielded four masks per structure and six unique pairwise comparisons. All corresponding binary masks were defined on the same native voxel grid. Interrater agreement was quantified using the Dice similarity coefficient, mean symmetric surface distance (MSD), and 95th-percentile Hausdorff distance (HD95).

Surface distances are reported in millimetres. Because each segmentation contributed to multiple pairwise comparisons, the mean and standard deviation across the six rater pairs is reported as descriptive summary statistics.

*Interrater agreement across exemplar structures*

Across the six unique pairwise comparisons, Dice coefficients ranged from 0.902 to 0.965 for the STN, from 0.904 to 0.979 for the RN, from 0.884 to 0.966 for the GPi, and from 0.766 to 0.891 for the DN.

Mean Dice coefficients were 0.921 ± 0.023 for the STN, 0.937 ± 0.028 for the RN, 0.929 ± 0.028 for the GPi, and 0.823 ± 0.052 for the DN. Mean MSD values were 0.118 ± 0.033 mm, 0.141 ± 0.065 mm, 0.156 ± 0.066 mm, and 0.075 ± 0.020 mm, respectively; the corresponding mean HD95 values were 0.363 ± 0.044 mm, 0.459 ± 0.129 mm, 0.519 ± 0.147 mm, and 0.246 ± 0.071 mm. Individual pairwise values are reported in *Supplementary Table 3* and visualized in *Supplementary Figure 13.*

***Supplementary Table 3.*** **Pairwise interrater agreement across exemplar structures.** *Note*: Rater 1 denotes the original segmentation; Raters 2–4 denote the three additional independent segmentations. Summary statistics are descriptive and were calculated across the six unique pairwise comparisons.

| Structure | Rater pair | Dice coefficient | MSD (mm) | HD95 (mm) |
|---|---|---|---|---|
| STN | Rater 1 vs. Rater 2 | 0.925 | 0.106 | 0.343 |
| | Rater 1 vs. Rater 3 | 0.907 | 0.132 | 0.343 |
| | Rater 1 vs. Rater 4 | 0.965 | 0.059 | 0.347 |

| | | | | |
|---|---|---|---|---|
| | Rater 2 vs. Rater 3 | 0.916 | 0.118 | 0.324 |
| | Rater 2 vs. Rater 4 | 0.913 | 0.141 | 0.447 |
| | Rater 3 vs. Rater 4 | 0.902 | 0.154 | 0.371 |
| | **Mean ± SD** | **0.921 ± 0.023** | **0.118 ± 0.033** | **0.363 ± 0.044** |
| RN | Rater 1 vs. Rater 2 | 0.917 | 0.189 | 0.526 |
| | Rater 1 vs. Rater 3 | 0.958 | 0.086 | 0.250 |
| | Rater 1 vs. Rater 4 | 0.979 | 0.048 | 0.371 |
| | Rater 2 vs. Rater 3 | 0.904 | 0.219 | 0.618 |
| | Rater 2 vs. Rater 4 | 0.923 | 0.171 | 0.493 |
| | Rater 3 vs. Rater 4 | 0.939 | 0.132 | 0.495 |
| | **Mean ± SD** | **0.937 ± 0.028** | **0.141 ± 0.065** | **0.459 ± 0.129** |
| GPi | Rater 1 vs. Rater 2 | 0.949 | 0.111 | 0.393 |
| | Rater 1 vs. Rater 3 | 0.913 | 0.198 | 0.634 |
| | Rater 1 vs. Rater 4 | 0.966 | 0.073 | 0.393 |
| | Rater 2 vs. Rater 3 | 0.933 | 0.141 | 0.425 |
| | Rater 2 vs. Rater 4 | 0.927 | 0.155 | 0.516 |
| | Rater 3 vs. Rater 4 | 0.884 | 0.259 | 0.754 |
| | **Mean ± SD** | **0.929 ± 0.028** | **0.156 ± 0.066** | **0.519 ± 0.147** |
| DN | Rater 1 vs. Rater 2 | 0.821 | 0.078 | 0.238 |
| | Rater 1 vs. Rater 3 | 0.770 | 0.085 | 0.250 |
| | Rater 1 vs. Rater 4 | 0.877 | 0.063 | 0.343 |
| | Rater 2 vs. Rater 3 | 0.891 | 0.041 | 0.124 |
| | Rater 2 vs. Rater 4 | 0.814 | 0.085 | 0.252 |
| | Rater 3 vs. Rater 4 | 0.766 | 0.096 | 0.268 |
| | **Mean±SD** | **0.823 ± 0.052** | **0.075 ± 0.020** | **0.246 ± 0.071** |

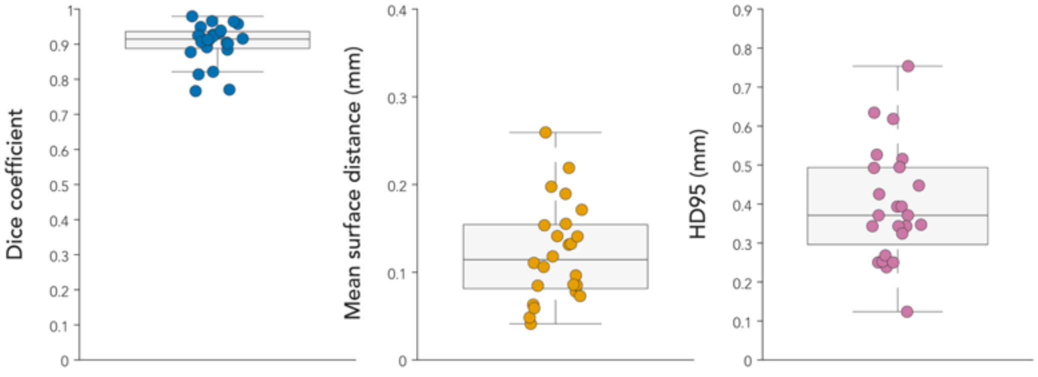

***Supplementary Figure 13*. Pairwise interrater agreement across exemplar structures.** Boxplots summarize Dice similarity coefficients, mean symmetric surface distances (MSD), and 95th-percentile Hausdorff distances (HD95) across the six unique pairwise comparisons among four raters for the subthalamic nucleus, red nucleus, globus pallidus internus and dentate nucleus. Summary statistics are reported in *Supplementary Table 3*.

The generally high agreement observed across four anatomically distinct exemplar structures indicates that the corresponding anatomical definitions and segmentation protocol can be applied consistently by different observers. Because this targeted analysis was restricted to four structures, it does not establish interrater agreement for each of the 180 segmentations individually; rather, it provides quantitative support for the transferability of the delineation framework beyond the primary atlas developer.

2. Refined coregistration workflow for precise subcortical alignment

To be able to make use of these segmentations in clinical (DBS) neuroimaging studies, they had to be transferred to a standard space. Lead-DBS employs the MNI152 2009b Nonlinear Asymmetrical space ([168]) as its reference. While [30] presented a native-to-MNI transformation primarily focusing on overall alignment and certain DBS targets, additional improvements were needed for the registration of the thalamus, subthalamus, and mesencephalon in this study. The warp of the basal ganglia, specific telencephalic regions, and the basal forebrain was also refined to fully utilize the increased detail offered by high-resolution structures in the standard space.

The segmentation-supported ANTs coregistration protocol is described in detail in the *Methods* section. A step-by-step guideline has been deposited in the *Data Repository* to facilitate reproducible use of the workflow. A comprehensive listing of the supporting anchor structures and the full sequence of iterative steps is provided below.

In brief, the native-to-MNI space transformation, adapted from[30], was conducted iteratively through meticulous creation and registration of congruent segmentations in both native and MNI152NLinAsym space. This process comprised more than 37 sequential coregistrations, each building upon the previous one. Anchor structures were bilaterally segmented and refined across iterations, with local manual adjustments applied in selected regions using WarpDrive ([172]). For computational efficiency, the individual anchor structures were grouped into three composite masks (for details see *Supplementary Table 4*). Qualitative assessment of the resulting registrations suggested that this grouping also had a beneficial effect on the overall topographic alignment of the warp.

The resulting transformation demonstrated markedly improved alignment within the thalamus, the mesencephalo-diencephalic junction, and the mesencephalon, as well as in targeted regions of the telencephalon and basal forebrain (see *Supplementary Figure 12*).

To illustrate the improvements in spatial normalization achieved by our updated pipeline, we selected the mammillothalamic tract (mtt) as an anatomically well-defined test case (see *Supplementary Figure 12)*. The mtt originates from the mammillary bodies, courses subthalamically, and ascends through the thalamus, making it particularly suitable to evaluate alignment accuracy in the subthalamic, mesodiencephalic transition, and thalamic zones. Importantly, the tract is clearly identifiable in the T1-weighted MNI template, allowing reliable manual segmentation.

We applied both the transformation developed in the current study and the method by [30] to the same high-resolution 7T segmentation of the mtt in native space. Both resulting segmentations were then compared to a manually defined mtt in standard space ("native MNI"), which served as ground truth.

Spatial correspondence was quantified using binarized nifti masks in 0.5 mm MNI152 space. We computed the Dice coefficient, Jaccard index, precision, and recall for each transformed segmentation. All metrics were computed voxelwise. The Dice coefficient was defined as $2 \times |A \cap B| / (|A| + |B|)$, the Jaccard index as $|A \cap B| / |A \cup B|$, precision as $|A \cap B| / |A|$, and recall as $|A \cap B| / |B|$, where A and B represent the binary masks of the transformed and reference segmentations, respectively, $|\cdot|$ denotes the number of nonzero voxels (i.e., the voxel count of a set), and $\cap$ and $\cup$ denote voxelwise intersection and union.

The transformation of the current study yielded a Dice coefficient of 0.7778, a Jaccard index of 0.6364, a precision of 0.8446, and a recall of 0.7208, with a resulting volume of 175.38 $mm^3$. In contrast, the transformation by [30] reached a Dice coefficient of only 0.2497, a Jaccard index of 0.1427, a precision of 0.3209, and a recall of 0.2044, with a corresponding volume of 130.88 $mm^3$. The volume of the native MNI reference segmentation was 205.50 $mm^3$.

These results demonstrate a more than threefold increase in anatomical overlap using the same input segmentation. It is important to note that ventral portions of the tract-namely the principal fasciculus (prf), from which the mtt emerges-are not captured in the T1-based ground truth segmentation due to limited visibility. This likely leads to a systematic underestimation of both Dice and precision scores. Nevertheless, the consistent and substantial gain across all metrics clearly reflects the improved anatomical accuracy of the updated normalization pipeline.

The mammillothalamic tract served as a straightforward yet illustrative example of how spatial misalignments in standard space can have substantial anatomical consequences. While the mtt is clearly identifiable in T1-weighted MNI space and thus allows direct comparison, many adjacent structures, such as thalamic and subthalamic nuclei, are less well delineated and cannot be directly assessed visually. The improved alignment of the mtt suggests that the updated pipeline yields more accurate spatial registration more broadly. As one of over 31 subcortical anchor structures used during normalization, the mtt exemplifies the increased topographic precision that the refined transformation achieves across subcortical and mesodiencephalic regions.

***Supplementary Table 4.*** **Overview of Iteratively Generated Anchor Structures.** The listed segmentations represent anatomical regions selected to constrain and refine the native-to-MNI transformation in areas of particular relevance to the atlas. During the iterative registration procedure, individual anchor segmentations were introduced, visually evaluated, and refined as needed. Throughout this process, anatomically related anchor segmentations were progressively combined and refined within three composite normalization masks. These masks integrated the accumulating anatomical constraints into spatially coherent and computationally efficient targets for deformation-field optimization. Additional single anatomical anchors provided targeted constraints at selected stages. The MNI reference image used to delineate each segmentation is also listed. All anchor segmentations are available in the *Data Repository*.

| Normalization mask | Anchor segmentation ID | (Aggregated) anatomical structures | MNI reference |
|---|---|---|---|
| Mask 1 | 1 | Pu, Ca, CaPu, Nac, Si, OT | t1 |
| | 2 | GPe | t1 |
| | 3 | GPi | t1 |
| | 4 | VP | t1 |
| | 5 | STN | t1/ t2 |
| | 6 | AQM | t1 |
| | 7 | AQ | t1 |
| | 8 | LQ | t1 |
| Mask 2 | 9 | SN, Pbn | t1 |
| | 10 | RN, RNc | t1/t2 |
| | 11 | PAG | t1 |
| | 12 | Pc, sCc | t1 |
| | 13 | Fx, mtt, MB, prf | t1 |
| | 14 | Rt | t1 |
| | 15 | ac | t1 |
| | 16 | Opn, ox, opt | t1 |
| | 17 | Parts of MFC, A25 | t1 |
| Mask 3 | 18 | AV, AD | t1 |
| | 19 | CM | t1 |
| | 20 | Hb (Hbm, Hbl) | t1 |
| | 21 | LGN | t1 |
| | 22 | MGN | t1 |
| | 23 | MD_Th | t1 |
| | 24 | Rt (ref.) | t1 |
| Single anatomical anchors | | | |
| | 25 | DN | t2 |
| | 26 | SCP | t1 |
| | 27 | V | t1 |
| | 28 | HIP | t1/t2 |
| | 29 | CRT | t1/t2 |
| | 30 | Th-outline (coarse thalamic alignment) | t1/t2 |
| | 31 | Pin (coarse global alignment) | t1 |

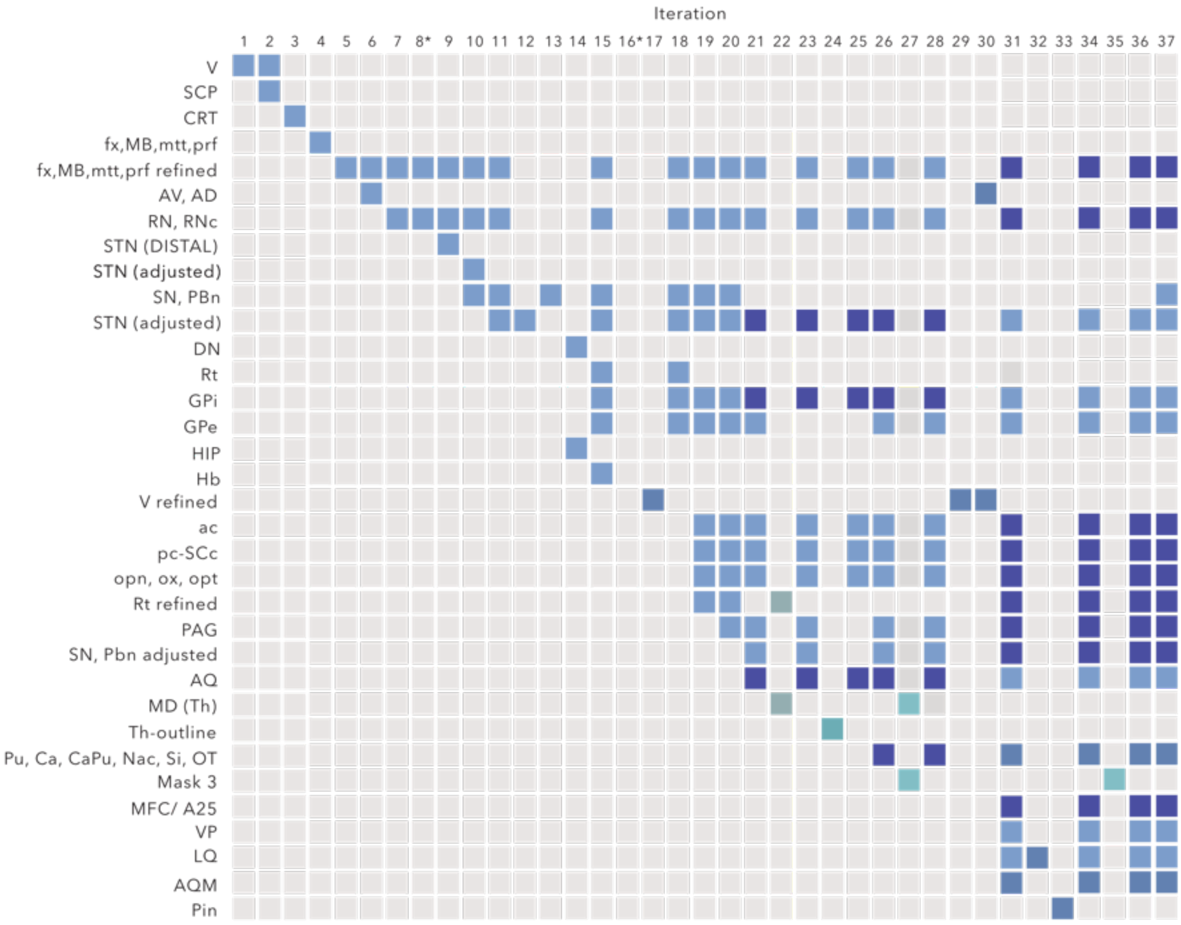

***Supplementary Figure 14.*** **Segmentation-Supported ANTs Coregistration Protocol.** The native-to-MNI space transformation, following the approach of Edlow et al. (2019), was conducted iteratively through meticulous creation and registration of congruent segmentations in both native and MNI152NLinAsym space. This process included over 37 sequential coregistrations, each building on the previous. Anchor structures were bilaterally segmented manually in both spaces and refined iteratively. Manual local adjustments were applied in selected areas using Warp-Drive[172]. The resulting transformation demonstrates exceptional refinement, particularly in the thalamic region, mesencephalo-diencephalic junction, mesencephalon, as well as in targeted regions of the telencephalon and basal forebrain. *Please note*: For computational efficiency, various ROIs were combined into masks, with their associations visually represented by corresponding colours in the figure. *: Adjusted ANTs-settings/ additional overall ANTs coregistration performed.

*Quantitative assessment of anchor alignment*

The normalization strategy was implemented as an iterative process of anatomically guided deformation-field refinement (*Supplementary Figure 14*). Structures and regions that are of particular relevance to the atlas and visible on native and MNI templates were delineated in both spaces and introduced progressively as spatial constraints. After each registration step, correspondence of the relevant anatomical boundaries was inspected in detail, and anchor segmentations were iteratively refined. This cycle was repeated until stable anatomical correspondence had been achieved across the principal atlas regions.

Throughout this process, anatomically related anchor segmentations were progressively integrated into three composite normalization masks (Masks 1-3; *Supplementary Table 4).* These masks represented the cumulative anatomical constraints established over the preceding iterations and provided spatially coherent targets for final deformation-field optimization. Registration quality was assessed by meticulous visual inspection after each iteration. To provide quantitative validation, we additionally calculated overlap metrics for the three composite masks as a summary measure of final registration accuracy, complemented by structure-level metrics for the individual segmentations.

To quantify the spatial accuracy of the final transformation, corresponding segmentation masks were brought into the same reference space. The MNI masks were already defined in MNI space, whereas the corresponding native-space masks were transformed into MNI space using the deformation field generated by the registration procedure developed in this study. Each analysis therefore compared an MNI-space segmentation with its native-space counterpart after application of the study-specific transformation.

For each mask pair, both images were resliced to the same fixed T1-weighted reference image using nearest-neighbour interpolation to preserve their binary definitions. The resliced images were subsequently binarized by thresholding all non-zero voxels. Voxel-wise overlap was quantified using the Dice similarity coefficient, defined as $2|A \cap B|/(|A| + |B|)$, where A and B denote the two binary masks. Surface agreement was quantified using the mean surface distance (MSD). Surface voxels were extracted using a 26-connected neighbourhood definition. For each surface voxel in one mask, the Euclidean distance to the nearest surface voxel in the corresponding mask was calculated in physical millimetre space using the voxel dimensions of the fixed reference image. Distances were calculated in both directions, concatenated, and summarized by their mean to obtain the MSD. The median and standard deviation of the concatenated bidirectional distances were additionally recorded.

The three composite normalization masks yielded Dice coefficients of 0.914, 0.815, and 0.786 for Masks 1-3 (mean ± SD, 0.838 ± 0.067). These values quantify the final correspondence of the aggregated anatomical constraints used to optimize the deformation field. Across the individual segmentation masks, the mean Dice coefficient was 0.775 (median, 0.799; range, 0.386-0.926), and the mean MSD was 0.388 mm (median, 0.283 mm; range, 0.118-1.712 mm). Dice coefficients for the individual anatomical anchors and masks are shown in *Supplementary Figure 15.* Distributions of dice coefficients and mean surface distances are additionally summarized in *Supplementary Figure 16.*

The cerebellar dentate nucleus - an exceptionally gyrified and convoluted structure - was initially represented by a dedicated anchor segmentation and was subsequently updated using the corresponding high-resolution FOCUS segmentation for the final registration framework. As expected, this preserved fine anatomical detail was not resolved to the same extent in the corresponding MNI reference, leading to a more envelope-like segmentation. The resulting lower Dice coefficient of 0.386 (MSD, 1.712 mm) should therefore be interpreted in the context of differences in boundary detail.

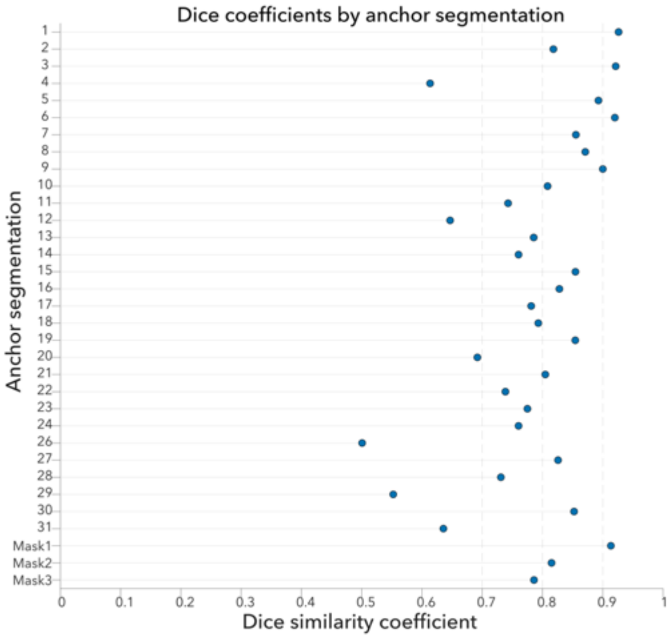

***Supplementary Figure 15.*** **Dice coefficients for individual anatomical anchor masks.** Dice coefficients were calculated between each segmentation defined in MNI space and its corresponding native-space segmentation after transformation into MNI space and reslicing to the common fixed T1-weighted reference image. Individual values characterize local anatomical correspondence across the regions contributing to the iterative registration. Dice coefficients for the three composite normalization masks are shown separately and summarize the overall fit of the aggregated anatomical constraints used for deformation-field optimization.

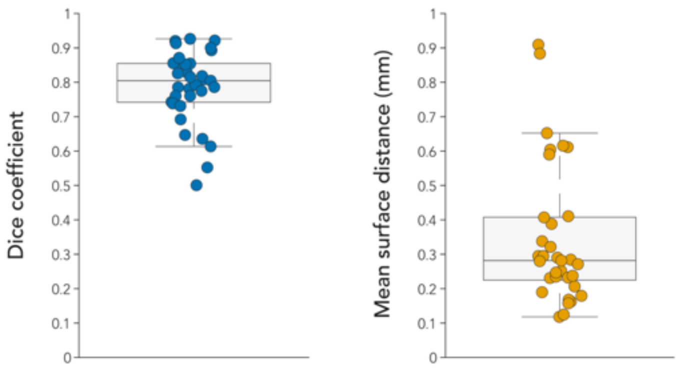

***Supplementary Figure 16*. Distribution of spatial agreement metrics across anatomical anchor masks.** Boxplots summarize the Dice coefficients and mean surface distances (MSD) obtained by comparing each MNI-Space anchor segmentation with its corresponding transformed native-space segmentation.

*WarpDrive Refinement*

In a small number of cases, residual local anatomical misalignments were corrected by limited manual refinement of the deformation field using WarpDrive.
To quantify the extent of manual intervention, we extracted the paired source and target fiducials stored by WarpDrive. Manual refinement comprised 17 individually placed point-to-point correspondences and one manually delineated anatomical outline, which was represented internally by 40 additional paired control points. Thus, 18 manual correction actions generated 57 source–target pairs. The magnitude of displacement across these pairs was small (mean, 0.99 mm; median, 1.06 mm; interquartile range, 0.74–1.20 mm; range, 0.15–2.54 mm), with only one pair exceeding 2 mm. All source-to-target pairs had identical z coordinates, indicating that the refinements were confined to the respective imaging planes.

***Supplementary Table 5.* Quantification of local manual refinement performed using WarpDrive**. *Please note:* Displacements are Euclidean distances between paired source and target fiducials in physical LPS coordinates. The manually delineated anatomical outline was represented internally by 40 paired control points. All source–target pairs had identical z coordinates; refinements were therefore confined to the respective imaging planes.

| Refinement type | Manual actions, n | Source–target pairs, n | Mean displacement, mm | Median displacement (IQR), mm | Range, mm |
|---|---|---|---|---|---|
| Point-to-point landmarks | 17 | 17 | 1.33 | 1.26 (1.08–1.62) | 0.51–2.54 |
| Anatomical outline | 1 | 40 | 0.85 | 0.91 (0.64–1.11) | 0.15–1.48 |
| **Overall** | **18** | **57** | **0.99** | **1.06 (0.74–1.20)** | **0.15–2.54** |

3. Cross-atlas variability of atlas definitions in MNI space

The preceding sections described two central objectives of the present atlas: first, to provide anatomically refined delineations of clinically relevant subcortical structures, and second, to transfer these definitions into MNI space with high spatial fidelity.

To illustrate why both aspects are critical, we compared existing atlas definitions already available in MNI space. Even for structures that are comparatively well resolved and readily identifiable on standard templates, such as the subthalamic nucleus (STN), pre-existing atlases differed markedly in spatial location, shape, extent, and anatomical boundary definition. These discrepancies indicate that the availability of an atlas definition in a common stereotactic space does not, by itself, ensure anatomical equivalence across resources.

We therefore visualized cross-atlas variability of STN definitions directly in MNI space and complemented this qualitative assessment with pairwise Dice and mean surface distance analyses. The visual comparisons (see *Supplementary Figure 17*) demonstrated substantial variation in the spatial configuration of STN labels across atlases ([7,42,44,46,47,173–179]). Across all unique pairwise comparisons among the 15 STN atlases, the mean Dice coefficient was 0.463 ± 0.186 and the mean surface distance was 0.929 ± 0.487 mm (mean ± SD), confirming broad dispersion in inter-atlas agreement (*Supplementary Figures 18 and 19*).

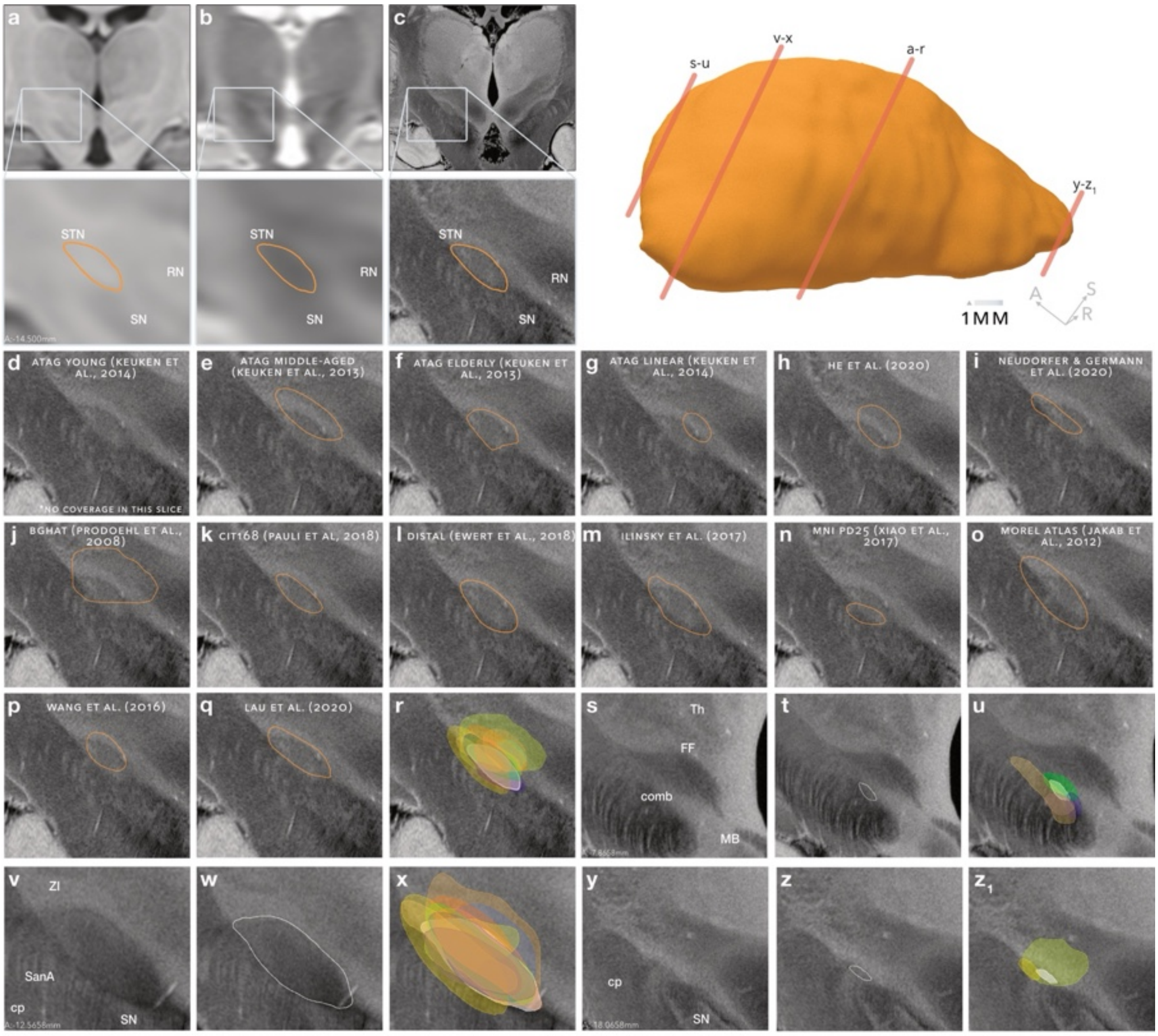

***Supplementary Figure 17.*** **Comparison of STN delineations across existing atlases in MNI-space.**
**a**–**c** Coronal sections through the subthalamic nucleus (STN) in the **a** MNI T1-weighted template, **b** MNI T2-weighted template, and **c** high-resolution 7-T MNI template, with corresponding magnified views of the STN. The selected section intersects a portion of the nucleus that is discernible across all three image contrasts. The three-dimensional rendering indicates the locations of the section planes shown in a–r, s–u, v–x, and y–z1. **d**–**q** STN delineations provided by existing MNI-space atlases, displayed on the corresponding high-resolution anatomical section. The **d** ATAG young-adult atlas does not cover the STN at this level, whereas the remaining atlases show variable agreement with the visible anatomical boundaries (**e**–**q**). **r** Overlay of all atlas delineations shown in d–q; the FOCUS STN delineation is shown in white. **s**–**u** Comparison at the anterior pole of the STN, where its boundary is difficult to distinguish because of the surrounding comb-fibre architecture. The coronal section is shown in **s**, the corresponding FOCUS delineation in **t**, and the combined delineations from the atlases shown in d–q in **u**. Most atlases incompletely or inaccurately capture this anterior extent. **v**–**x** Comparison at a section located anterior to the section shown in a–r, where the STN remains clearly discernible. The anatomical MR section is shown in **v**, the corresponding FOCUS delineation in **w**, and the overlay of all atlas delineations shown in d–q in **x**, with FOCUS shown in white as a reference. **y**–**z1** Comparison at the posterior pole of the STN, which is less readily visible on conventional MNI templates. The section is shown in **y**, the corresponding FOCUS delineation in **z**, and the overlay of existing atlas delineations in **z1**, with FOCUS again shown in white as a reference. Only a subset of atlases includes this posterior extent, with substantial variation in its localization and shape. These comparisons demonstrate marked discrepancies among existing atlases, even at levels where the nucleus is anatomically well visible in 0.5mm MNI space, underscoring the importance of both careful anatomical delineation and accurate registration to MNI space.

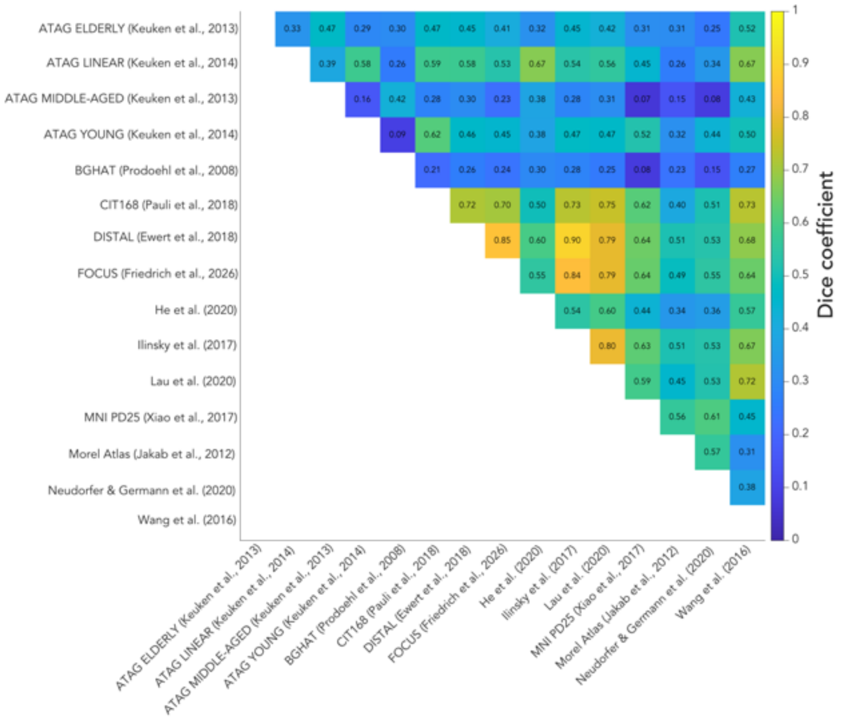

**Supplementary Figure 18. Pairwise dice coefficients of STN segmentations across atlases.** Across comparisons, the mean Dice coefficient was 0.463 ± 0.186 and the mean surface distance was 0.929 ± 0.487 mm (mean ± SD).

To position FOCUS within this existing atlas landscape, we distinguished between pairwise comparisons among established STN atlases, termed the non-FOCUS baseline, and comparisons between FOCUS and each established STN atlas, termed FOCUS vs non-FOCUS comparisons (*Supplementary Figure 19*). The non-FOCUS baseline showed a mean Dice coefficient of 0.447 ± 0.180 and a mean surface distance of 0.957 ± 0.482 mm. For FOCUS vs non-FOCUS comparisons, the corresponding values were 0.565 ± 0.198 and 0.744 ± 0.496 mm, respectively. Thus, FOCUS did not constitute a spatial outlier relative to existing STN definitions.

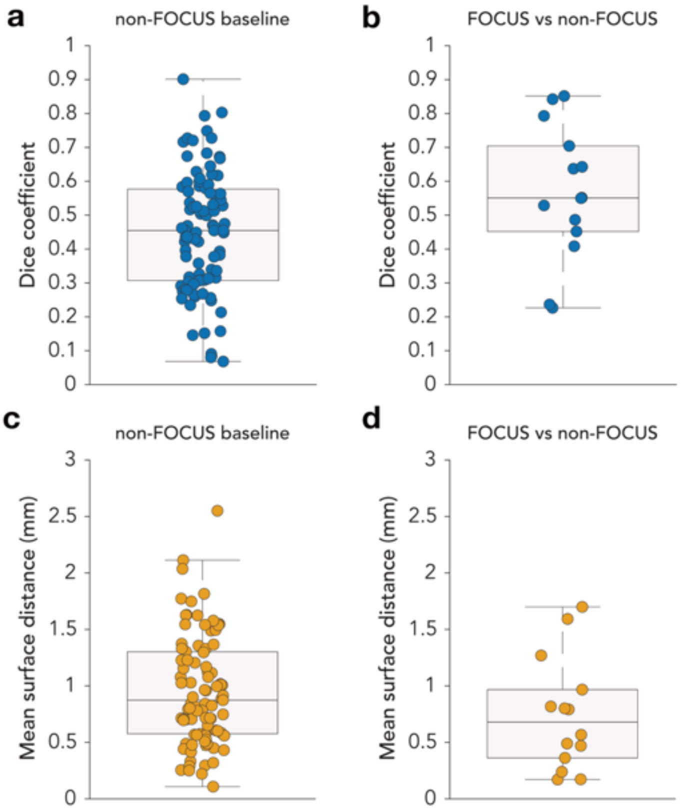

***Supplementary Figure 19.*** **Distribution of pairwise spatial agreement of the STN definition across MNI atlases.** Boxplots with individual pairwise comparisons showing Dice similarity coefficients **a**,**b** and mean surface distances **c**,**d**. **a**,**c** Pairwise comparisons among established MNI atlases, defined as the 'non-FOCUS baseline'. **b**,**d** Comparisons between FOCUS and each established MNI atlas. The 'non-FOCUS baseline' yielded a mean Dice coefficient of 0.447 ± 0.180 and a mean surface distance of 0.957 ± 0.482 mm, whereas 'FOCUS vs non-FOCUS' comparisons yielded 0.565 ± 0.198 and 0.744 ± 0.496 mm.

#### 4. Application of the FOCUS Atlas across healthy and pathological brain anatomy

To address the generalizability of the FOCUS atlas to diverse in-vivo anatomies, we performed an illustrative validation of the atlas across a heterogeneous cohort of openly available MRI scans. Using Lead-DBS[180] and WarpDrive [172], we registered the FOCUS atlas to each subject and examined the atlas fit in native space. The aim was to demonstrate how the same atlas definition behaves when applied to individual brains spanning different ages, diagnoses, and acquisition settings.

Ten subjects were randomly selected from a heterogeneous sample comprising healthy individuals and patients with neurological or psychiatric disorders to illustrate the applicability of the FOCUS atlas across diverse clinical populations (see *Supplementary Table 6* and *Supplementary Figure 20*). Structural scans were obtained from publicly available repositories. Public data included T1-weighted MRI from OpenNeuro datasets ds002790 (healthy young adults; [181]), ds000030 (controls and schizophrenia; [182]), ds004471 (Parkinson's disease;[183]), as well as one T1-weighted scan from OASIS-1 ([184]), and one healthy middle-aged adult from figshare (healthy adult;[185]). For each public dataset, the specific OpenNeuro or OASIS version used in this analysis is listed in *Supplementary Table 6* together with the corresponding DOI.

This cohort was intentionally heterogeneous with respect to age, sex, diagnosis, scanner vendor and field strength. The purpose of this heterogeneity was to demonstrate atlas behaviour under realistic clinical and research imaging conditions rather than to perform a balanced group comparison.

***Supplementary Table 6.*** **Cohort characteristics of structural MRI scans used for illustrative FOCUS atlas warping.** Ten heterogeneous T1-weighted scans from OpenNeuro, OASIS-1, and figshare, spanning healthy adulthood, schizophrenia, Parkinson disease, and dementia. Repository accessions and DOIs are provided.

| Subject ID | Age | Sex | Clinical Group | Repository | DOI |
|---|---|---|---|---|---|
| sub-001 | 35y | m | Healthy middle-aged adult | figshare 20134091 (Charité_01/2021_3T) | https://doi.org/10.6084/m9.figshare.20134091 |

| | | | | | |
|---|---|---|---|---|---|
| sub-0070 | 18.8y | m | Healthy young adult | OpenNeuro ds002790 | https://doi.org/10.18112/openneuro.ds002790.v2.0.0 |
| sub-0198 | 25y | m | Healthy young adult | OpenNeuro ds002790 | https://doi.org/10.18112/openneuro.ds002790.v2.0.0 |
| sub-10631 | 21y | f | Healthy young adult | OpenNeuro ds000030 | https://doi.org/10.18112/openneuro.ds000030.v1.0.0 |
| sub-50014 | 22y | m | Schizophrenia | OpenNeuro ds000030 | https://doi.org/10.18112/openneuro.ds000030.v1.0.0 |
| sub-50006 | 44y | f | Schizophrenia | OpenNeuro ds000030 | https://doi.org/10.18112/openneuro.ds000030.v1.0.0 |
| sub-10388 | 50y | f | Healthy middle-aged adult | OpenNeuro ds000030 | https://doi.org/10.18112/openneuro.ds000030.v1.0.0 |
| sub-10290 | 48y | m | Healthy middle-aged adult | OpenNeuro ds000030 | https://doi.org/10.18112/openneuro.ds000030.v1.0.0 |
| sub-oas28 | 86y | f | Dementia | OASIS-1 | https://doi.org/10.1162/jocn.2007.19.9.1498 |
| sub-106 | 63y | m | Parkinson disease | OpenNeuro ds004471 | https://doi.org/10.18112/openneuro.ds004471.v1.0.1 |

All subjects were processed using Lead-DBS v.3 [180]. For each subject, one primary structural volume was used as input (native T1-weighted MRI, or the MP2RAGE-derived UNIT1 image where applicable, T2 where applicable). Structural images were nonlinearly registered to MNI152NLin2009bAsym space[169] using the ANTs SyN diffeomorphic registration algorithm using the default parameters implemented in Lead-DBS[186]. In brief, this follows a standard five-stage preset with subcortical refinement masks [187]. No manual AC-PC correction, CT-MRI coregistration, brain-shift correction, or dataset-specific parameter tuning was performed. After visual quality control (QC) of the native-space atlas overlay, WarpDrive was used for optional fine adjustment of the deformation field when residual local misalignment was judged to compromise atlas fit.

Atlas labels defined in MNI152NLin2009bAsym template space were then transformed into each subject's preprocessed native T1 space via the inverse normalization field estimated above. These patient-specific native-space segmentations constituted the data for subsequent quantitative assessment of atlas fit.

For qualitative display, atlas overlays were rendered in native space using the patient-specific warped FOCUS atlas. The preprocessed native structural image served as the background volume, with the corresponding native-space atlas labels overlaid as live slice contours in 3D-Slicer[188] via the Slicer Netstim WarpDrive/ImportAtlas tools (https://github.com/netstim/SlicerNetstim )[189].

Quantitative descriptors of the atlas fit were computed from the patient-specific warped segmentations in native space. For each subject and structure of interest, binary labels derived from the Lead-DBS native-space warp were used to calculate descriptive metrics including structure volume, centroid coordinates, and inter-structure distances (e.g., STN-GPi separation). This approach quantifies individual atlas placement and morphology after the warping pipeline. Metrics were intended to summarize relative variability across subjects under a uniform workflow.

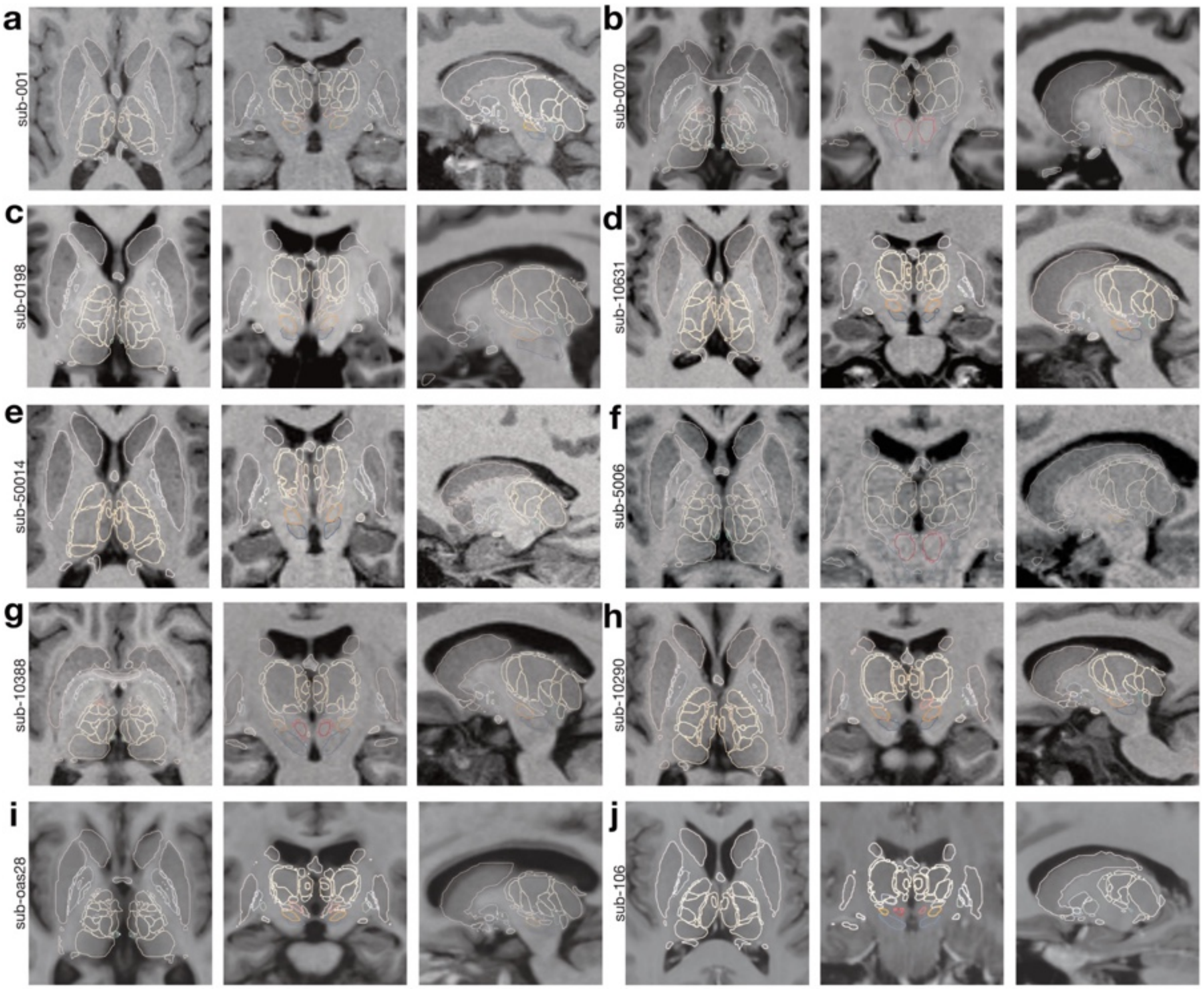

***Supplementary Figure 20.*** **Atlas fit in native space shown in axial, coronal, and sagittal views for each subject.** Note that some subjects, such as sub-10388 or sub-oas28 show ventricular enlargement as examples of ageing- and disease-related anatomical alterations.

Descriptive atlas fit metrics were computed from patient-specific FOCUS segmentations warped into native space (voxel threshold > 0.5). Across the cohort, bilateral STN volumes were 285.3 ± 46.0 mm³ (coefficient of variation (CV) = 16.1%; range 226.9-375.2 mm³). GPi and GPe volumes showed similar relative dispersion (CV = 12.4% and 10.6%, respectively). The within-subject STN-GPi centroid distances were 8.32 ± 0.42 mm (CV = 5.0%; range 7.42–9.02 mm), indicating modest inter-individual variability in the spatial relationship between these structures after application of the same patient-specific registration workflow. The GPe/GPi volume ratios were highly stable (2.31 ± 0.06; CV = 2.7%). Left-right STN volume asymmetries averaged 5.2 ± 2.2%.
Registration quality, assessed by z-scored normalized (spatial) cross-correlations between each subject's MNI-normalized T1 and the MNI152NLin2009bAsym template, ranged from -0.20 to 0.57 (median 0.45). Lower values occurred in an elderly patient with dementia (sub-oas28; NCC = 0.10) and in one with advanced Parkinson's Disease (sub-106; NCC = -0.20), likely reflecting pathology-related anatomical alterations rather than necessarily indicating poorer atlas fit, although residual registration error cannot be fully excluded.

***Supplementary Table 7.*** **Patient-specific atlas fit metrics derived from native-space FOCUS warping.** Descriptive metrics for each of the 10 figure-cohort subjects after identical Lead-DBS preprocessing, ANTs normalization, and patient-specific FOCUS atlas warping (ptnative = true). Bilateral structure volumes (mm³) and the STN-GPi distance (mm) were computed from thresholded (> 0.5) warped segmentations in each subject's native space. STN-GPi dis-tance denotes the Euclidean separation between bilateral-averaged structure centroids within the same subject. The GPe/GPi ratio summarizes relative pallidal compartment vol-umes. STN L/R asymmetry is defined as |V_left − V_right| / (V_left + V_right) × 100. MNI normalization NCC is the z-scored intensity correlation between the subject's normalized T1-weighted image and the MNI152NLin2009bAsym template.

| Subject ID | Age | Sex | Clinical Group | STN volume (mm³) | GPi volume (mm³) | GPe volume (mm³) | STN-GPi distance (mm) | GPe/GPi ratio | STN L/R asymmetry (%) | MNI normalization NCC |
|---|---|---|---|---|---|---|---|---|---|---|

| | | | | | | | | | | |
|---|---|---|---|---|---|---|---|---|---|---|
| sub-001 | 35y | m | Healthy middle-aged adult | 265.4 | 647.6 | 1545.8 | 8.42 | 2.39 | 7.4 | 0.36 |
| sub-0070 | 18.8y | m | Healthy young adult | 319.3 | 723.8 | 1683.8 | 8.32 | 2.33 | 3.5 | 0.56 |
| sub-0198 | 25y | m | Healthy young adult | 375.2 | 816.8 | 1828.1 | 9.02 | 2.24 | 3.4 | 0.57 |
| sub-10631 | 21y | f | Healthy young adult | 278.7 | 684.4 | 1537.5 | 8.42 | 2.25 | 2.7 | 0.51 |
| sub-50014 | 22y | m | Schizo-phrenia | 340.6 | 800.8 | 1792.4 | 8.17 | 2.24 | 6.9 | 0.44 |
| sub-50006 | 44y | f | Schizo-phrenia | 266.5 | 638.9 | 1515.3 | 7.42 | 2.37 | 8.2 | 0.35 |
| sub-10388 | 50y | f | Healthy middle-aged adult | 239.1 | 614.3 | 1392.8 | 8.21 | 2.27 | 2.7 | 0.48 |
| sub-10290 | 48y | m | Healthy middle-aged adult | 269.8 | 637.8 | 1475.6 | 8.77 | 2.31 | 4.1 | 0.46 |
| sub-oas28 | 86y | f | Dementia | 226.9 | 541.0 | 1297.9 | 8.23 | 2.40 | 7.6 | 0.10 |
| sub-106 | 63y | m | Parkinson disease | 271.6 | 669.5 | 1571.8 | 8.26 | 2.35 | 5.5 | −0.20 |

***Supplementary Table 8.*** **Cohort-level variability in atlas-derived metrics (n = 10).** Summary statistics for key atlas fit descriptors across the same cohort shown in *Supplementary Table 6*. Values were derived from patient-specific native-space FOCUS warps under a uniform registration pipeline. CV denotes the coefficient of variation [(SD / mean) × 100]. Lower CV for inter-structure distances and volume ratios indicates greater consistency in relative subcortical geometry across subjects; higher CV for absolute structure volumes reflects combined anatomical heterogeneity, acquisition differences, and registration variability. These statistics characterize interindividual variability in patient-specific atlas placement and do not represent validation against histological or manual reference standards.

| Metric | n | Mean | SD | CV (%) | Median | Range |
|---|---|---|---|---|---|---|
| STN volume ($mm^3$) | 10 | 285.3 | 46.0 | 16.1 | 270.7 | 226.9–375.2 |
| GPi volume ($mm^3$) | 10 | 677.5 | 84.0 | 12.4 | 658.5 | 541.0–816.8 |
| GPe volume ($mm^3$) | 10 | 1564.1 | 165.9 | 10.6 | 1541.7 | 1297.9–1828.1 |
| RN volume ($mm^3$) | 10 | 500.3 | 75.6 | 15.1 | 477.8 | 401.1–627.8 |
| STN-GPi distance (mm) | 10 | 8.32 | 0.42 | 5.0 | 8.29 | 7.42–9.02 |
| GPe/GPi volume ratio | 10 | 2.31 | 0.06 | 2.7 | 2.32 | 2.24–2.40 |
| STN L/R volume asymmetry (%) | 10 | 5.2 | 2.2 | 41.8 | 4.8 | 2.7–8.2 |
| MNI normalization NCC | 10 | 0.36 | 0.24 | 66.1 | 0.45 | −0.20–0.57 |

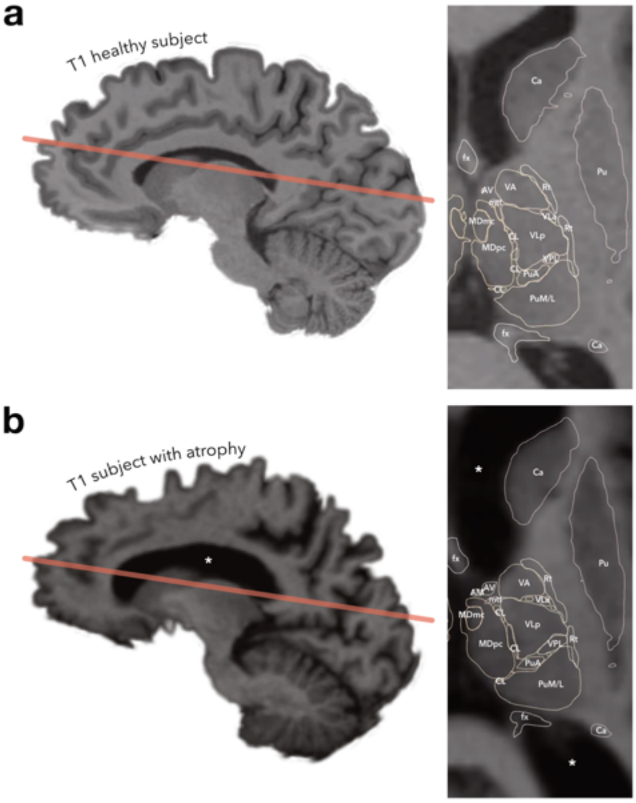

***Supplementary Figure 21.* Close-up demonstration of atlas fit in a healthy adult and in a patient with dementia-associated brain atrophy**. **a** T1-weighted MRI of a healthy adult (sub-001). **b** T1-weighted MRI of a patient with dementia-associated brain atrophy (sub-oas28), characterized by pronounced ventricular enlargement (white asterisks).

Two complementary patterns emerged from these analyses. First, relative subcortical geometry was substantially more stable than absolute structure size: the STN-GPi centroid distance varied by only ~1.6 mm across the cohort (CV ≈ 5%), and the GPe/GPi volume ratio remained nearly constant (CV ≈ 3%), whereas absolute STN and pallidal volumes showed greater dispersion (CV ≈ 11–16%). This dissociation suggests that, under a uniform registration workflow, the FOCUS atlas preserves expected topological relationships among neighbouring deep nuclei even when overall structure volumes differ across individuals, scanners, and clinical contexts.

Second, the largest deviations in registration QC and in absolute volumes clustered in predictable edge cases, most notably advanced age with atrophy (sub-oas28) and/or Parkinson's Disease (sub-106) – both well known to affect normalization accuracy ([172]) – rather than being distributed uniformly across the cohort. Close-up native-space inspection can be particularly important in subjects with pronounced brain atrophy, as illustrated by subject sub-oas28 (see *Supplementary Figure 21b*). In this setting, disease-related morphological changes and ventricular enlargement may lead to local mismatches in the initial atlas fit, underscoring the importance of careful visual quality control. As demonstrated in sub-oas28, these local discrepancies can be effectively corrected through WarpDrive refinement without altering the overall atlas definition.

The quantitative metrics should be read as descriptors of inter-individual variability after application of the same patient-specific registration workflow, not as proof of histological accuracy. Absolute volumes are sensitive to segmentation threshold, partial-volume effects, and residual registration error; distances and volume ratios are comparatively robust summaries of relative atlas geometry. The MNI normalization NCC likewise provides a coarse whole-brain registration QC signal and is not a structure-wise ground-truth metric for deep nuclei.

In this illustrative in vivo evaluation, the FOCUS atlas could be deployed through a standardized Lead-DBS workflow across a deliberately heterogeneous adult cohort spanning healthy controls, schizophrenia, Parkinson's disease, and dementia. After patient-specific warping into native space using the standardized patient-specific registration workflow, relative subcortical geometry remained more consistent than absolute structure size, supporting practical generalizability of the atlas definition beyond the single post-mortem template from which it was derived. At the same time, geriatric and clinical acquisitions showed the greatest registration QC deviations and benefited most from careful visual review, with the WarpDrive tool available for fine adjustment when needed.

69. Meynert, T. (1833-1892). A. du texte (1884). Psychiatrie : Klinik der Erkrankungen des Vorderhirns, begründet auf dessen Bau, Leistungen und Ernährung / von Dr Theodor Meinert,... (W. Braumüller (Wien)).

70. MEYNERT T. (1872). Vom Gehirne der Saugethiere. Handb. Lehre Von Den Geweben Menschen Thiere *2*, 694–808.

71. Gheban, B.-A., Colosi, H.A., Gheban-Rosca, I.-A., Pop, B., Domșa, A.-M.T., Georgiu, C., Gheban, D., Crișan, D., and Crișan, M. (2021). Age-Related Changes of the Pineal Gland in Humans: A Digital Anatomo-Histological Morphometric Study on Autopsy Cases with Comparison to Predigital-Era Studies. Medicina (Mex.) *57*, 383. https://doi.org/10.3390/medicina57040383.

72. Harisha, K., Arava, S., Singh, S., and Pariplavi, M. (2023). A morphological study of human pineal gland in post-mortem cases. Morphologie *107*, 193–198. https://doi.org/10.1016/j.morpho.2022.06.095.

73. Díaz, E., Bravo, D., Rojas, X., and Concha, M.L. (2011). Morphologic and immunohistochemical organization of the human habenular complex. J. Comp. Neurol. *519*, 3727–3747. https://doi.org/10.1002/cne.22687.

74. Müller, U.J., Ahrens, M., Vasilevska, V., Dobrowolny, H., Schiltz, K., Schlaaff, K., Mawrin, C., Frodl, T., Bogerts, B., Gos, T., et al. (2021). Reduced habenular volumes and neuron numbers in male heroin

addicts: a post-mortem study. Eur. Arch. Psychiatry Clin. Neurosci. *271*, 835–845. https://doi.org/10.1007/s00406-020-01195-y.

75. An, X., Bandler, R., Öngür, D., and Price, J.L. (1998). Prefrontal cortical projections to longitudinal columns in the midbrain periaqueductal gray in Macaque monkeys. J. Comp. Neurol. 1911 *401*, 455–479.

76. Büttner-Ennever, J.A., and Büttner, U. (1988). Neuroanatomy of the oculomotor system. The reticular formation. Rev. Oculomot. Res. *2*, 119.
